# Exploration of the Tunability of BRD4 Degradation by DCAF16 *Trans*-labelling Covalent Glues

**DOI:** 10.1101/2023.10.07.561308

**Authors:** Muhammad Murtaza Hassan, Yen-Der Li, Michelle W. Ma, Mingxing Teng, Woong Sub Byun, Kedar Puvar, Ryan Lumpkin, Brittany Sandoval, Justine C. Rutter, Cyrus Y. Jin, Michelle Y. Wang, Shawn Xu, Anna M. Schmoker, Hakyung Cheong, Brian J. Groendyke, Jun Qi, Eric S. Fischer, Benjamin L. Ebert, Nathanael S. Gray

**Affiliations:** Department of Chemical and Systems Biology, ChEM-H and Stanford Cancer Institute, Stanford School of Medicine, Stanford University, Stanford, CA; Department of Molecular and Cellular Biology, Harvard University, Cambridge, MA; Department of Medical Oncology, Dana-Farber Cancer Institute, Boston, MA; Cancer Program, Broad Institute of MIT and Harvard, Cambridge, MA; Department of Cancer Biology, Dana-Farber Cancer Institute, Boston, MA; Department of Biological Chemistry and Molecular Pharmacology, Harvard Medical School, Boston, MA; Center for Drug Discovery, Department of Pathology & Immunology, and Verna and Marrs McLean Department of Biochemistry and Molecular Pharmacology, Baylor College of Medicine, Houston, TX; Howard Hughes Medical Institute, Boston, MA

## Abstract

Small molecules that can induce protein degradation by inducing proximity between a desired target and an E3 ligase have the potential to greatly expand the number of proteins that can be manipulated pharmacologically. Current strategies for targeted protein degradation are mostly limited in their target scope to proteins with preexisting ligands. Alternate modalities such as molecular glues, as exemplified by the glutarimide class of ligands for the CUL4^CRBN^ ligase, have been mostly discovered serendipitously. We recently reported a *trans*-labelling covalent glue mechanism which we named ‘Template-assisted covalent modification’, where an electrophile decorated small molecule binder of BRD4 was effectively delivered to a cysteine residue on an E3 ligase DCAF16 as a consequence of a BRD4-DCAF16 protein-protein interaction. Herein, we report our medicinal chemistry efforts to evaluate how various electrophilic modifications to the BRD4 binder, JQ1, affect DCAF16 *trans*-labeling and subsequent BRD4 degradation efficiency. We discovered a decent correlation between the ability of the electrophilic small molecule to induce ternary complex formation between BRD4 and DCAF16 with its ability to induce BRD4 degradation. Moreover, we show that a more solvent-exposed warhead presentation is optimal for DCAF16 recruitment and subsequent BRD4 degradation. Unlike the sensitivity of CUL4^CRBN^ glue degraders to chemical modifications, the diversity of covalent attachments in this class of BRD4 glue degraders suggests a high tolerance and tunability for the BRD4-DCAF16 interaction. This offers a potential new avenue for a rational design of covalent glue degraders by introducing covalent warheads to known binders.

## Introduction

Chemically induced protein degradation has attracted significant interest owing to its unique pharmacology and modular design strategy.^1–5^ Unlike the traditional ‘occupancy-driven’ pharmacology, targeted protein degradation (TPD) offers a catalytic ’event-driven’ pharmacology. Furthermore, degraders have shown to have a potential target scope beyond protein targets that are typically perceived as druggable.^5,6^ Traditional inhibitors require deep lipophilic protein pockets suitable for ligand-binding, which hinders inhibitor development for protein targets containing smooth, shallow, and featureless surfaces lacking ligand-protein complementarity.^7–9^ Degraders, however, are capable of harnessing protein-protein surface complementarity, which expands the target scope to proteins previously considered undruggable, such as transcription factors.^10–13^

TPD strategies which exploit the ubiquitin-dependent proteolysis pathway are commonly classified into two main chemical degrader categories: Proteolysis Targeting Chimeras (PROTACs) and molecular glue degraders (MGDs).^14,15^ PROTACs are heterobifunctional molecules that contain an E3 ligase binder tethered via a linker to a protein-of-interest (POI) binder. Although PROTACs offer a modular design strategy, their target scope is limited to POIs that are ligandable and thereby precludes access to ‘undruggable’ targets. Unlike PROTACs, molecular glues are monovalent molecules that can alter a protein’s interactome to induce protein-protein interactions with new protein-partners. Since MGD chemical design is not plug-and-play by nature, their discovery has largely been serendipitous.

Recently we reported MGD that use minimalistic covalent handles. Previous studies had reported the degradation of BRD4 by GNE011, an analog of the BRD4 selective inhibitor JQ1 (Figure 1A).^16^ Studies by our group and others showed that GNE011 induces the degradation of BRD4 via recruitment of the DDB1 and CUL4 associated factor 16 (DCAF16) E3 ligase to bromodomain 2 (BRD4_BD2_).^17–19^ Via a series of electrophilic substitutions of the propargyl amine tail of GNE011, we discovered that covalent warhead attachment to JQ1 can lead to improved BRD4 degradation compared to GNE011. Further mechanistic and structural analyses uncovered a novel mechanism of action which we termed ‘template-assisted covalent modification’ where BRD4 degradation required the covalent *trans*-labelling of Cys58 on DCAF16. A cryo-electron microscopy (cryo-EM) structure of BRD4_BD2_-MMH2-DCAF16-DDB1 showed structural complementarity between BRD4_BD2_ and DCAF16, and suggested that upon binding to covalent JQ1 analogs, BRD_BD2_ serves as a structural template to facilitate DCAF16 covalent modification (Figure 1C).^18^

**Figure 1.**
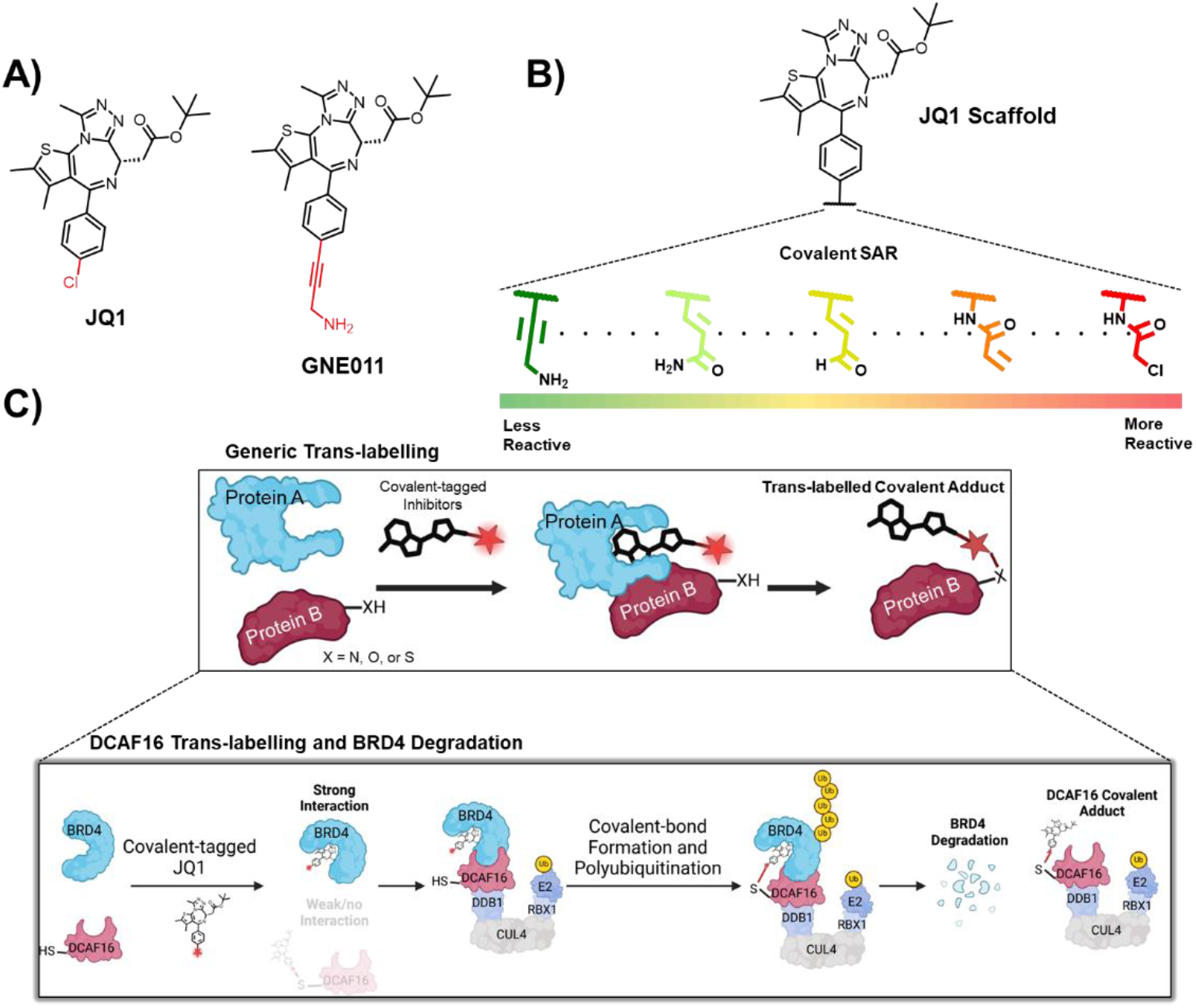
A) The chemical structures of the BRD4 inhibitor JQ1 and its related analog GNE011. B) Investigating various cysteine reactive electrophilic handles. C) A schematic depiction of a general covalent trans-labelling event and its relevance to our previous study showing the degradation of BRD4 by covalently trans-labelling DCAF16.

Herein, we describe the medicinal chemistry campaign that led to the discovery of the covalent JQ1 analogs and explore the relationship between BRD4 degradation with covalent-warhead substitution (Figure 1B).

## Results

### Covalent Analogs show tunability of DCAF16-dependent BRD4_BD2_ degradation

In order to investigate the degradation of BRD4 by GNE011 and related analogs, we used a fluorescent stability reporter assay reported previously (Figure 2A).^16^ This assay involves the co-expression of mCherry with BRD4_BD1/2_ fused to an enhanced green fluorescent protein (eGFP) tag, and the subsequent fluorescence measurement of BRD4_BD1/2_-eGFP normalized to mCherry as a readout for BRD4_BD1/2_ degradation. It allows for a quantitative, rapid, and facile screening of several MGDs, in a dose-and time-dependent manner. Hence, this assay was first used to test GNE011 with a 16 h incubation in K562 cells. In this assay, GNE011 showed selective degradation of the second bromodomain (BRD4_BD2_) over the first (BRD4_BD1_) with ∼50% maximal degradation (D_max/16h_) at 16 h (Figure 2C).^18^ GNE011 was confirmed by us and others, to be degraded via recognition by DCAF16.^17–19^ Thus, the BRD4_BD2_ stability system was used to test all subsequent GNE011 analogs in wild type and *DCAF16-/-* cells.

**Figure 2.**
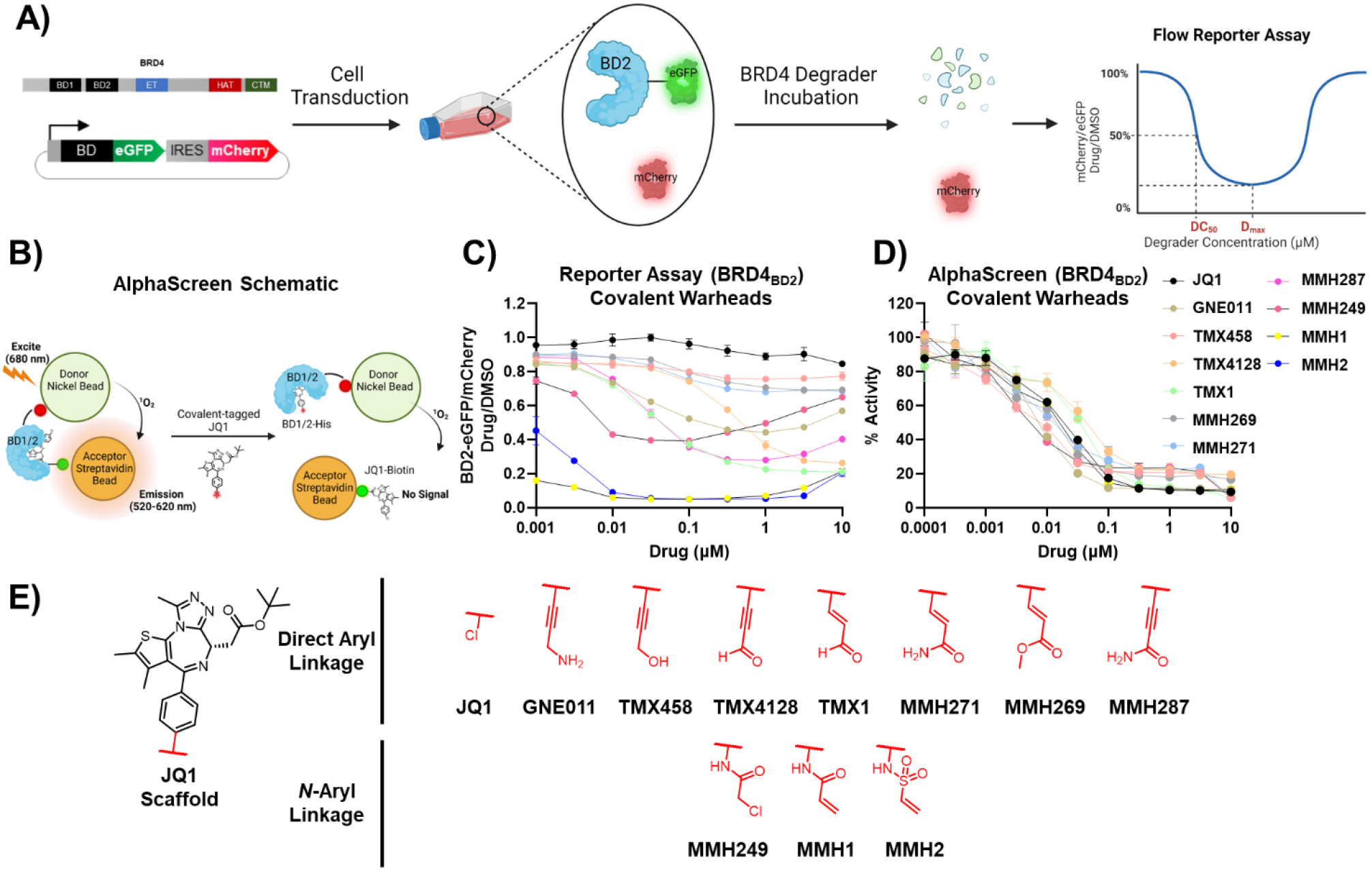
A) A schematic depiction of the BRD4_BD2_-eGFP and mCherry reporter assay and B) an AlphaScreen assay in the context of BRD4 degraders. C) Dose-dependent BRD4_BD2_ degradation by covalent JQ1 analogs as determined by the BRD4_BD2_-eGFP and mCherry reporter assay. D) The dose-dependent inhibition of BRD4_BD2_ by covalent JQ1 analogs as determined by an AlphaScreen assay. E) The chemical structures of the covalent JQ1 degradation tails included in the covalent SAR. Non-covalent analogs are not included. Note: MMH287, and MMH2 were not tested for AlphaScreen BRD4_BD2_.

Next, to understand the role of the propargyl amine group for GNE011 induced BRD4 degradation, we introduced propargyl alcohol (TMX458) and propargyl aldehyde (TMX4128) to JQ1. TMX458 and TMX4128 showed disparate reactivities, with TMX458 being completely inactive (D_max/16h_ = 10%, DC_50_ > 10 µM) and TMX4128 showing enhanced BRD4 degradation in comparison to GNE011 (D_max/16h_ = 69%, DC_50_ = 703 nM). Given the literature precedence of propargyl aldehydes and alkynes as covalent warhead targeting thiols, we suspected the degradation might be linked to the covalency of GNE011 and its analogs.^20–22^ Furthermore, drugs with propargyl amine handles such as rasagiline and selegiline are reported to irreversibly inhibit monoamine oxidase enzymes.^23–26^ Hence, we focused our subsequent structural activity relationship (SAR) studies to survey various covalent warhead attachments (Figure 1b).

Firstly, we designed a series of Michael acceptor chemotypes, including acrolein (TMX1), methyl acrylate (MMH269), aryl acrylamide (MMH271), and propiolamide (MMH287). MMH269 and MMH271 showed almost no DCAF16 dependent BRD4_BD2_ degradation, whereas TMX1 and MMH287 were essentially equipotent with respect to their D_max/16h_ (75% and 69%) and DC_50_ values (64 nM and 51 nM; figure S1).

Next, we wanted to confirm whether covalent bond formation was a requirement for DCAF16 dependent BRD4_BD2_ degradation. Intact mass spectrometry and mutagenesis experiments validated the mass adduct formation of TMX1-DCAF16 upon co-incubation with BRD4_BD2_ and identified DCAF16 Cys58 as the targeted residue. Moreover, mutagenesis studies using a CRISPR alanine screen (C58A), and TR-FRET experiments (C58S) showed a complete rescue of BRD4 degradation and DCAF16 recruitment respectively confirming the necessity for DCAF16 covalent-bond formation for BRD4 degradation.^18^

We hypothesized that a more solvent exposed terminal covalent analog could yield more potent BRD4 degraders and allow for a more optimal DCAF16 recruitment. Inspired by the *N*-aryl attachment handles on JQ1 used by Dragovich *et al*. for Genentech’s antibody-PROTAC conjugate, we sought to create *N*-aryl terminally solvent exposed covalent JQ1 analogs. We synthesized a series of terminally solvent exposed covalent analogs with an *N*-aryl linkage. The chloroacetamide (MMH249), acrylamide (MMH1), and vinyl sulfonamide (MMH2) all demonstrated a significant improvement in degradation potency. Additionally, their activity was almost entirely ablated in DCAF16 knockout (KO) K562 cells (figure S1). MMH249 showed a DC_50_ of 8 nM but failed to completely degrade BRD4_BD2_ and reached a D_max/16h_ of 56%. MMH1 and MMH2, however, demonstrated nearly quantitative BRD4_BD2_ degradation (95%), and DC_50_ values of 0.3 nM and 1 nM respectively. These compounds displayed a ∼3333- and 1000-fold increase in degradation potency in comparison to GNE011.

### Differential BRD4_BD2_ degradation potency does not correlate with a differential BRD4_BD2_ engagement

To test whether differential degradation potencies could be explained by differences in BRD4_BD2_ affinity, we measured the *in vitro* binding potencies of the covalent library against BRD4_BD1/2_ using a competition-based luminescent Amplified Luminescent Proximity Homogenous Assay (AlphaScreen). This assay utilizes donor and acceptor beads, each conjugated to a (bio)molecule of interest. Upon excitation by light, donor beads generate singlet oxygen species (^1^O_2_), that can get transferred to the acceptor beads given sufficient donor-acceptor proximity, which then emit luminescence. In this case, His tagged BRD4_BD1/2_ and biotin-tagged JQ1 (JQ1-biotin) were used in conjunction with Nickel-acceptor and Streptavidin-donor beads, respectively (Figure 2B).^27^ The competitive displacement of JQ1-biotin by the covalent analogs was measured to determine BRD4_BD1/2_ binding potencies.

All covalent analogs tested demonstrated comparable nanomolar BRD4_BD1_ and BRD4_BD2_ binding potencies (IC_50_ values ∼1 to 49 nM and 2 to 33 nM respectively; table S1) suggesting that the differential BRD4_BD2_ degradation potencies of the covalent analogs were not due to differences in affinity for BRD4 (Figure 2D, S6A). Likewise, all non-covalent analogs despite showing nanomolar BRD4_BD1/2_ engagement showed no significant DCAF16-depdendent BRD4_BD2_ degradation (figure S2, S6B, C), further suggesting that the DCAF16 covalent interaction and DCAF16-induced BRD4_BD2_ degradation have a causal relationship that is tunable by electrophilic warhead substitution.

### Characterization of BRD4_BD2_-DCAF16 Ternary Complex Formation Induced by JQ1 Analogs

To determine the relationship between covalent warheads and DCAF16 recruitment to BRD4_BD2_, we used a time-resolved fluorescence energy transfer (TR-FRET) assay to measure induced BRD4_BD2_-DCAF16 ternary complex formation (Figure 3A). This assay measures fluorescence transfer from a donor fluorophore-conjugated biomolecule to an acceptor fluorophore-conjugated biomolecule, which can be facilitated by chemically induced proximity between the two biomolecules (i.e., by PROTACs or MGDs).^28^ Using biotinylated BRD4_BD2_ with terbium-tagged streptavidin (donor), and BODIPY-FL labelled DCAF16 (acceptor), TR-FRET ratio (520 nm/490 nm) measurements were made after a 6 h degrader incubation time (Figure 3B).

**Figure 3.**
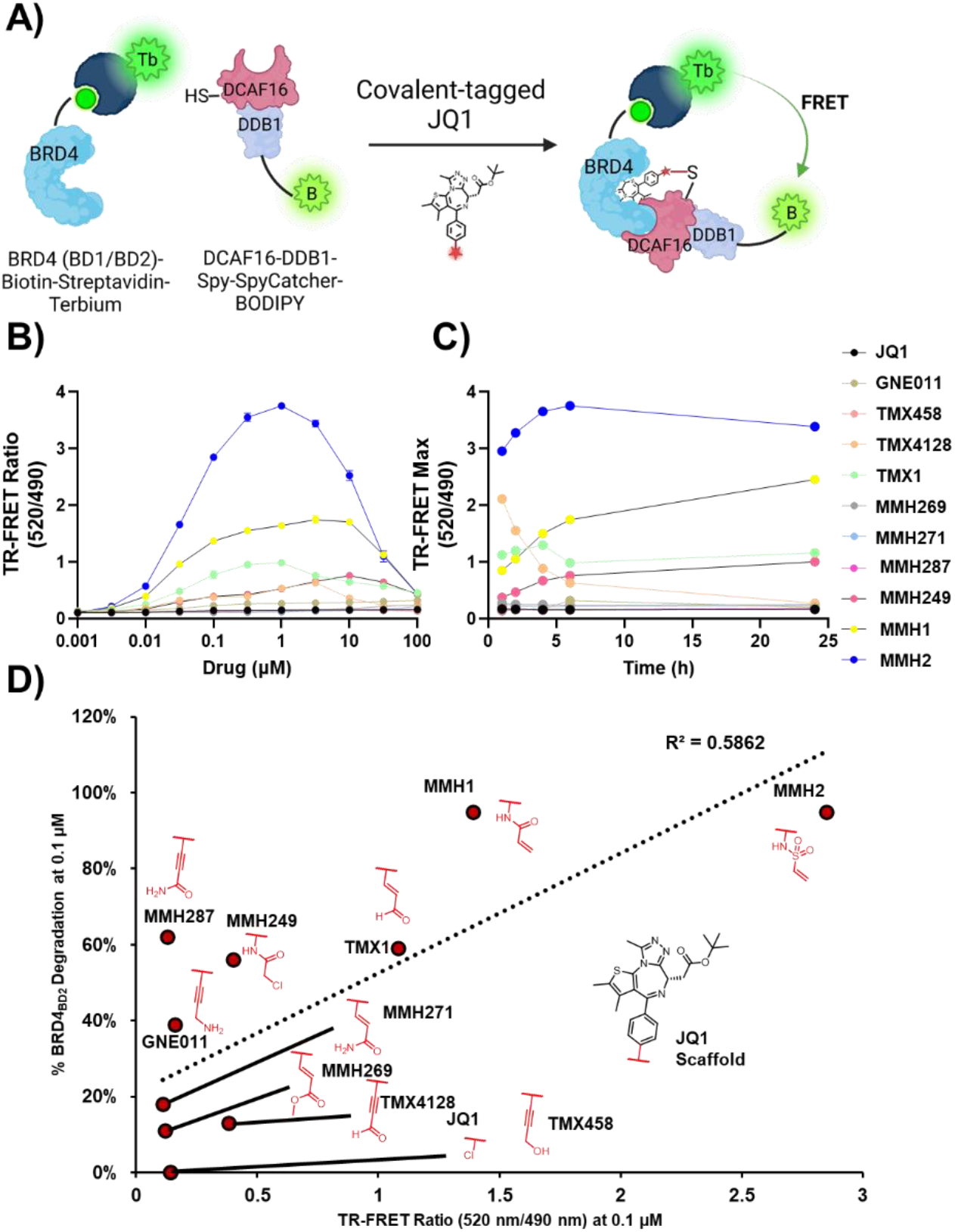
A) A schematic diagram of the BRD4_BD2_-DCAF16 TR-FRET assay. B) A 6 h TR-FRET assay for all covalent JQ1 analogs (520 nm/490 nm). C) A graph of TR-FRET maximum ratio vs. time for all covalent JQ1 analogs at 1, 2, 4, 6, and 24 h incubation times. D) A correlational analysis between the TR-FRET Ratio (520 nm/490 nm) and the % BRD4_BD2_ degradation for all JQ1 analogs.

Furthermore, we wanted to determine the kinetics of the BRD4_BD2_-DCAF16 interaction given its covalent nature. Hence, we conducted a time-dependent TR-FRET analysis where we measured the TR-FRET ratio after incubation for 1, 2, 4, 6, and 24 h (Figure 3C, S4, S5). Additionally, to understand the relationship between ternary complex formation kinetics and warhead reactivity we measured the depletion of the covalent analogs in the presence of glutathione using MS (Table S2). MMH1 and MMH249 which showed modest half-lives (*t*_1/2_ = 572 and 531 min) showed a progressive ternary complex formation over time and showed the maximum TR-FRET signal at 24 h, whereas TMX1 and MMH2 (*t*_1/2_ = 415 and > 750 min) showed saturation at 1 h. There did not appear to be a direct correlation between warhead reactivity and time-dependent saturation. For instance, MMH287 and TMX1, the two fastest reacting analogs (*t*_1/2_ = 380 min and 415 min), showed completely disparate ternary complex formation kinetics where unlike TMX1 (saturation at 1h) MMH287 showed almost no increase in ternary complex formation (figure S4). Peculiarly, TMX4128, a reactive warhead (*t*_1/2_ = 530 min) showed maximum TR-FRET signal at 1 h but showed a disintegration of the ternary complex overtime (TR-FRET_max_ = 2.12 and 0.28 at 1 and 24 h respectively).

Expectedly, non-degradative covalent analogs such as TMX458, MMH269, and MMH271, as well as the non-covalent analogs showed no significant DCAF16 recruitment (figure S4, S5).

We hypothesized that the BRD4_BD2_ degradation potency of the covalent analogs would correlate with ternary complex formation (Figure 3D). As expected, potent degraders such as TMX1, MMH1, and MMH2 showed a strong BRD4_BD2_- DCAF16 ternary complex formation with TR-FRET_max_ of 1.34, 1.78, and 3.75 respectively. However, other modest degraders such as TMX4128, MMH249, MMH287, and GNE011 demonstrated TR-FRET_max_ of 0.65, 0.77, 0.15, and 0.18, with only TMX4128 and MMH249 showing significant ternary complex formation. Non-or poorly-degrading covalent analogs such as TMX458, MMH269, and MMH271 showed minimal or no-ternary complex formation (TR-FRET_max_ = 0.16, 0.18, 0.14) on par with JQ1 (TR-FRET_max_ = 0.15; figure S3). To determine whether there was a linear correlation between BRD4_BD2_ degradation potency and ternary complex formation, we conducted a correlational analysis where the TR-FRET ratio was plotted against the % degradation at 0.1 µM concentration (Figure 3D). The graph showed an R^2^ value of 0.59, indicating a modest linear trend. GNE011 and MMH287 might possibly show a disparity between their *in vitro* TR-FRET ratios and cellular BRD4 degradation since the cellular activity could be driven by a chemically reactive metabolite of the alkyne chemotype.^23–26^

### Determining cellular DCAF16-dependent BRD4 degradation potencies

Western blotting was used to determine the degradation potencies of the covalent BRD4 degraders. Both wild-type and DCAF16 KO K562 cells were treated with the covalent JQ1 library at 1 µM concentrations and a 16 h incubation time (Figure 4A). All moderate to strong degraders such as TMX1, TMX4128, MMH1, MMH2, and MMH287 showed nearly complete BRD4 degradation that was completely rescued by DCAF16 KO. Similar to the BRD4_BD2_ reporter assay results for GNE011 and MMH249 which showed a 49 and 56% D_max_, both degraders displayed moderate degradation by western blot (Figure 4A).

**Figure 4.**
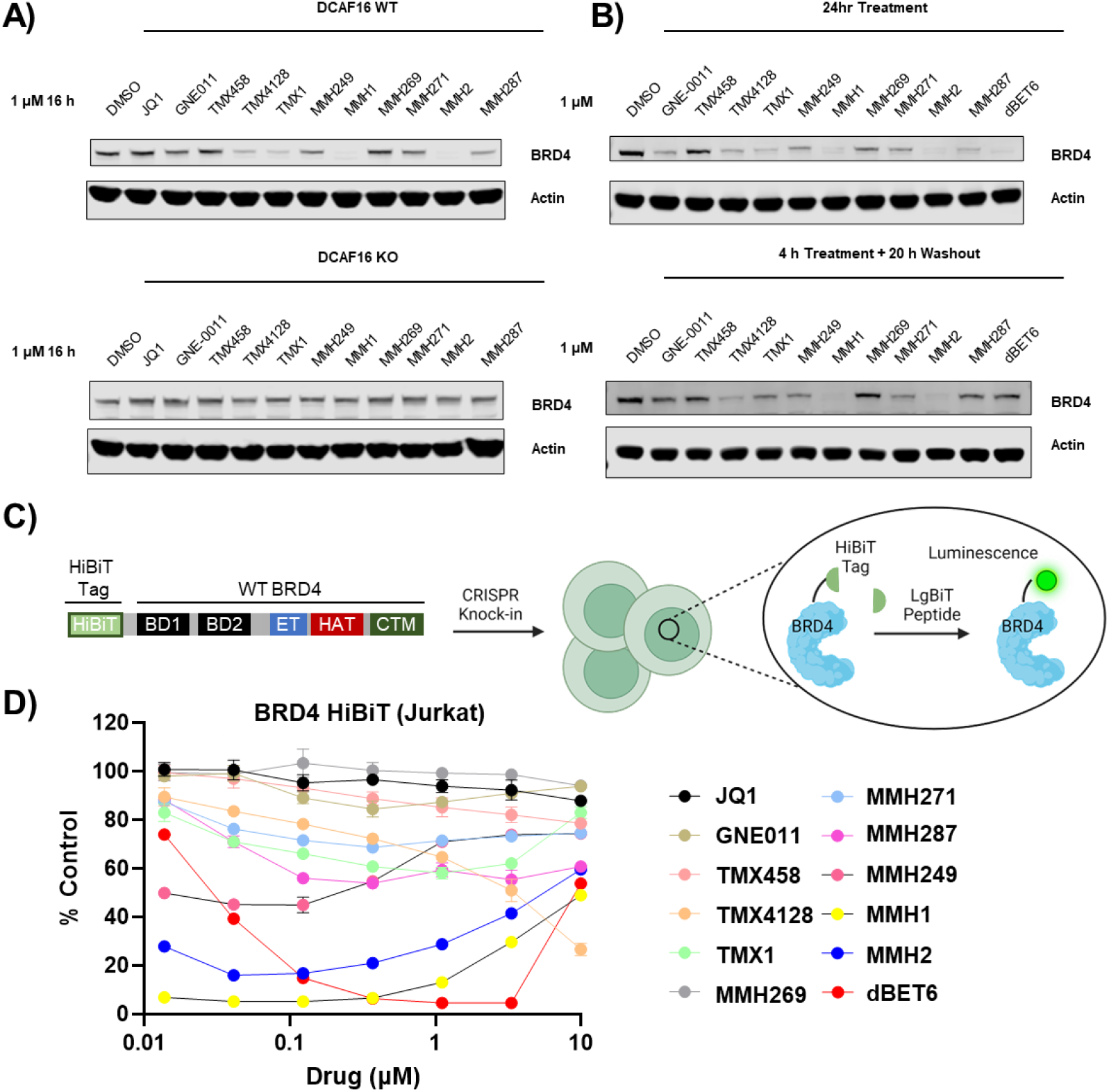
Cellular activity of the covalent BRD4 degraders. Western blotting analysis of BRD4 degradation by covalent analogs in K562 cells with A) WT and DCAF16 KO, and B) washout after 4 h of treatment (24 h total). C) A schematic depiction of the HiBiT system for quantification of protein abundance. D) Dose-dependent degradation of BRD4 by covalent JQ1 analogs in a BRD4- HiBiT JURKAT cell line. dBET6 is used as a positive control.

Upon DCAF16 covalent adduct formation by covalent JQ1 analogs, the DCAF16-JQ1 covalent adduct could presumably engage in several BRD4 degradation cycles, and thereby lead to a sustained BRD4 degradation even after removal of ‘free-compound’ from the cellular media. Hence, to confirm the DCAF16-JQ1 covalent adduct formation in cells, a washout experiment was conducted. This experiment involves cell treatment and incubation with a ‘covalent’ compound, followed by a wash-step where the cells are washed and resuspended in compound-free cell media.^29^ To confirm covalent-engagement of DCAF16 by covalent JQ1 degraders, K562 cells were treated with 1 µM concentrations, 4 h treatment time, followed by a washout and 20 h compound-free incubation. BRD4 degradation from the washout experiment was compared to a 24 h incubation without washout (Figure 4B). BRD4 degradation was completely rescued by washout for the reversible cereblon-based BRD4 degrader, dBET6. TMX1, TMX4128, MMH1, MMH2, and MMH249 showed no or minimal difference in BRD4 degradation between the washout and no-washout blots. Therefore, these compounds induce BRD4 degradation via an irreversible interaction with DCAF16 commensurate with our previous findings.^18^ Weaker degraders such as GNE011 and MMH287, showed nearly a complete rescue perhaps owing to the incubation time (4 h) being inadequate for weaker degraders to establish covalent DCAF16 covalent adducts. Surprisingly, MMH269 and MMH271 showed BRD4 degradation in 24 h no washout experiments suggesting these degraders might be slow reacting with DCAF16 in cells.

Thereafter, we sought to characterize the BRD4 cellular degradation potencies in a dose-dependent manner at a shorter time point to minimize secondary effects. This was performed using the bioluminescent HiBiT-BRD4 assay, which consists of an 11 amino acid peptide knock-in tag to a protein-of-interest using CRISPR insertion. A complementary polypeptide (LgBiT) detects and interacts with all HiBiT tagged proteins, thereby reconstituting a luminescent NanoBiT enzyme, that can be quantified by its bioluminescence.^30^ WT and DCAF16 KO JURKAT cell lines expressing the BRD4-HiBiT system were generated to quantify dose-dependent BRD4 degradation after 6 h incubation (Figure 4C). JQ1 and dBET6 were used as negative and positive controls respectively. Quantitative BRD4 degradation by dBET6 was maintained in both WT and DCAF16 KO cells. Only TMX1, TMX4128, MMH1, MMH2, MMH249, and MMH287 showed a noticeable BRD4 degradation that was completely rescued by DCAF16 KO (Figure 4D, S8). Only MMH1 and MMH2 showed robust degradation (>90% degradation at <100 nM) after 6 h, whereas modest degraders TMX1, TMX4128, and MMH249 only showed <60% degradation. Surprisingly, TMX4128 showed ∼73% degradation albeit at 10 µM, confirming the rapid DCAF16-dependent BRD4 degradation from the washout experiments at higher concentrations.

As in the BRD4_BD2_ reporter assay, neither the non-degradative covalent nor non-covalent analogs showed any significant BRD4 degradation (figure S8, S9).

## Discussion

We describe the generation of a library of BRD4 degraders through addition of a minimal covalent moiety to the reversible BRD4 inhibitor JQ1. The library consists of derivatives with varying DCAF16-dependent BRD4 degradation potencies with DC_50_ values ranging from <10 nM to > 10 µM. These were be achieved by subtle changes such as a transposition of the acrylamide warhead (ie., MMH1 vs MMH271). The C-aryl vs. *N*-aryl warhead linkages on the JQ1 scaffold lead to drastic changes in degradation potencies, with all *N*-aryl covalent analogs such as MMH1, MMH2, and MMH249, demonstrating the lowest DC_50_ values. This could be due to the surface presentation of the warhead to DCAF16, where *N-*aryl warheads could presumably furnish a greater solvent exposure (figure S7).

The high tolerance of DCAF16 for the recognition of covalent warheads on the BRD4 surface may indicate a naturally evolved function of DCAF16. Given that E3 ligases are an essential component of protein homeostasis that recognize damaged, oxidized, or dysregulated protein substates for degradation, DCAF16’s covalent sensing ability may be linked to a protein-damage or protein-oxidation response. Furthermore, DCAF16 has also been shown to endogenously ubiquitinate Spindlin 4 (SPIN4) — an epigenetic reader similar to BRD4, containing a Tudor domain which can bind trimethylated H3K4 (PDB 4UY4) — which may further indicate a natural surface complementarity with epigenetic machinery.^31^ This is reminiscent of the damage/oxidation-sensing role of Von Hippel-Lindau’s (VHL) in degrading the hypoxia inducing factor-1α (HIF-1α) upon sensing a surface proline oxidation, and of CRBN’s role in removing protein fragments with C-terminal cyclized asparagine or glutamine residues.^32,33^

Likewise, DCAF16-induced degradation of BRD4 upon binding covalent JQ1 analogs may mimic the native function of DCAF16 to prevent epigenetic reader-mediated transcriptional regulation by inducing the degradation of epigenetic reader proteins such as BRD4 and SPIN4, upon binding damaged histone lysine tails. Identifying post-translation modifications recognized by DCAF16 as a histone damage response, could serve as a promising chemical starting point to yield new and robust DCAF16 recruiters, similar to the retrospective success of thalidomide/lenalidomide and the discovery of the first VHL ligand.^10,32,34–41^

Moreover, the *trans*-labelling covalent mechanism of these JQ1 covalent glues, where a covalent ligand reversibly binds a ligandable protein ‘A’ (i.e, BRD4), yet covalently labels another protein ‘B’ (ie., DCAF16) presents the possibility of going beyond the traditional *cis*-labelling covalent pharmacology to address ‘undruggable’ proteins.

We provide a novel potential inhibitor-to-glue conversion strategy whereby covalent tagging of preexisting inhibitors could confer a gain-of-function neointeraction which can be tuned by warhead variation.

## AUTHOR INFORMATION

### Corresponding Authors

Nathanael S. Gray, Department of Chemical and Systems Biology Stanford University; Benjamin L. Ebert, Department of Medical Oncology, Dana-Farber Cancer Institute, Cancer Program, Broad Institute of MIT and Harvard. Howard Hughes Medical Institute; Eric S. Fischer, Department of Cancer Biology, Dana-Farber Cancer Institute, Department of Biological Chemistry and Molecular Pharmacology, Harvard Medical School.

## ACKNOWLEDGMENTS

This work was supported by the National Institutes of Health (NIH) grants R01HL082945, P01CA066996, P50CA206963, and R35CA253125 (to B.L.E.), and the Howard Hughes Medical Institute (to B.L.E.), NIH grants R01CA262188 and P01CA066996 (to E.S.F.), and the Mark Foundation for Cancer Research 19-001- ELA (to E.S.F.), NIH High End Instrumentation grant (1S10OD028697-01) (to N.S.G.), and departmental funds from Stanford Chemical and Systems Biology and Stanford Cancer Institute (to N.S.G). M.T. is a CPRIT Scholar in cancer research, and M.T. thanks the CPRIT for research funding support (RR220012).

## Supplementary Information for

## Supplementary Figures and Tables

**Figure S1.**
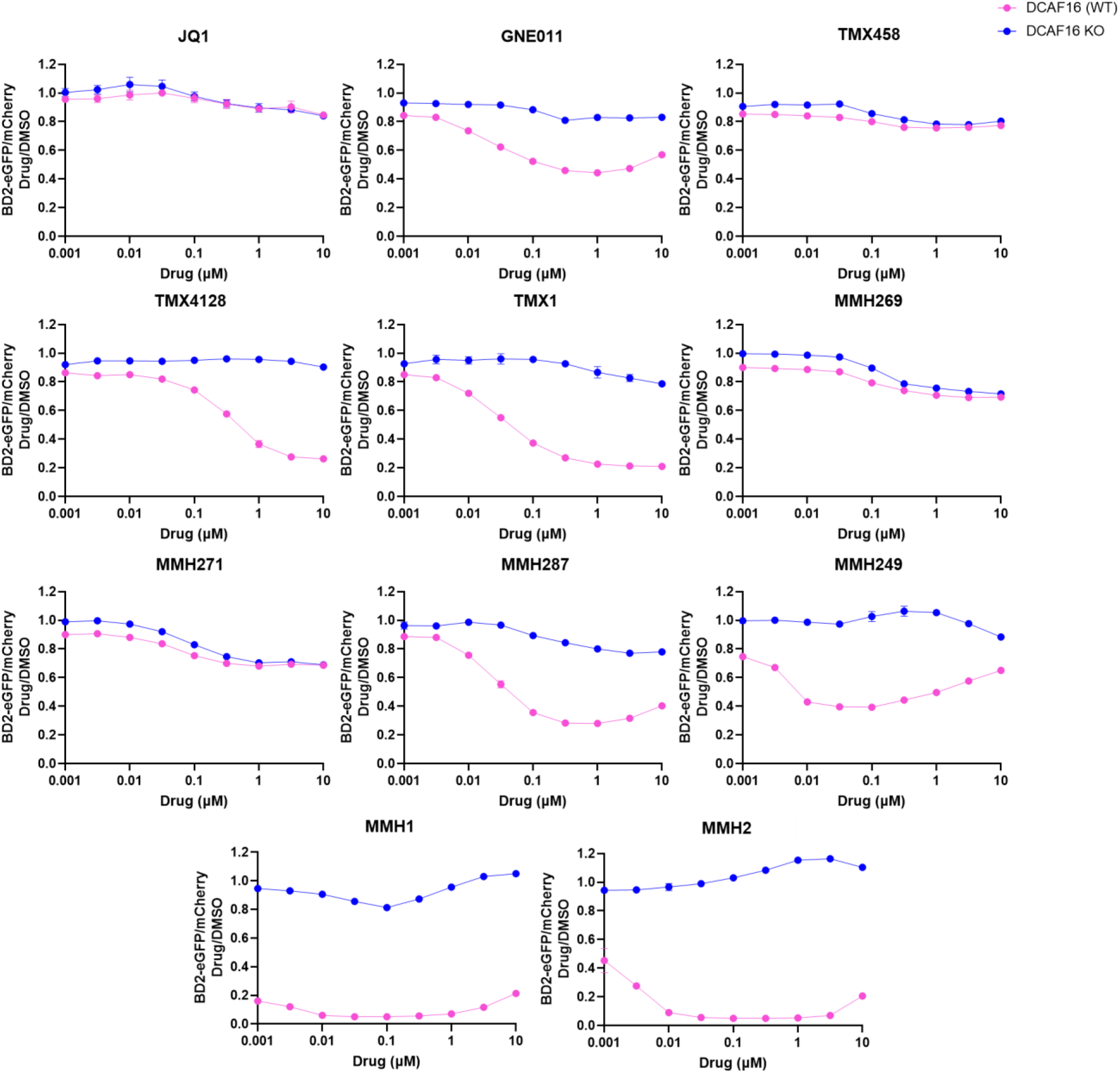
Degradation of BRD4_BD2_ in K562 cells (wild type vs. DCAF16 KO) via a BRD4_BD2_- eGFP and mCherry flow reporter assay for all covalent JQ1 analogs. JQ1 is included for comparison.

**Figure S2.**
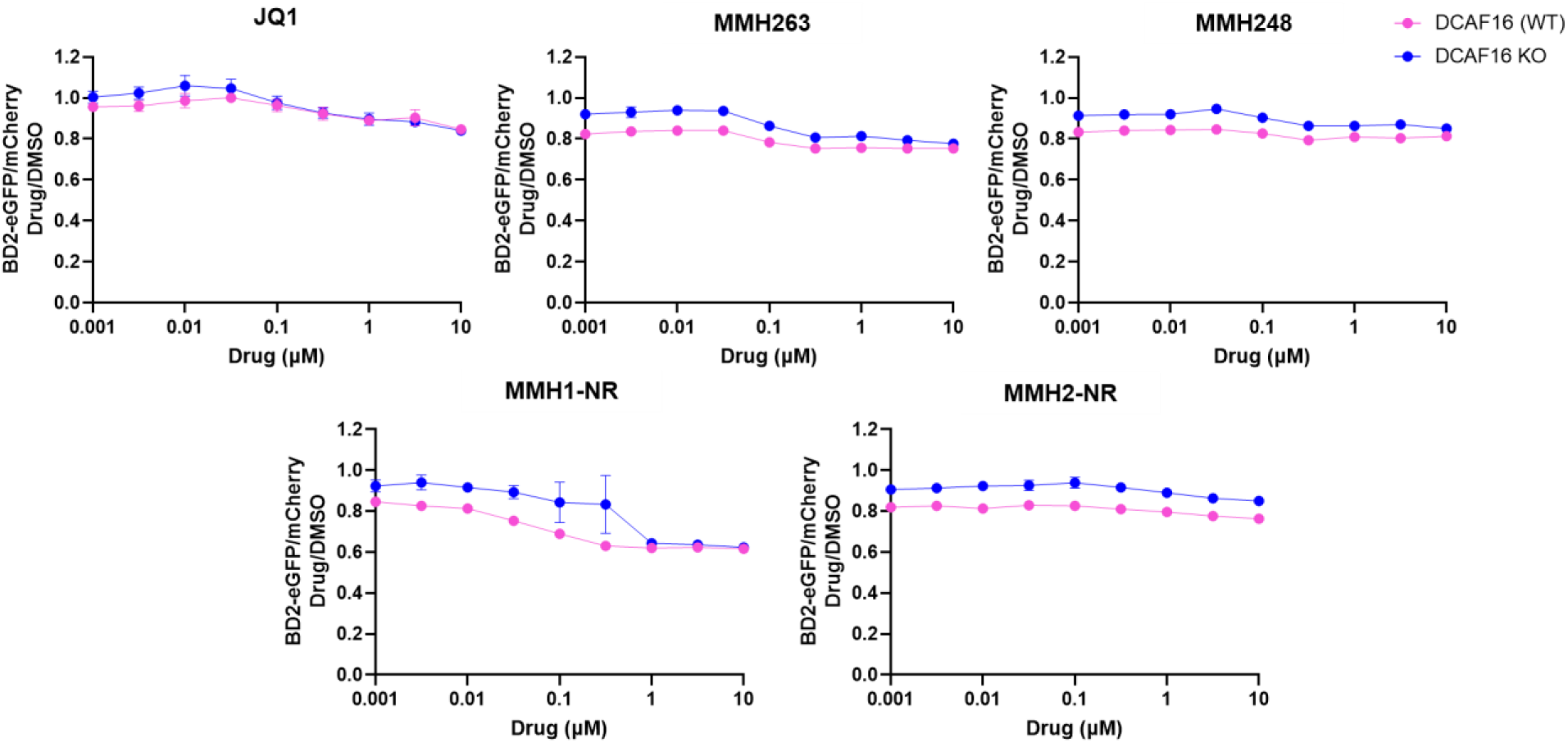
Degradation of BRD4_BD2_ in K562 cells (wild type vs. DCAF16 KO) via a BRD4_BD2_- eGFP and mCherry flow reporter assay for all non-covalent JQ1 analogs. JQ1 is included for comparison.

**Figure S3.**
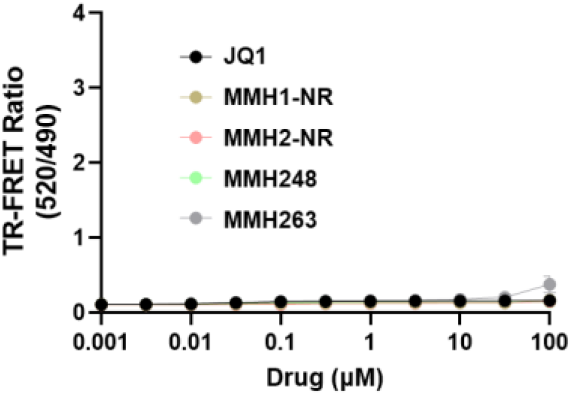
A 6 h BRD4_BD2_-DCAF16 TR-FRET assay for non-covalent analogs.

**Figure S4.**
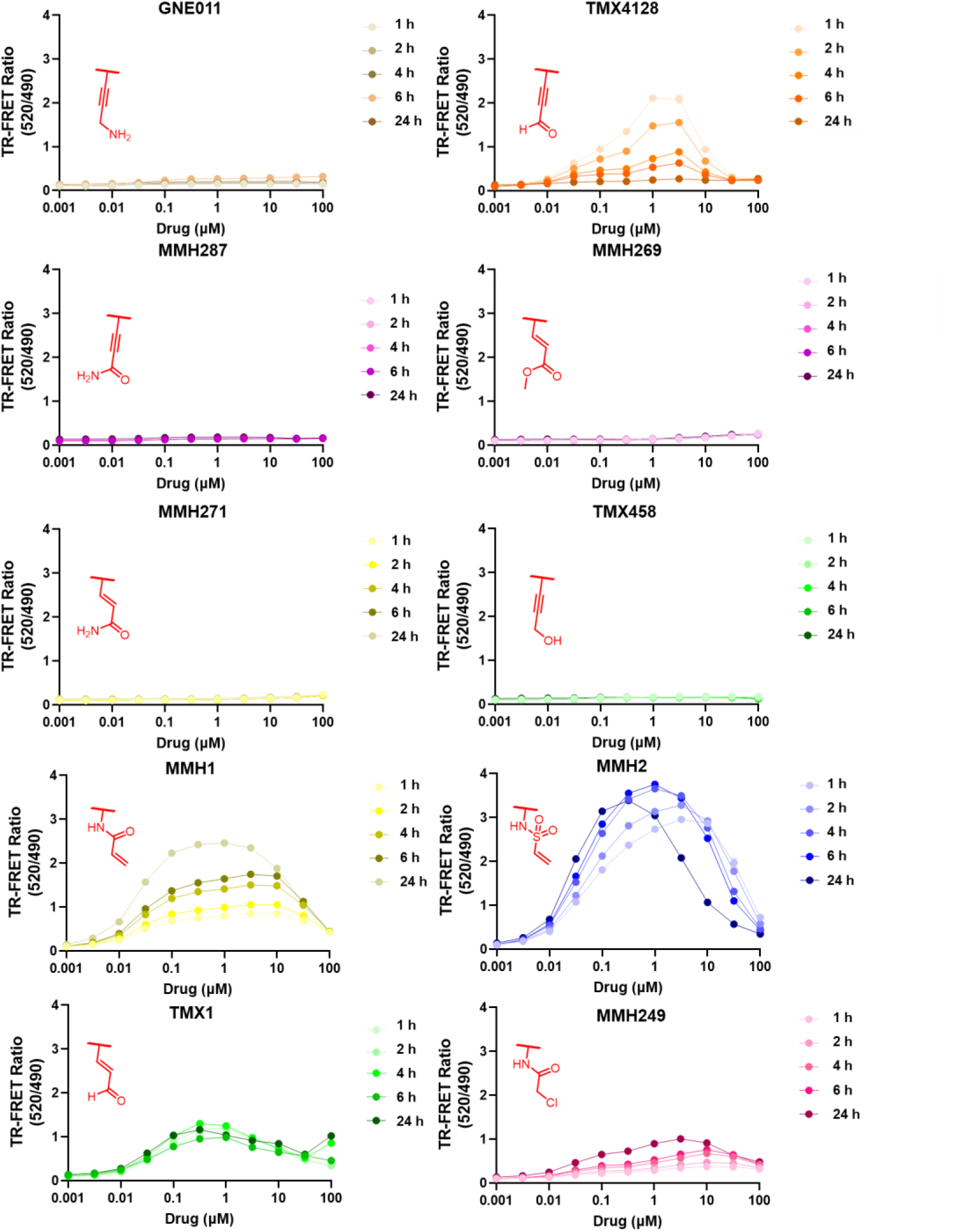
Time-dependent *in vitro* BRD4_BD2_-DCAF16 ternary complex formation for JQ1 covalent analogs using a TR-FRET assay. Four compound incubation times are used (1, 2, 4, 6, and 24 h) at 11 concentrations (0.001 to 100 µM). The covalent warhead appendages to the parent JQ1 scaffold are drawn in red.

**Figure S5.**
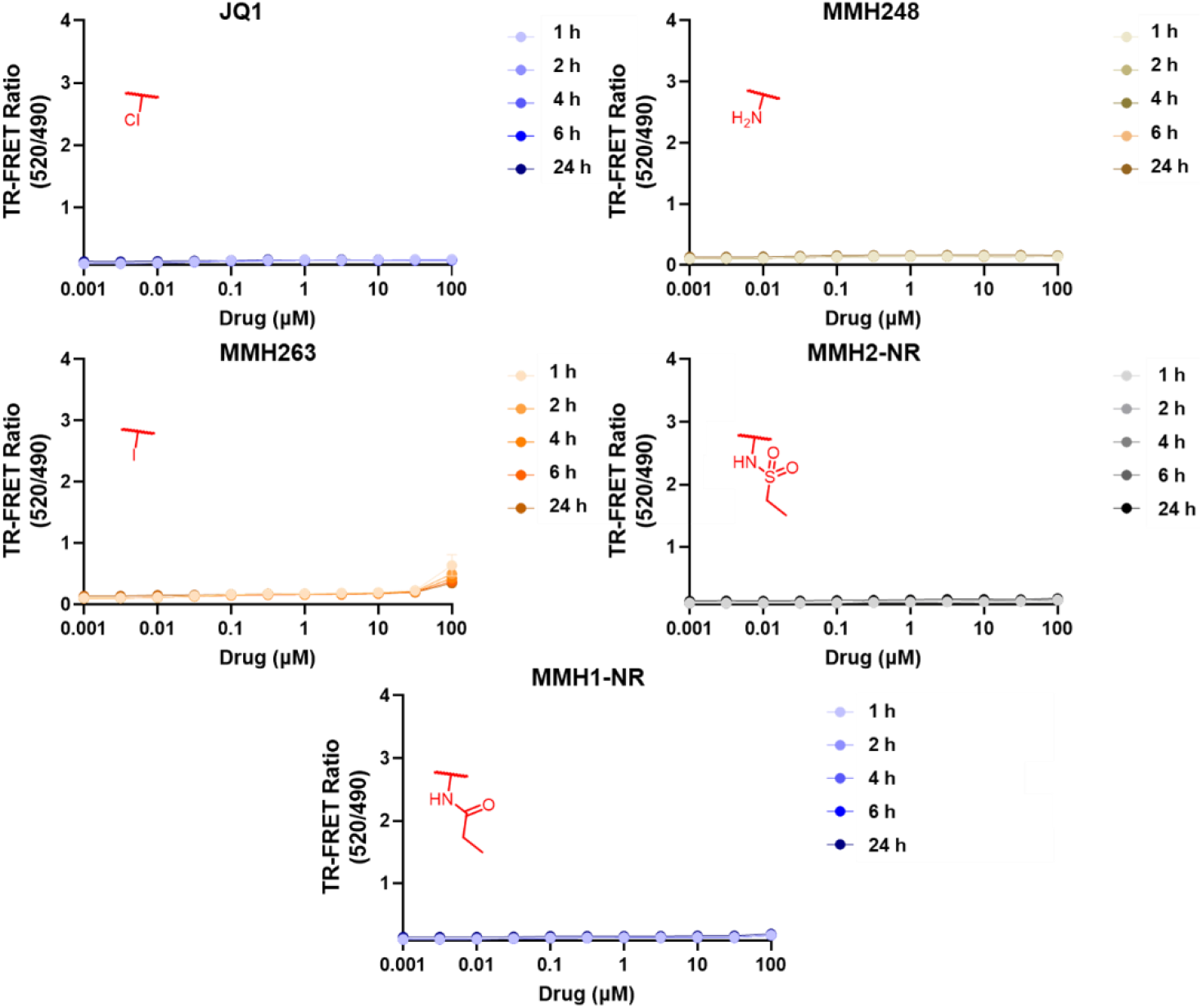
Time-dependent *in vitro* BRD4_BD2_-DCAF16 ternary complex formation for all non-covalent covalent analogs using a TR-FRET assay. Four compound incubation times are used (1, 2, 4, 6, and 24 h) at 11 concentrations (0.001 to 100 µM). The non-covalent appendages to the parent JQ1 scaffold are drawn in red.

**Figure S6.**
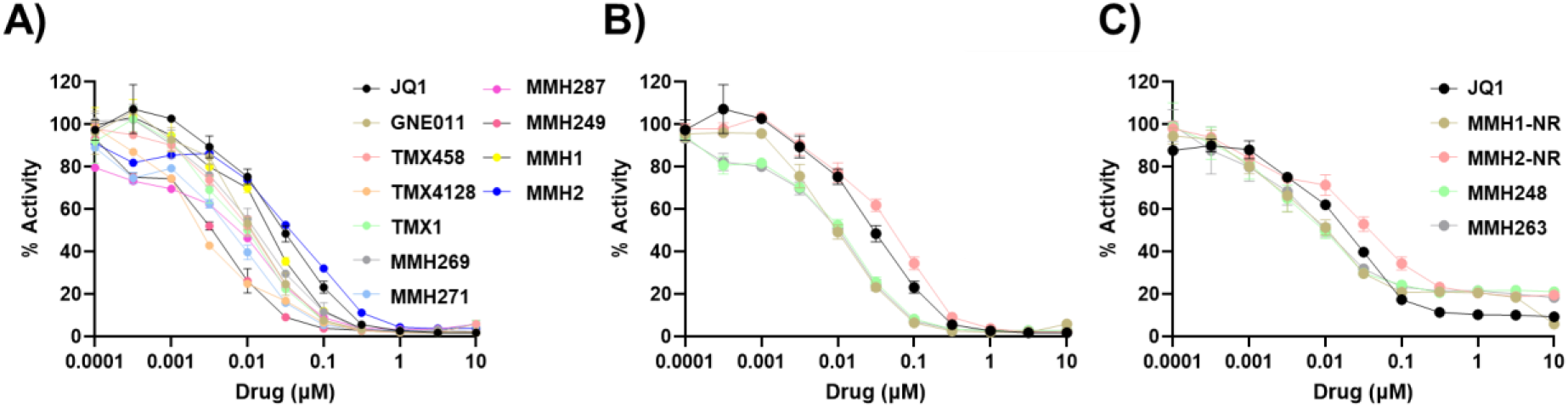
BRD4_BD1_ and BRD4_BD1_ binding potencies for JQ1 compound libraries using a competition based Alphascreen. A) BRD4_BD1_ binding potencies for covalent JQ1 analogs. B) and C) BRD4_BD1_ and BRD4_BD2_ binding potencies for non-covalent analogs.

**Figure S7.**
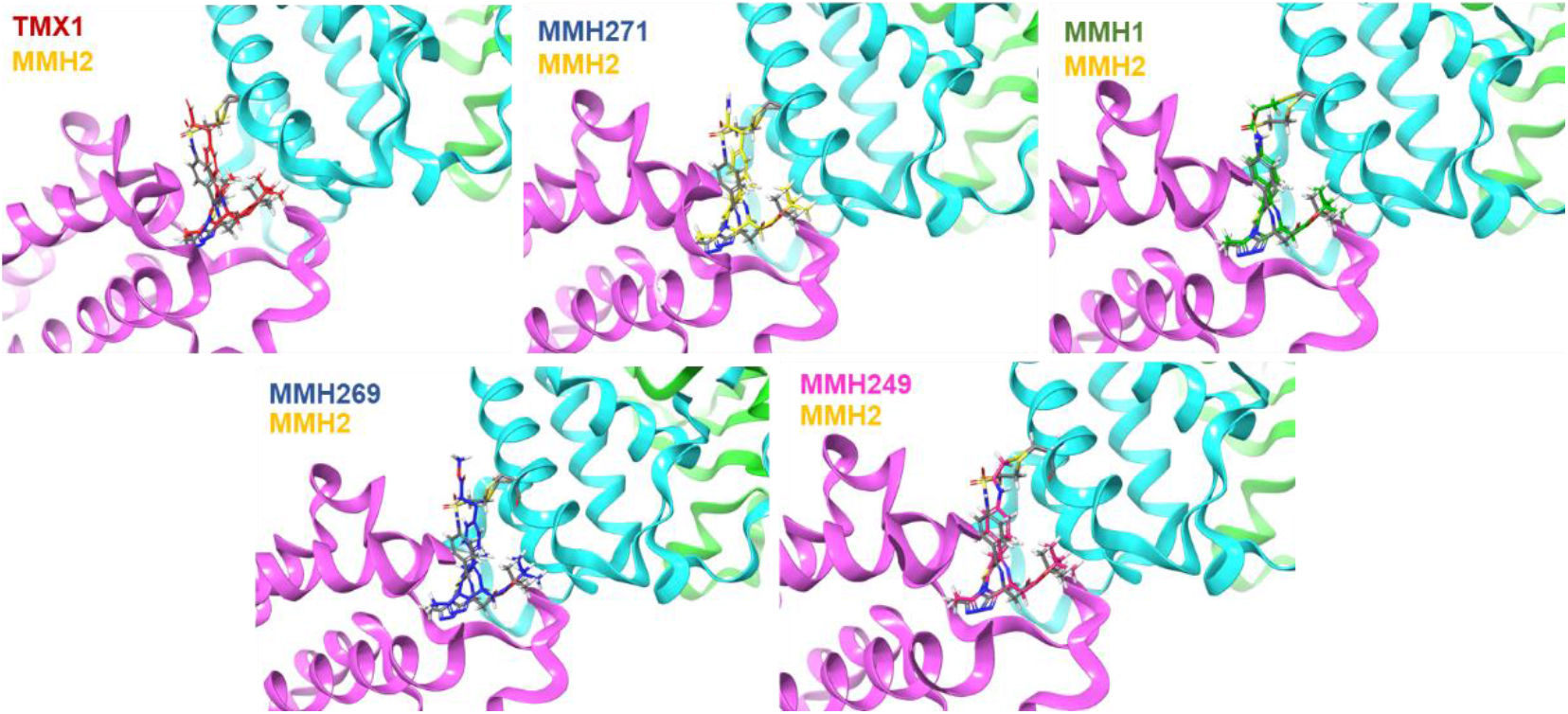
The predicted covalent docking poses of select covalent JQ1 analogs using the Schrödinger Maestro 11.9.011 software and Glide (PDB: 8G46). The docking poses of covalent JQ1 analogs are superimposed on MMH2 (yellow).

**Figure S8.**
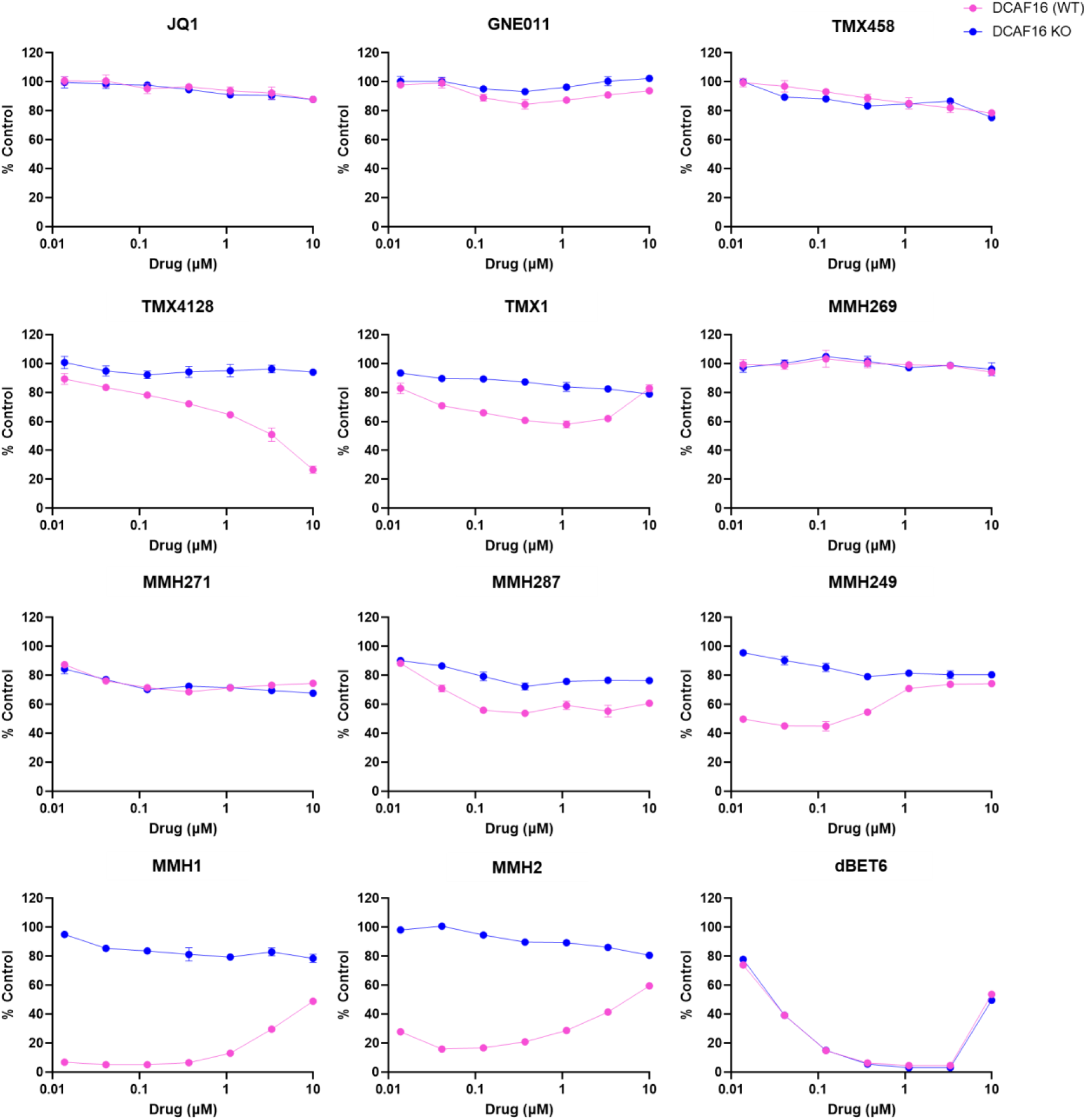
Degradation of BRD4 in JURKAT cells (wild type vs. DCAF16 KO) in a HiBiT assay (6 h) for all covalent JQ1 analogs with dBET6 as a positive control. JQ1 is included for comparison.

**Figure S9.**
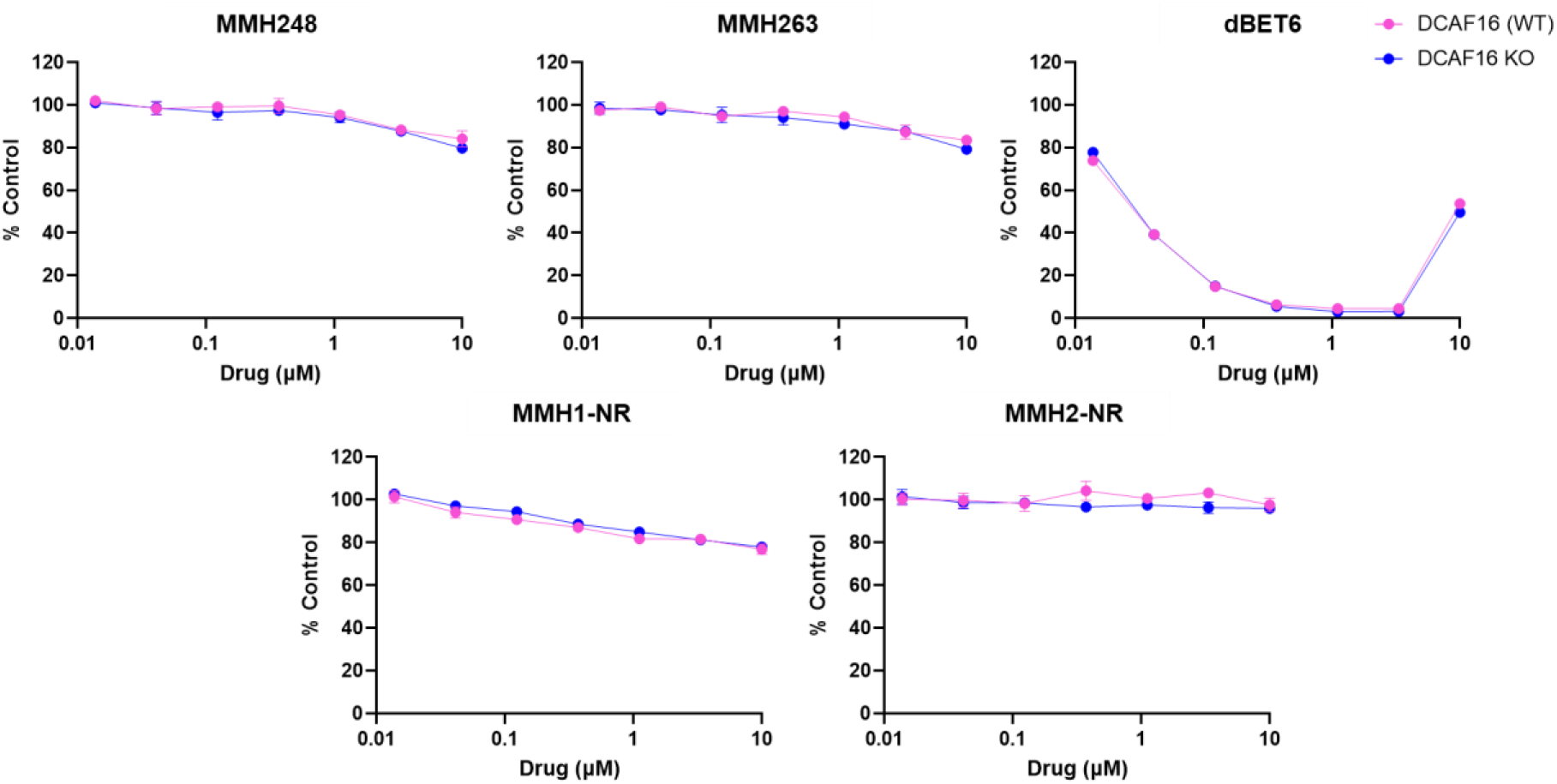
Degradation of BRD4 in JURKAT cells (wild type vs. DCAF16 KO) in a HiBiT assay (6 h) for all non-covalent JQ1 analogs, with dBET6 as a positive control.

**Figure S10.**
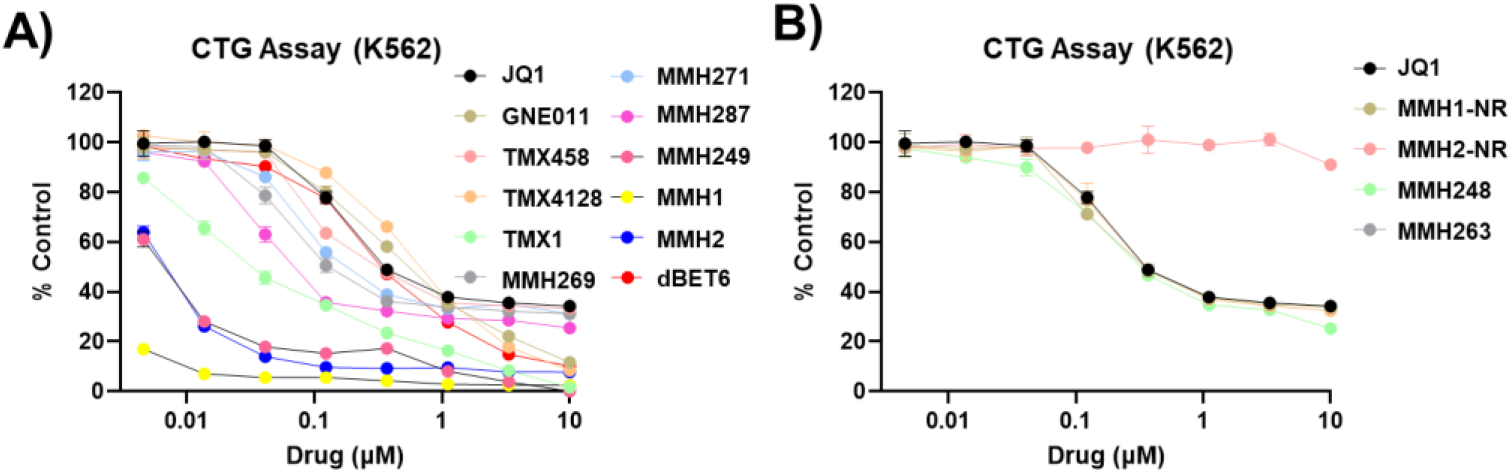
A 72 h CTG assay in K562 cells for all JQ1 analogs including both A) covalent and B) non-covalent analogs. JQ1 and dBET6 are used for comparison.

**Table S1.** IC_50_ values for all JQ1 analogs for BRD4_BD1_ and BRD4_BD2_ as determined by an AlphaScreen assay.

| Compound | IC<sub>50</sub> (nM) |  |
| --- | --- | --- |
|  | BRD4<sub>BD1</sub> | BRD4<sub>BD2</sub> |
| (+)-GNE-0011 | 11.16 | 7.79 |
| (+)-JQ1 | 26.99 | 18.21 |
| MMH1 | 17.67 | 15.50 |
| MMH2 | 48.60 | - |
| MMH248 | 12.49 | 4.29 |
| MMH249 | 3.98 | 2.59 |
| MMH263 | 11.66 | 4.96 |
| MMH269 | 10.54 | 5.07 |
| MMH271 | 8.08 | 7.58 |
| MMH277 | 9.72 | 6.00 |
| MMH284 | 47.18 | 20.43 |
| MMH287 | 12.91 | - |
| TMX1 | 8.78 | 33.86 |
| TMX4128 | 2.20 | 21.70 |
| TMX458 | 10.14 | 3.92 |

**Table S2.**
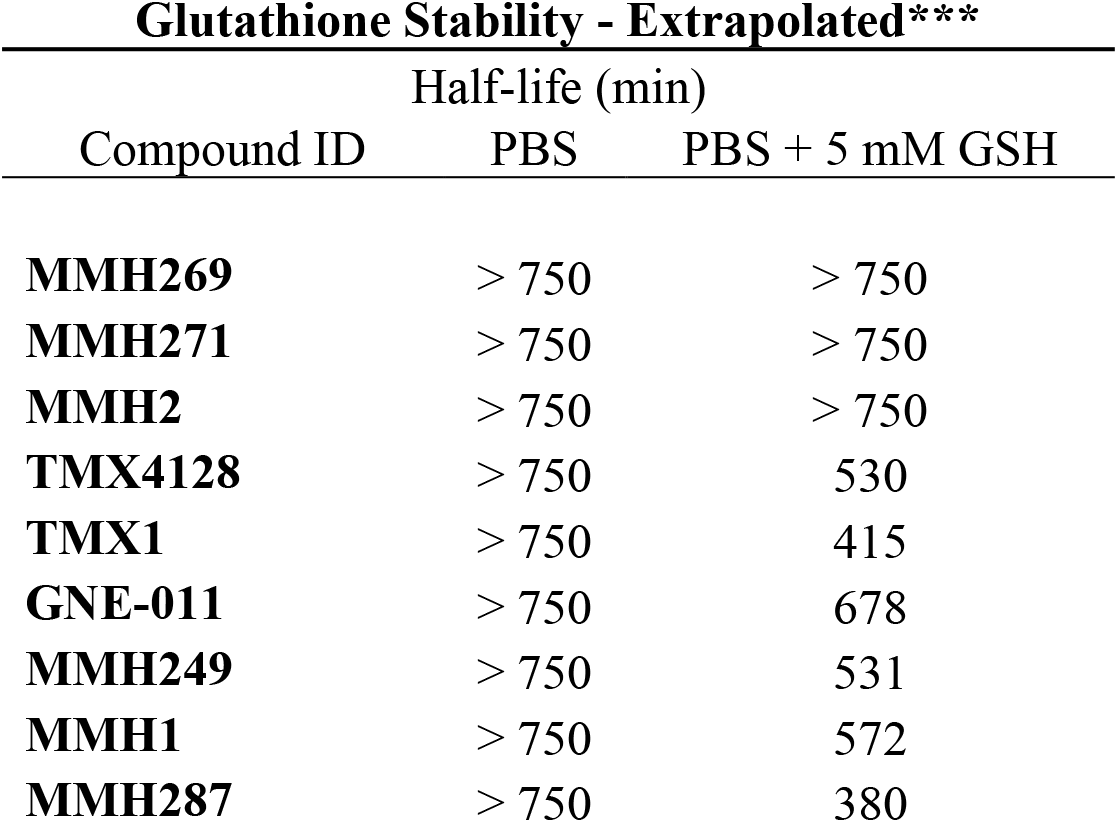
The half-lives of JQ1 covalent analogs extrapolated from a 240 min Glutatione reactivity assay, where a >750 min half-life would result in >20% depletion in 240 min of GSH incubation with covalent analogs.

## Materials and Methods

### Cell culture

Human leukemia cell lines (JURKAT and K-562) were obtained from the American Type Culture Collection (ATCC, Manassas, VA, USA) and K562-Cas9 cell line was provided by Zuzana Tothova (Dana-Farber Cancer Institute). The cells were cultured in RPMI 1640 medium supplemented with 10% heat-inactivated FBS (Gibco, Grand Island, NY, USA), 100 units/mL penicillin, 100 µg/mL streptomycin, 0.25 µg/mL amphotericin B, and 5% glutamine. Incubation of the cells was carried out at 37 °C with 5% CO_2_ in a humidified environment. Regular mycoplasma testing was conducted using the MycoAlert mycoplasma detection kit (Lonza, Basel, Switzerland), and all cell lines tested negative for mycoplasma contamination.

### HiBiT assay

HiBiT-edited JURKAT cells were plated in white 384-well cell culture plates (Corning #3570) at a density of 2 × 10^4^ cells per well in 50 μL of growth medium and incubated with indicated concentrations of compounds. After 6 h, the plates were subjected to Nano-Glo HiBiT Lytic detection system (Promega, #N3040) as described in manufacturer’s manual. The assays were performed in biological triplicate. DC_50_ values were determined using a non-linear regression curve fit in GraphPad PRISM 9.5.1.

### Cell viability assay (CellTiter-Glo assay)

Cell viability was evaluated using the CellTiter-Glo assay (Promega, Madison, WI, USA). Briefly, K-562 cells (1000 cells/well) were seeded in 384-well cell culture plates (Corning, #3570) and then cells were incubated with the indicated concentrations of compounds. After 72 h, the plates were subjected to CellTiter-Glo assay as described in manufacturer’s manual. The proliferation assays were performed in biological triplicate. IC_50_ values were determined using a non-linear regression curve fit in GraphPad PRISM 9.5.1.

### Antibodies

The following antibodies were used: anti-BRD4 (Bethyl Laboratories, A301-985A100), anti-β- actin (Cell Signaling Technology, #3700), anti-mouse 800CW (LI-COR Biosciences, 926-32211), anti-rabbit 680LT (LI-COR Biosciences, 925-68021).

### Commercial Compounds

JQ1 (HY-13030), dBET6 (HY-112588) were obtained from MedChemExpress.

### Plasmids

The following plasmids were used in this study: Cilantro (PGK.BsmBICloneSite.FlexibleLinker.eGFP.IRES.mCherry.cppt.EF1α.PuroR, Addgene 74450) for degradation characterization; sgBFP (U6.sgRNA.cppt.SFFV.tBFP) for validation of DCAF16 knockout phenotypes. The following plasmids were used for the TR-FRET assay: StrepII-Avi-DCAF16 in a pAC-derived vector, His6-3C-Spy-DDB1ΔB in a pAC-derived vector, and His6-Avi-BRD4_BD2_ in E. Coli pET100/D-TOPO vector.

### Immunoblots

Cells were washed with PBS and lysed in RIPA lysis buffer (Thermo Fisher Scientific) with Halt Protease Inhibitor Cocktail (Thermo Fisher Scientific) and Benzonase (Sigma-Aldrich) for 20 min on ice. The insoluble fraction was removed by centrifugation, the protein concentration was quantified using a BCA protein assay kit (Thermo Fisher Scientific), and an equal amount of lysate was run on SDS–PAGE 4–12% Bis–Tris Protein Gels (Thermo Fisher Scientific) and then transferred to nitrocellulose membrane with a XCell II Blot Module Wet Tank Transfer System (Thermo Fisher Scientific). Membranes were blocked in Intercept (PBS) Blocking Buffer (LI-COR Biosciences) and incubated with primary antibodies overnight at 4 °C. The membranes were then washed in Tris-buffered saline with Tween-20 (TBS-T), incubated for 1 h with secondary IRDye-conjugated antibodies (LI-COR Biosciences) and washed three times in TBS-T for 5 min before near-infrared western blot detection on an Odyssey Imaging System (LI-COR Biosciences).

### Reporter cell line generation

Reporter constructs were generated by BsmBI (New England Biolabs) digestion of Cilantro reporter vector and the insert containing protein of interest coding sequence, followed by ligation with T4 DNA Ligase (New England Biolabs). Constructs were transformed into Stbl3 E. coli and purified using the MiniPrep Kit (Qiagen), and sequences were confirmed by Sanger sequencing (Quintara Biosciences Service). Lentiviruses for reporters were packaged into lentivirus as follows. 0.5 × 10^6^ HEK293T cells were seeded in 2 mL of DMEM media. The next day, a packaging mix including 1.5 μg of psPAX2, 0.15 μg of pVSV-G, and 1.5 μg of transgene plasmid was prepared in 37.5 µL of OptiMEM (Thermo Fisher Scientific). This mix was combined with 9 μL of TransIT-LT1 (Mirus) and 15 µL of OptiMEM, incubated for 30 min at room temperature, and then applied dropwise to cells. Cells were allowed to incubate for another 48 h. Lentivirus was collected by 0.4 μM filters, and then transduced to 2 × 10^6^ of K562-Cas9 cells at 50% volume ratio by spin infection. One day after infection, reporter cells were selected with puromycin at a concentration of 2 μg/mL.

### DCAF16 knockout cell line generation

sgRNAs targeting DCAF16 (sgDCAF16) or control (sgNTC) were cloned into the sgBFP vector using BsmBI cloning. In brief, vectors were linearized with BsmBI (New England Biolabs) and gel-purified with QIAquick Gel Extraction Kit (Qiagen). Annealed oligos containing sgRNA sequences were phosphorylated with T4 polynucleotide kinase (New England Biolabs) and ligated into linearized vector backbone. sgRNA constructs were transformed, purified, and verified, and lentivirus was generated as described above. Lentivirus containing sgRNA was transduced to 2 × 10^6^ of K562-Cas9 cells at 10% volume ratio by spin infection. FACS sorting was performed to enrich BFP+ cells one week after infection. For the generation of single-clone DCAF16 knockout cells, pooled K562-Cas9 cells stably expressing sgRNA targeting DCAF16 were seeded in 384- well plates at the density of 0.25 cells per well. Clonal sgDCAF16-expressing K562-Cas9 cells were isolated after one month of expansion, and the genomic sequences were validated via deep sequencing of PCR amplicons targeting sgDCAF16 cutting sites (MGH CCIB DNA Core Service).

### Flow Reporter degradation assays

K562 cells stably expressing degradation reporter were dosed with DMSO or degraders at various times and concentrations using D300e Digital Dispenser (HP). The fluorescent signal was quantified by flow cytometry (LSRFortessa flow cytometer, BD Biosciences) and analyzed using FlowJo (flow cytometry analysis software, BD Biosciences). The geometric mean of the eGFP and mCherry fluorescent signal for round and mCherry-positive cells was calculated. GFP expression was normalized to mCherry signal and drug treatments were compared to DMSO controls.

### *In silico* Modelling

The cryo-EM structure of MMH2 induced BRD4_BD2_-DCAF16-DDB1 ternary structure (PDB 8G46) was used for pose-prediction of the covalent analogs. Default settings for ligand preparation, and protein preparation and refinement were used. Cys58 was selected as the reactive residue. The centroid of the workspace ligand was selected to be that of MMH2. The Michael addition reaction type was chosen, with a ‘pose-prediction’ Docking Mode. The output poses per ligand was restricted to 5.

### Intact protein mass spectrometry

Prior to intact mass analysis, recombinant human DDB1ΔB-DCAF16 variants were incubated with DMSO, TMX1, KB02-JQ1, MMH1, or MMH2 with and without the presence of recombinant human BRD4BD2 for 16 h at 4°C. For GNE11, recombinant proteins were incubated with drug at room temperature for 16 h. Intact mass analysis of DCAF16 variants was performed similarly to a previously described protocol49 with modifications. Briefly, drug-treated proteins were injected on a self-packed column (6 cm POROS 50R2 packed in 0.5 mm I.D. tubing), desalted for 4 minutes, and then eluted to an LTQ ion trap mass spectrometer (Thermo Fisher Scientific) using an HPLC gradient (0-100% B in 20 minutes, A=0.1M acetic acid, B=0.1 M acetic acid in acetonitrile, ESI spray voltage=5kV). The mass spectrometer acquired full scan mass spectra (m/z 300-2000) in profile mode. Mass spectra were deconvoluted using MagTran version 1.03 b250. Labeling efficiency was calculated from zero charge mass spectra using peak heights according to [peak height labeled protein] / [peak height labeled protein + peak height unlabeled protein] x 100%.

### DDB1-DCAF16–BRD4_BD2_ TR-FRET

Titrations of compounds to induce the DCAF16-BRD4BD complex were carried out by mixing 100 nM biotinylated BRD4_BD2_, 500 nM BODIPY-FL labeled DDB1ΔB-DCAF16 variants, and 2 nM terbium-coupled streptavidin (Invitrogen) in an assay buffer containing 50 mM HEPES pH 8.0, 200 mM NaCl, 0.1% Pluronic F-68 solution (Sigma), 0.5% bovine serum albumin (BSA) (w/v) and 1 mM TCEP. After dispensing the assay mixture (15 μL volume), increasing concentrations of compounds were dispensed in a 384-well microplate (Corning, 4514) using a D300e Digital Dispenser (HP) normalized to 1% DMSO. After excitation of terbium fluorescence at 337 nm, emission at 490 nm (terbium) and 520 nm (BODIPY FL) were recorded with a 70 μs delay over 600 μs to reduce background fluorescence, and the reaction was followed over 60 cycles of each data point using a PHERAstar FS microplate reader (BMG Labtech). The TR-FRET signal of each data point was extracted by calculating the 520/490 nm ratio.

### GSH Stability Assay

Individual compound spiking solutions were prepared at 100 µM in DMSO and spiked into glass HPLC autosampler vials containing either phosphate buffered saline (PBS) or PBS containing 5 mM reduced glutathione. The samples were immediately injected onto an LC-MS/MS to determine reaction at time zero. At specific timepoints over four hours, the samples were reinjected to determine compound depletion.

### BRD4 AlphaScreen

Assays were performed with minimal modifications from the manufacturer’s protocol (PerkinElmer, USA). All reagents were diluted in 50 mM HEPES, 150 mM NaCl, 0.1% w/v BSA, 0.01% w/v Tween20, pH 7.5, and allowed to equilibrate to room temperature prior to addition to plates. After addition of Alpha beads to master solutions, all subsequent steps were performed under low light conditions. A 2x solution of components with final concentrations of His-BRD4- BD1 or His-BRD4-BD2 at 40 nM, Ni-coated Acceptor Bead at 15 µg/ml, and biotinylated-JQ1 at 20nM was added in 10 µL to 384-well plates (AlphaPlate-384, PerkinElmer, USA). Plates were spun down at 150x g, and 100 nL of compound in DMSO from stock plates were added by pin transfer using a Janus Workstation (PerkinElmer, USA). Streptavidin-coated donor beads (15 µg/ml final concentration) were added as to the solution in a 2x, 10 µL volume. Following this addition, plates were sealed with foil to prevent light exposure and evaporation. The plates were spun down again at 150g. Plates were incubated at room temperature for 1 hour, and then read on an Envision 2104 (PerkinElmer, USA), using the manufacturer’s protocol.

## Synthetic Methods for

### Chemical Procedures

#### General Procedure

All solvents and chemical reagents were purchased from commercial vendors such as Sigma Aldrich, Oakwood Chemicals, Fischer Scientific, Combi Blocks, TCI Chemicals, and Alfa Aesar. Anhydrous solvents such as THF, and DMF were used from Sure-Seal bottles without further drying treatments. All reactions were conducted in a scintillation vial and were monitored by LCMS. Compound purity was determined by the relative area of compound peaks as determined by the automatic integration of the UV spectra.

#### Analytical Conditions and Instrumentation

All NMR spectra were obtained on a 500 MHz Bruker Avance III spectrometer or on 500 MHz Bruker Avance Neo equipped with carbon detect, liquid nitrogen cooled, Prodigy cryoprobe. All deuterated solvents were purchased from either Sigma Aldrich or Cambridge isotope laboratories. Coupling constants (*J*) are reported in Hertz (Hz) and multiplicities are reported as singlet (s), doublet (d), triplet (t), quartet (q), pentet (p), sextet (sex), septet (sep), multiplet (m), and broad (br), and the combination of these are listed as a combination of their abbreviations.

LCMS was conducted on an Acquity I-Class LCMS system equipped with a PDA-UV system, a QDa mass detector, and a dual column set-up (BEH and CSH columns). The solvent gradient consisted of LCMS grade acetonitrile purchased from Fischer Scientific, and milliQ water. A gradient of 10 to 100% over 2.5 minutes followed by a 0.5 minute flush, was used for the final compounds described herein.

Preparative HPLC was used to purify all final compounds unless otherwise stated. HPLC grade methanol purchased from Fischer Scientific, and milliQ water, both containing 0.045% trifluoroacetic acid (TFA) were used for the solvent system. A 254 nm UV-light absorption was used to visualize compound chromatograms. Two methods were used for the solvent gradient system:

Method 1: Waters Sunfire C18 column (19 mm X 50 mm, 5 μm) using a gradient of 15-95% methanol in water over 60 min at a flow rate of 43 mL/min.

Method 2: Waters Sunfire C18 column (30 mm X 250 mm, 5 µm) using a gradient of 10 to 100% methanol in water containing 0.045% TFA over 40 min followed by a 5 min flush with 100% methanol, at a flow rate of 40 mL/min.

## Abbreviations

B_2_pin_2_: bis(pinacolato)diboron
Pd_2_dba_3_: tris(dibenzylideneacetone)dipalladium(0)
XPhos: dicyclohexyl[2′,4′,6′-tris(propan-2-yl)[1,1′-biphenyl]-2-yl]phosphane
KOAc: potassium acetate
NaI: sodium iodide
Chloramine-T: sodium chloro(4-methylbenzene-1-sulfonyl)azanide
THF: tetrahydrofuran
PdCl_2_(PPh_3_)_2_: bis(triphenylphosphine)palladium(II) dichloride
Et_3_N: triethylamine
DMF: dimethylformamide
rt: room temperature
K_2_CO_3_: potassium carbonate
KCl: potassium chloride
[n-Bu_4_N]OAc: tetrabutylammonium acetate
Pd(OAc)_2_: palladium(II) acetate
Xphos-PdG2: Chloro(2-dicyclohexylphosphino-2′,4′,6′-triisopropyl-1,1′-biphenyl)[2-(2′- amino-1,1′-biphenyl)]palladium(II)
DCM: dichloromethane
DIPEA: diisopropylethylamine
prep: preparative
LC: liquid chromatography
MS: mass spectrometry.

### Synthetic Scheme 1. Synthesis of Aryl-linked JQ1 Analogs

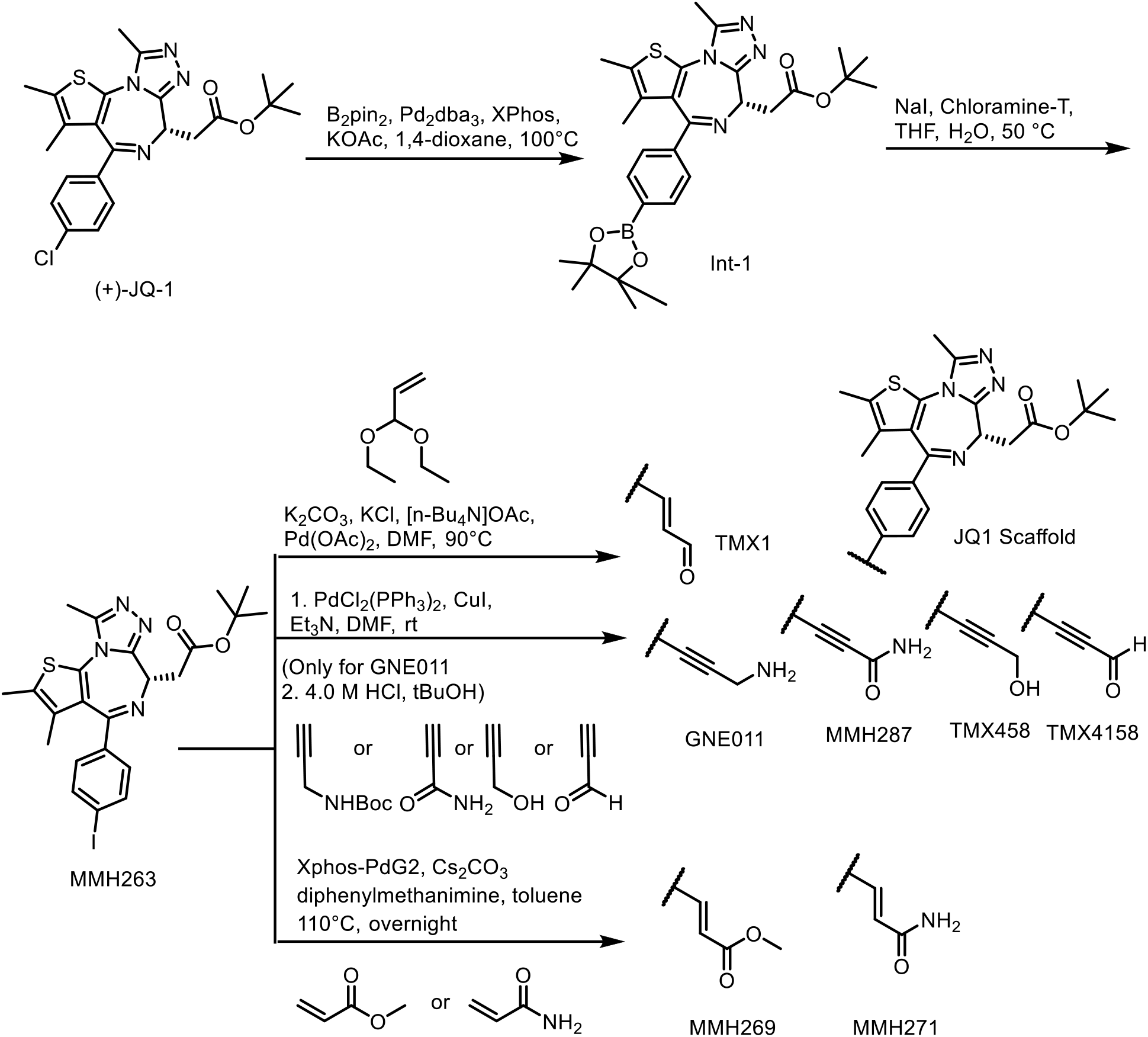

### Synthetic Scheme 2. Synthesis of N-Aryl-linked JQ1 Analogs

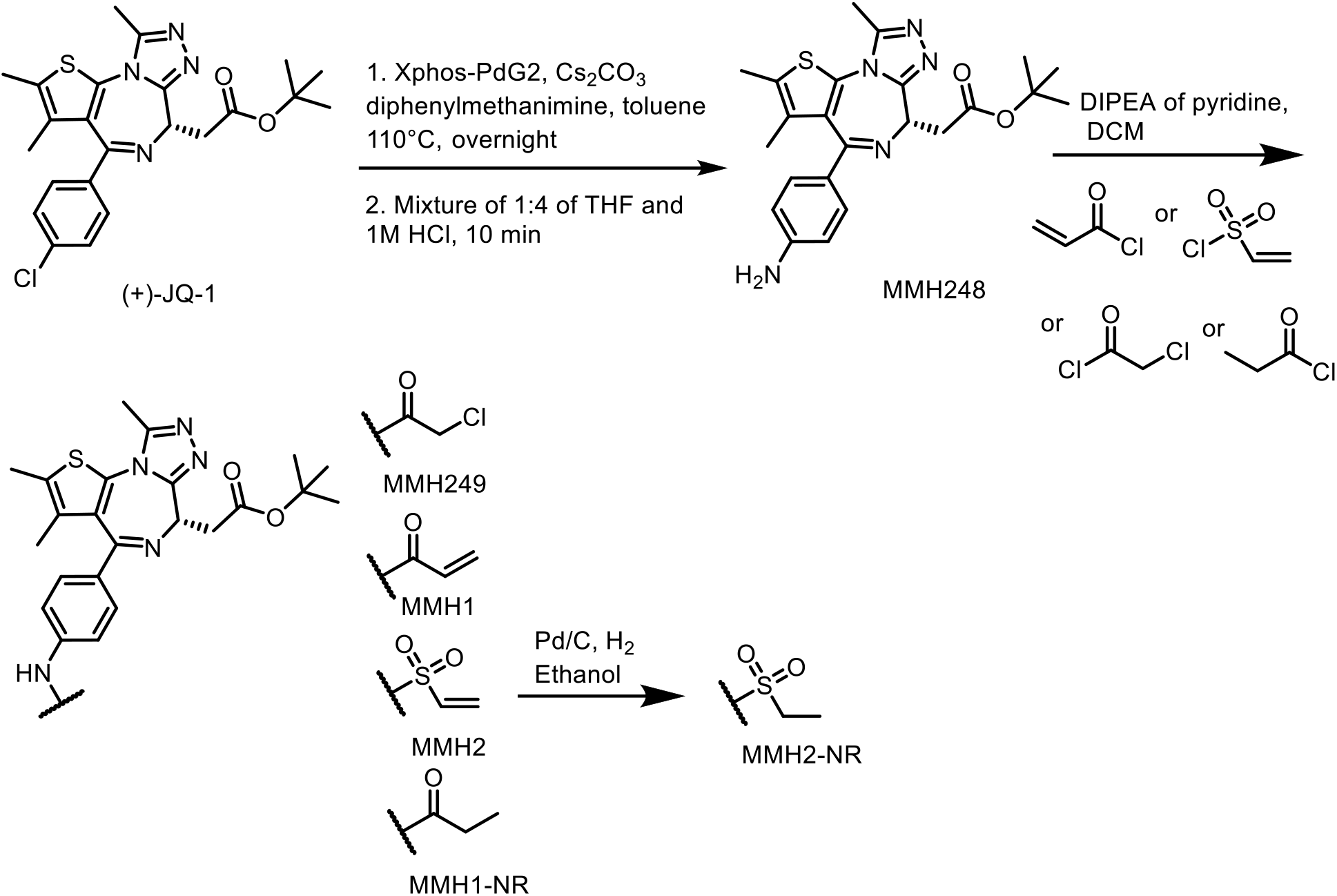

## Chemical Synthesis and Compound Characterization

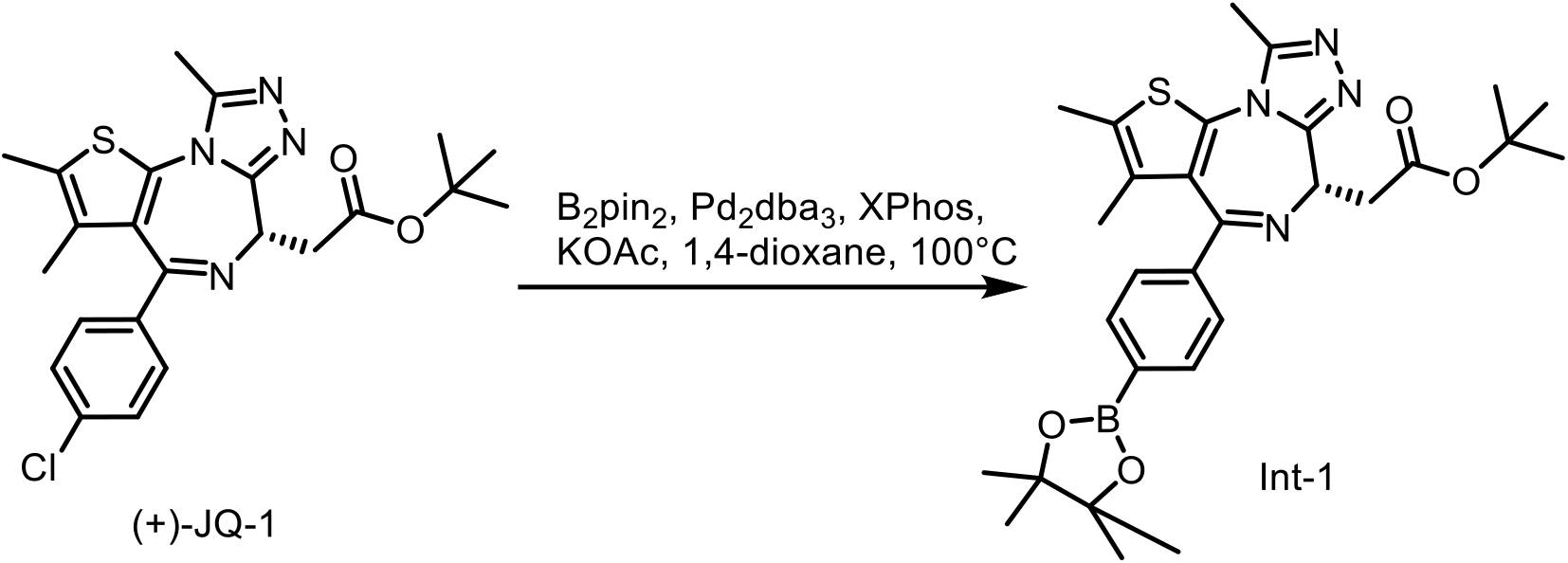

**tert-butyl (S)-2-(2,3,9-trimethyl-4-(4-(4,4,5,5-tetramethyl-1,3,2-dioxaborolan-2-yl)phenyl)- 6H-thieno[3,2-f][1,2,4]triazolo[4,3-a][1,4]diazepin-6-yl)acetate (Int-1)**

To a solution of (+)-JQ-1 (600.0 mg, 1.31 mmol) and B_2_pin_2_ (332.0 mg, 1.31 mmol) in 1,4-dixoane (5.0 mL) was added XPhos (31.0 mg, 0.065 mmol), Pd_2_dba_3_ (30.0 mg, 0.033 mmol), and KOAc (386 mg, 3.93 mmol). The reaction mixture was stirred at 100 °C for 17 hours. The reaction mixture was purified via column chromatography (silica gel, eluted with 0% to 15% methanol in dichloromethane) to give **Int-1** (395 mg, 64% yield) as a yellow oil. For **Int-1**, MS (ESI) for C_29_H_38_BN_4_O_4_S [M+H]^+^: m/z calcd, 549.51; found, 549.49.

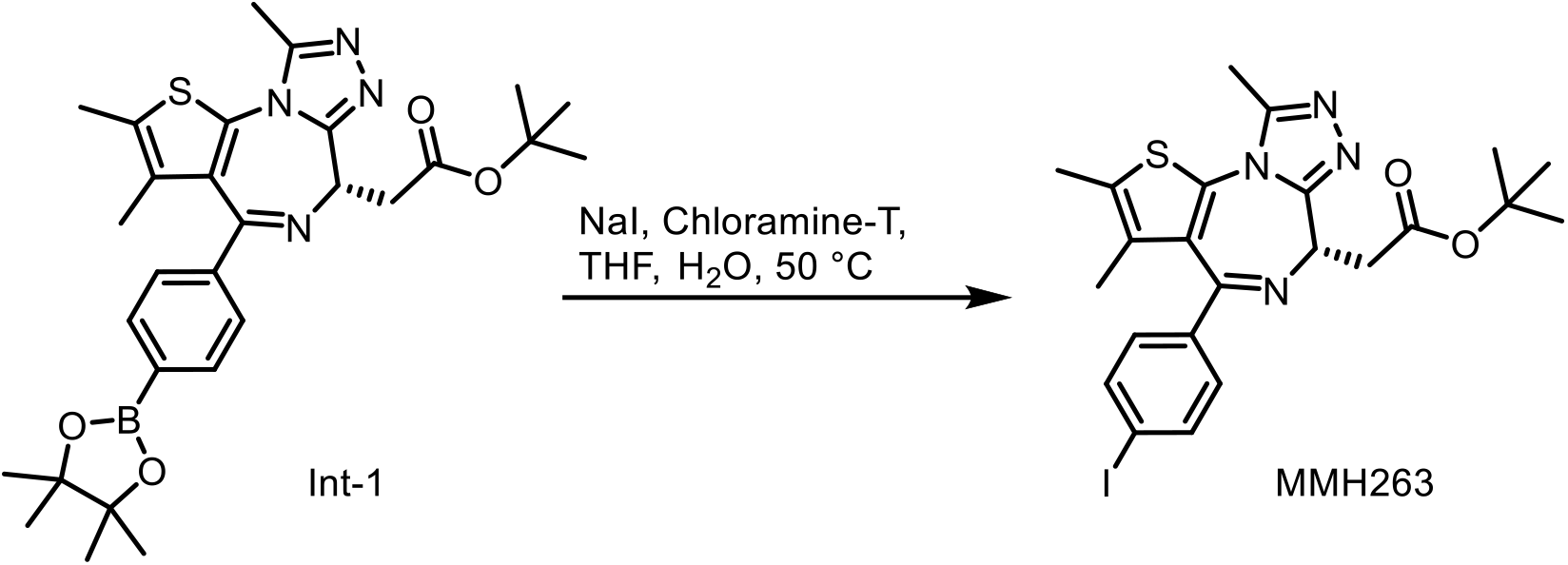

**tert-butyl (S)-2-(4-(4-iodophenyl)-2,3,9-trimethyl-6H-thieno[3,2-f][1,2,4]triazolo[4,3- a][1,4]diazepin-6-yl)acetate (MMH263)**

To a solution of **Int-1** (200.0 mg, 0.43 mmol) in 1,4-dixoane/H_2_O (v/v=2/1, 3.0 mL) was added NaI (129.0 mg, 0.86 mmol) and Chloramine-T 294.0 mg, 1.29 mmol). The reaction mixture was stirred at 50 °C for 4 hours. The reaction mixture was purified directly via prep HPLC (method 1) to give **MMH263** (142 mg, 60% yield) as a yellow solid. ^1^H NMR (500 MHz, DMSO-*d*_6_) δ 7.81 (d, *J* = 8.5 Hz, 2H), 7.20 (d, *J* = 8.0 Hz, 2H), 4.40 (dd, *J* = 8.0, 6.5 Hz, 1H), 3.35-3.29 (m, 2H), 2.60 (s, 3H), 2.41 (s, 3H), 1.63 (s, 3H), 1.42 (s, 9H). MS (ESI) for C_23_H_26_IN_4_O_2_S [M+H]^+^: m/z calcd, 549.45; found, 549.38.

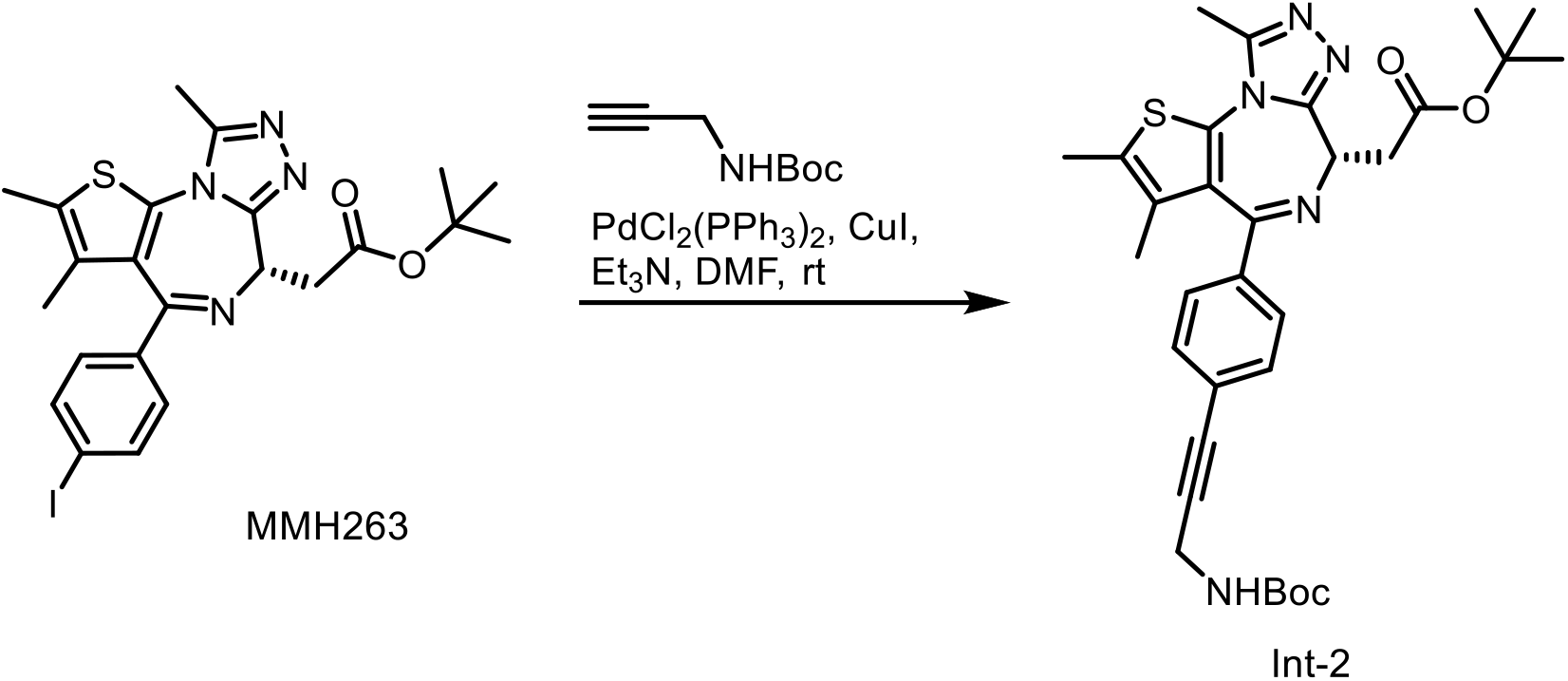

**tert-butyl (S)-2-(4-(4-(3-((tert-butoxycarbonyl)amino)prop-1-yn-1-yl)phenyl)-2,3,9- trimethyl-6H-thieno[3,2-f][1,2,4]triazolo[4,3-a][1,4]diazepin-6-yl)acetate (Int-2)**

To a solution of **MMH263** (40.0 mg, 0.073 mmol) and N-Boc-propargylamine (23.0 mg, 0.15 mmol) in Et_3_N/DMF (v/v=1/1, 1.2 mL) was added PdCl_2_(PPh_3_)_2_ (5.1 mg, 7.3 µmol), and CuI (2.8 mg, 14.6 µmol). The reaction mixture was stirred at rt for 3 hours. The reaction mixture was purified directly via prep HPLC (method 1) to give **Int-2** (33.1 mg, 79% yield) as a yellow oil. MS (ESI) for C_31_H_38_N_5_O_4_S [M+H]^+^: m/z calcd, 576.26; found, 576.18.

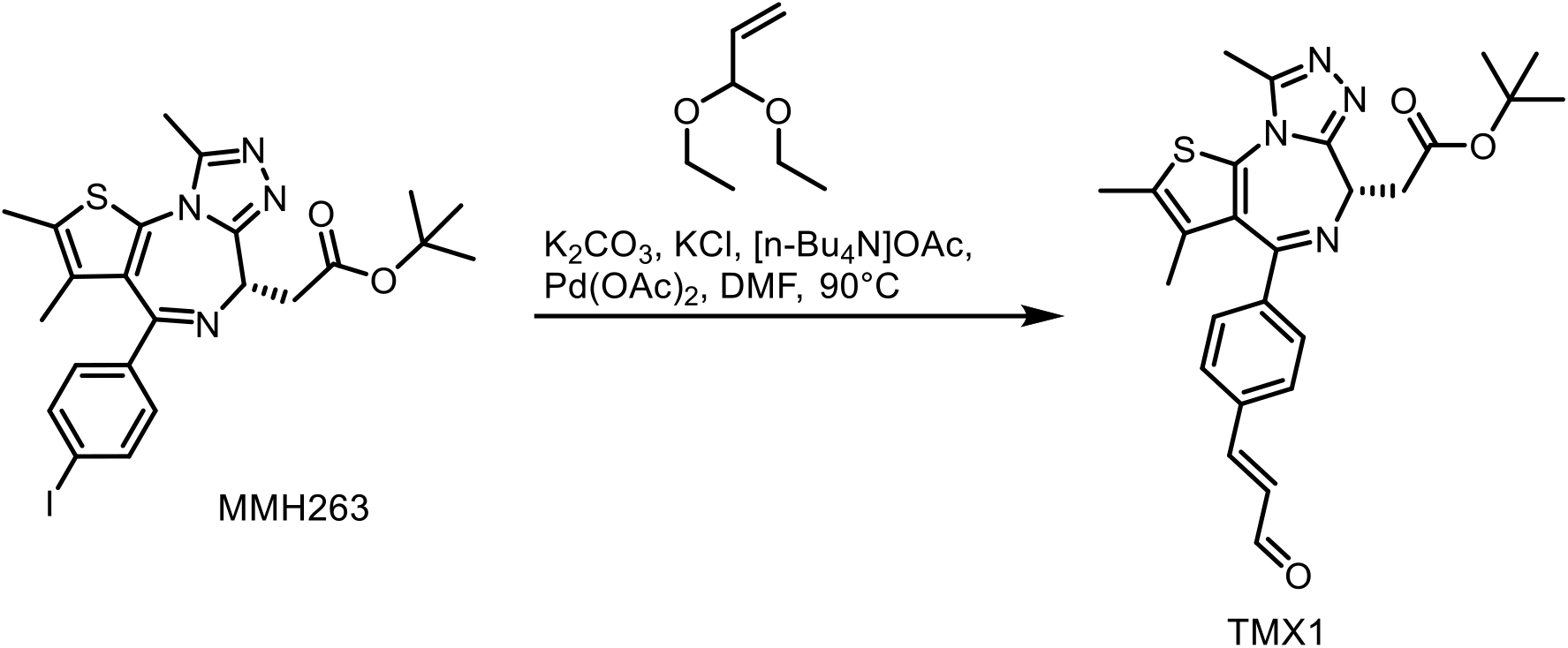

**tert-butyl (S,E)-2-(2,3,9-trimethyl-4-(4-(3-oxoprop-1-en-1-yl)phenyl)-6H-thieno[3,2- f][1,2,4]triazolo[4,3-a][1,4]diazepin-6-yl)acetate (TMX1)**

To a solution of **MMH263** (35.0 mg, 0.064 mmol) and 3,3-diethoxyprop-1-ene (13.0 mg, 0.096 mmol) in DMF (0.6 mL) was added K_2_CO_3_ (18.0 mg, 0.13 mmol), KCl (5.0 mg, 0.064 mmol), [n-Bu_4_N]OAc (39.0 mg, 0.13 mmol), and Pd(OAc)_2_ (4.0 mg, 0.019 mmol). The reaction mixture was stirred at 90 °C for 8 hours. The reaction mixture was purified directly via prep HPLC (method 1) to give **TMX1** (15.3 mg, 50% yield) as a yellow oil. ^1^H NMR (500 MHz, DMSO-*d*_6_) δ 9.69 (d, *J* = 7.5 Hz, 1H), 7.81 (d, *J* = 8.5 Hz, 2H), 7.76 (d, *J* = 16.5 Hz, 1H), 7.49 (d, *J* = 8.0 Hz, 2H), 6.91 (dd, *J* = 16.0, 7.5 Hz, 1H), 4.45 (dd, *J* = 8.0, 6.5 Hz, 1H), 3.40-3.29 (m, 2H), 2.61 (s, 3H), 2.42 (s, 3H), 1.64 (s, 3H), 1.44 (s, 9H). MS (ESI) for C_26_H_29_N_4_O_3_S [M+H]^+^: m/z calcd, 477.20; found, 477.50.

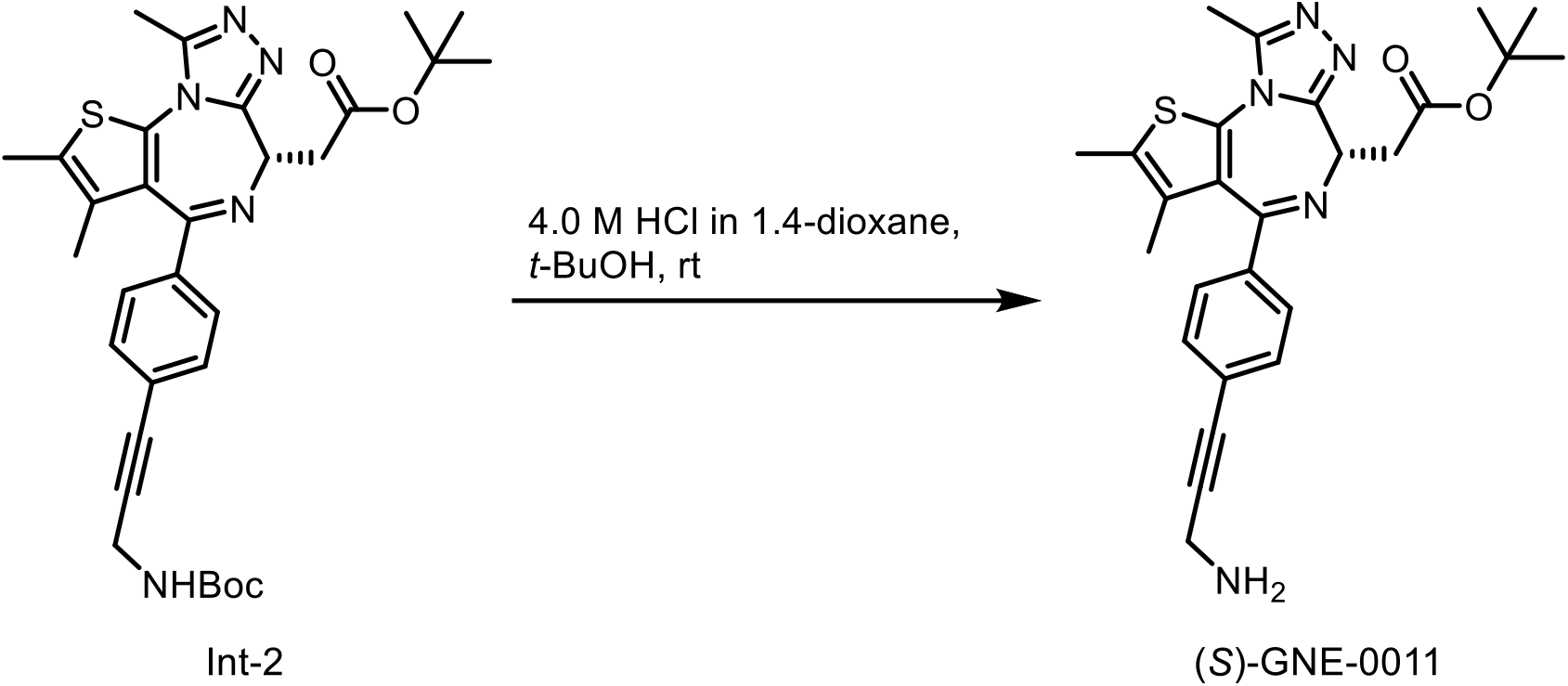

**tert-butyl (S)-2-(4-(4-(3-aminoprop-1-yn-1-yl)phenyl)-2,3,9-trimethyl-6H-thieno[3,2- f][1,2,4]triazolo[4,3-a][1,4]diazepin-6-yl)acetate ((S)-GNE-0011)**

A solution of **Int-2** (45.0 mg, 0.078 mmol) in *t*-BuOH/4.0 M HCl solution in 1,4-dixoane (v/v=1/1, 1.0 mL) was stirred at rt for 1.5 hours. The reaction mixture was purified directly via prep HPLC to give **(*S*)-GNE-0011 as a TFA salt** (20.0 mg, 41% yield) as a yellow oil. ^1^H NMR (500 MHz, DMSO-*d*_6_) δ 8.35 (brs, 3H), 7.53 (d, *J* = 8.5 Hz, 2H), 7.47 (d, *J* = 8.3 Hz, 2H), 4.45 (dd, *J* = 8.2, 6.2 Hz, 1H), 4.02 (q, *J* = 5.5 Hz, 2H), 3.39-3.28 (m, 2H), 2.61 (s, 3H), 2.42 (s, 3H), 1.63 (s, 3H), 1.43 (s, 9H). MS (ESI) for C_26_H_30_N_5_O_2_S [M+H]^+^: m/z calcd, 476.21; found, 476.31.

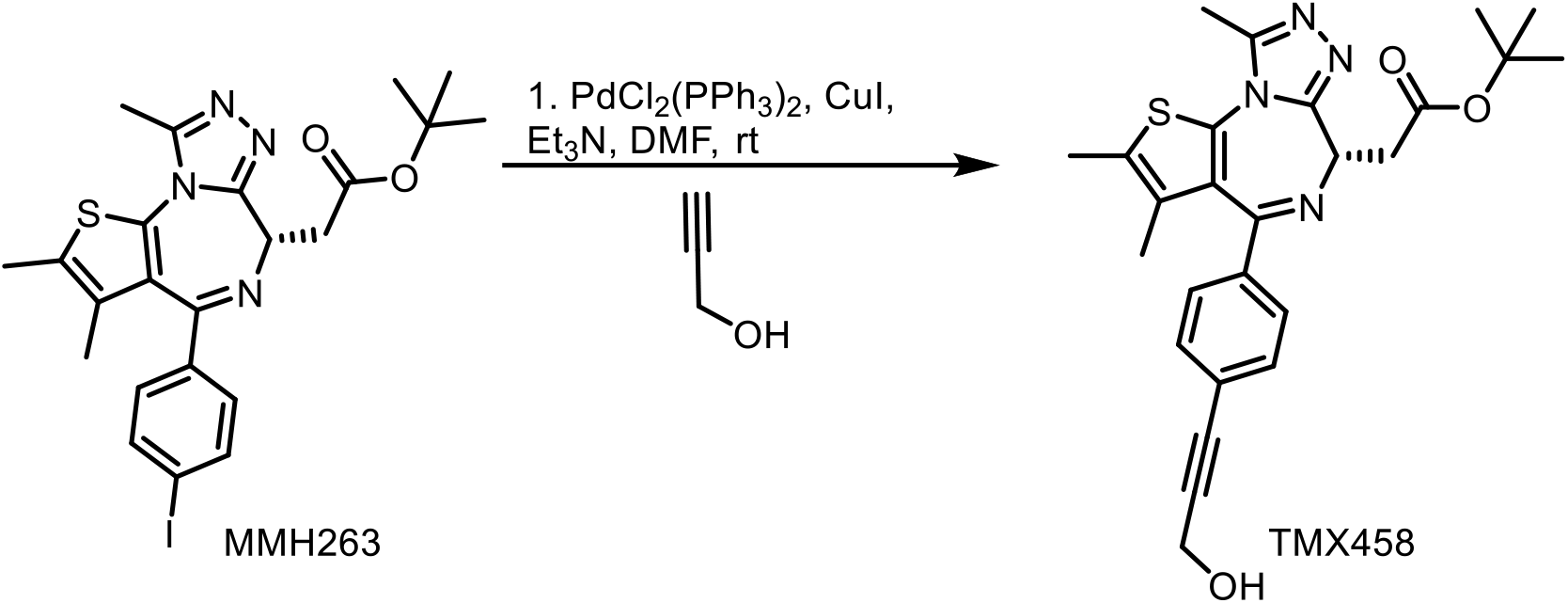

**tert-butyl (S)-2-(4-(4-(3-hydroxyprop-1-yn-1-yl)phenyl)-2,3,9-trimethyl-6H-thieno[3,2- f][1,2,4]triazolo[4,3-a][1,4]diazepin-6-yl)acetate (TMX458)**

To a solution of **MMH263** (20.0 mg, 0.037 mmol) and propargyl alcohol (2.04 mg, 0.037 mmol) in Et_3_N/DMF (v/v=0.5 µL/0.5 mL) was added PdCl_2_(PPh_3_)_2_ (5.0 mg, 7.3 µmol), and CuI (2.0 mg, 10.4 µmol). The reaction mixture was stirred at rt for 3 hours and subsequently purified by HPLC (method 1) to yield **TMX458** (10.0 mg, 57.6% yield). ^1^H NMR (500 MHz, DMSO) δ 7.48 (d, *J* = 8.6 Hz, 2H), 7.42 (d, *J* = 8.2 Hz, 2H), 4.44 (dd, *J* = 8.2, 6.2 Hz, 1H), 4.31 (s, 2H), 3.42 – 3.26 (m, 2H), 2.61 (s, 3H), 2.43 (s, 3H), 1.64 (s, 3H), 1.43 (s, 9H). MS (ESI) for C_26_H_28_N_4_O_3_S [M+H]^+^: m/z calcd, 477.60; found 477.31.

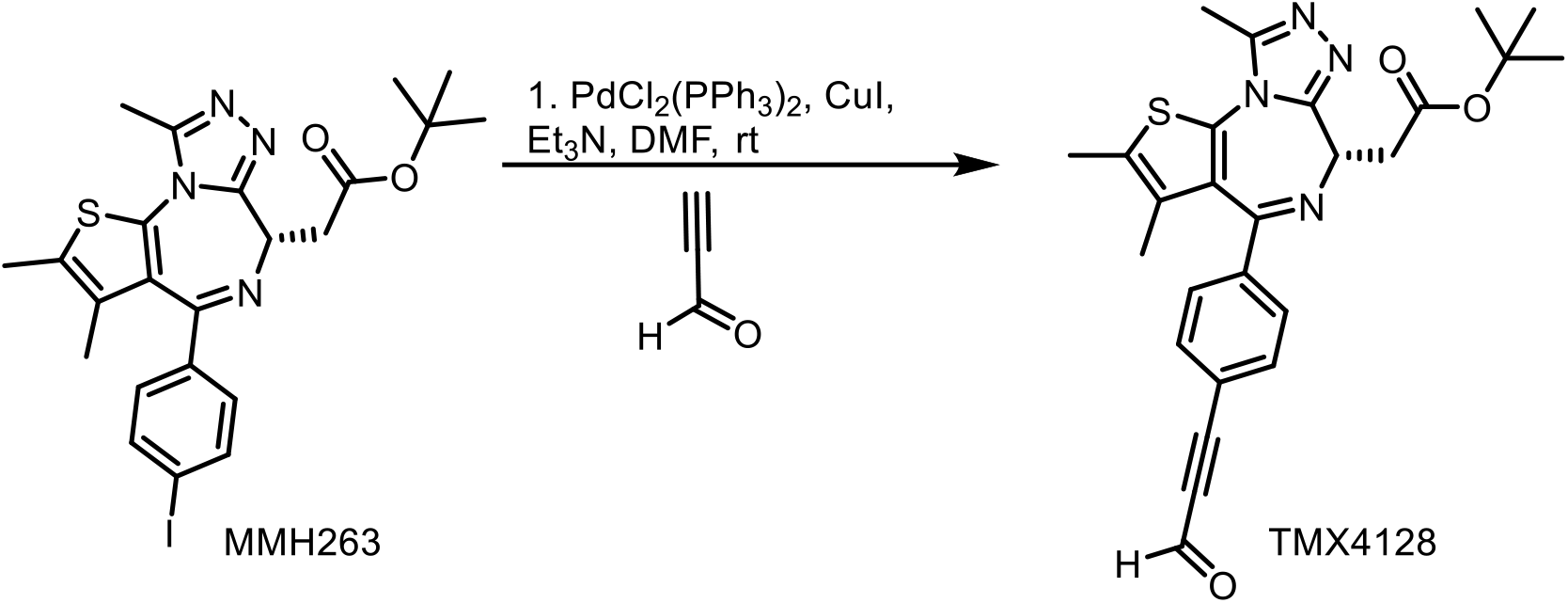

**tert-butyl (S)-2-(2,3,9-trimethyl-4-(4-(3-oxoprop-1-yn-1-yl)phenyl)-6H-thieno[3,2- f][1,2,4]triazolo[4,3-a][1,4]diazepin-6-yl)acetate (TMX4128)**

To a solution of **MMH263** (8.0 mg, 0.015 mmol) and propiolaldehyde (0.82 mg, 0.015 mmol) in Et_3_N/DMF (v/v=0.5 µL/0.5 mL) was added PdCl_2_(PPh_3_)_2_ (2.0 mg, 2.9 µmol), and CuI (0.8 mg, 4.2 µmol). The reaction mixture was stirred at rt for 3 hours and subsequently purified by HPLC (method 1) to yield **TMX4128** (3.6 mg, 50.5% yield). ^1^H NMR (500 MHz, DMSO) δ 9.45 (s, 1H), 7.76 (d, *J* = 8.2 Hz, 2H), 7.53 (d, *J* = 8.1 Hz, 2H), 4.47 (dd, *J* = 8.1, 6.2 Hz, 1H), 3.46 – 3.27 (m, 2H), 2.61 (s, 3H), 2.43 (s, 3H), 1.64 (s, 3H), 1.44 (s, 9H). MS (ESI) for C_26_H_26_N_4_O_3_S [M+H]^+^: m/z calcd, 475.58; found 475.26.

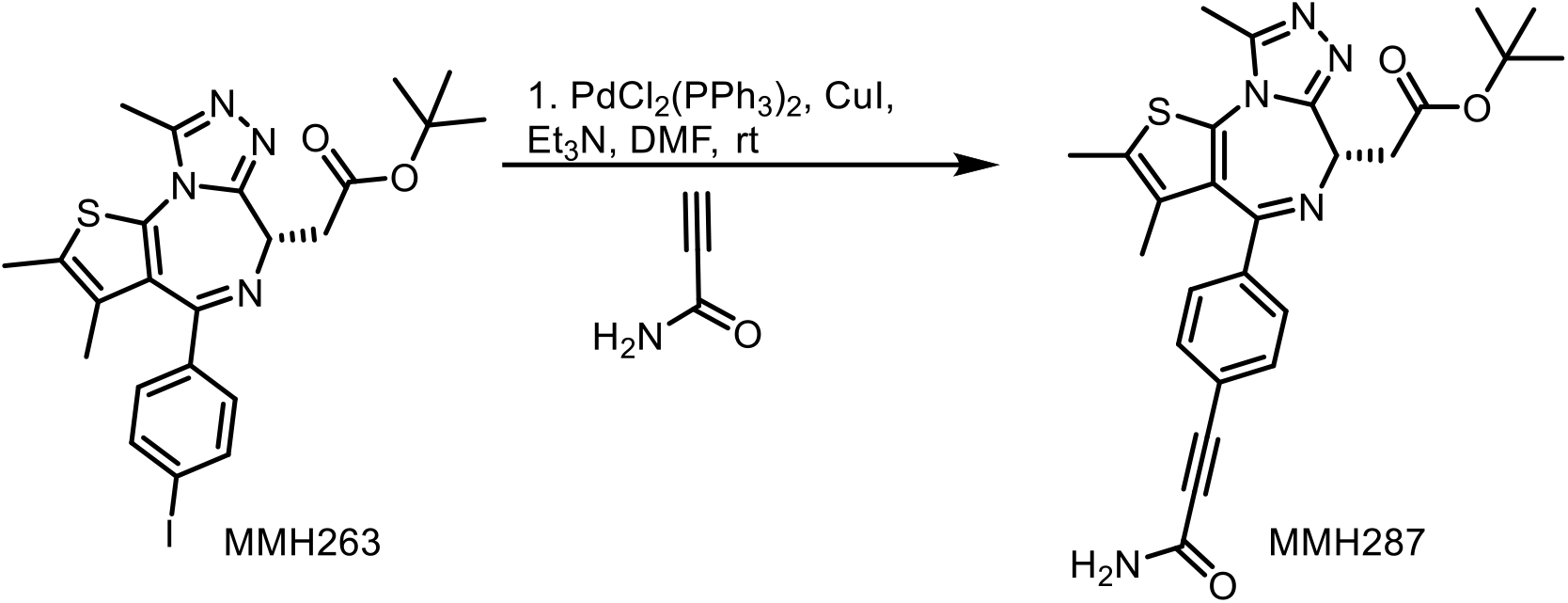

**tert-butyl (S)-2-(4-(4-(3-amino-3-oxoprop-1-yn-1-yl)phenyl)-2,3,9-trimethyl-6H-thieno[3,2- f][1,2,4]triazolo[4,3-a][1,4]diazepin-6-yl)acetate (MMH287)**

To a solution of **MMH263** (30.0 mg, 0.055 mmol) and propiolamide (7.56 mg, 0.11 mmol) in Et_3_N/DMF (v/v=1/1, 1.2 mL) was added PdCl_2_(PPh_3_)_2_ (15.4 mg, 0.022 mmol), and CuI (6.25 mg, 0.033 mmol). The reaction mixture was stirred at rt for 4 hours. The reaction mixture was purified twice via prep HPLC (method 2) to give **MMH287** (6 mg, 23% yield). ^1^H NMR (500 MHz, DMSO) δ 8.19 (br, 1H), 7.72 (br, 1H), 7.62 (d, *J* = 8.19 Hz, 2H), 7.49 (d, *J* = 8.06 Hz, 2H), 4.45 (t, J = 7.2 Hz, 1H), 3.40-3.29 (m, 2H), 2.61 (s, 3H), 2.43 (s, 3H), 1.63 (s, 3H), 1.43 (s, 9H). MS (ESI) for C_26_H_27_N_5_O_3_S [M+H]^+^: m/z calcd, 490.60; found 490.37.

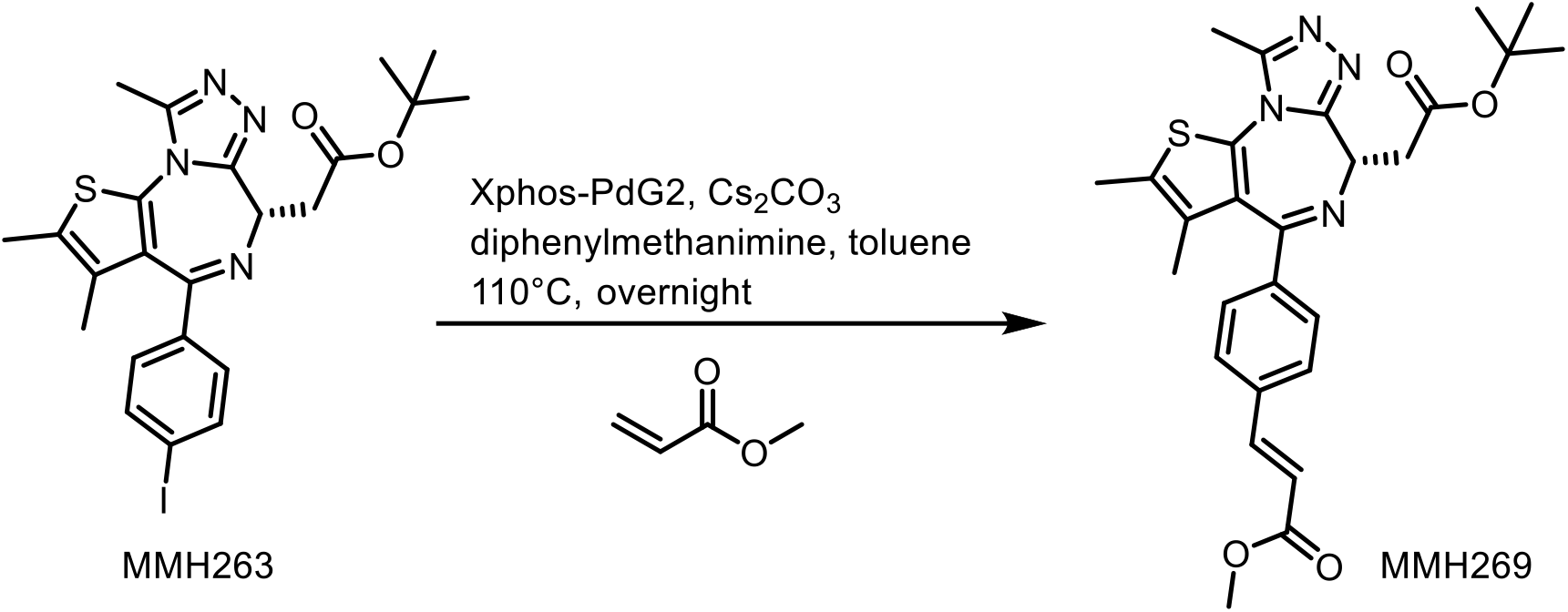

**Methyl (S,E)-3-(4-(6-(2-(tert-butoxy)-2-oxoethyl)-2,3,9-trimethyl-6H-thieno[3,2- f][1,2,4]triazolo[4,3-a][1,4]diazepin-4-yl)phenyl)acrylate (MMH269)**

To a solution of **MMH263** (20.9 mg, 0.03 mmol) in 3 mL of toluene, was added cesium carbonate (37.2 mg, 0.114 mmol), Xphos Pd G2 (5.08 mg, 0.0065 mmol), and methyl acrylate (3.6 mg, 0.0418). The reaction was heated to 110 °C and left to stir overnight. The reaction was filtered through celite and evaporated under reduced pressure. The mixture was purified by HPLC (method 2) to yield **MMH269** (8.91 mg, 46.3% yield). ^1^H NMR (500 MHz, DMSO) δ 7.79 (d, *J* = 8.3 Hz, 2H), 7.68 (d, *J* = 16.1 Hz, 1H), 7.46 (d, *J* = 8.0 Hz, 2H), 6.72 (d, *J* = 16.1 Hz, 1H), 4.45 (ddd, *J* = 7.7, 6.2, 1.4 Hz, 1H), 3.74 (s, 3H), 3.39 – 3.29 (m, 2H), 2.62 (s, 3H), 2.43 (d, *J* = 0.9 Hz, 3H), 1.64 (d, *J* = 0.9 Hz, 3H), 1.44 (s, 9H). MS (ESI) for C_27_H_30_N_4_O_4_S [M+H]^+^: m/z calcd, 507.63; found 507.31.

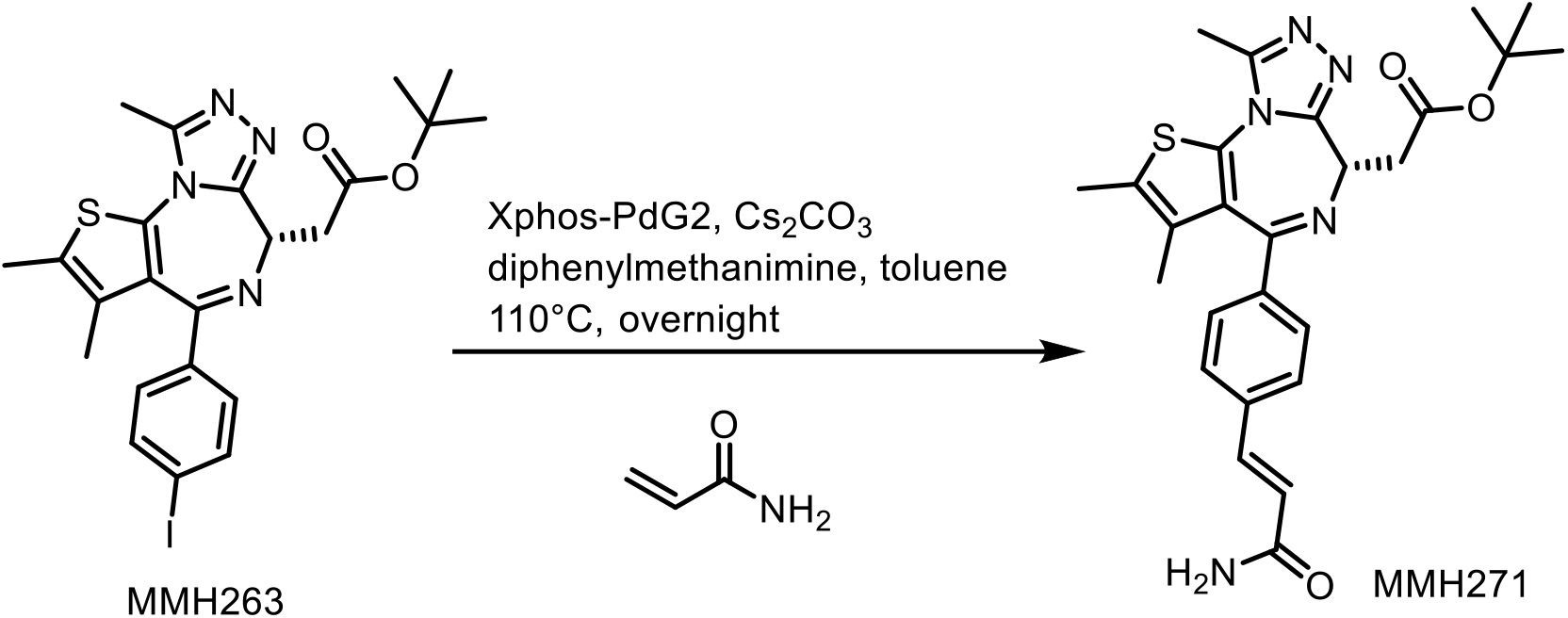

**tert-butyl (S,E)-2-(4-(4-(3-amino-3-oxoprop-1-en-1-yl)phenyl)-2,3,9-trimethyl-6H-thieno[3,2-f][1,2,4]triazolo[4,3-a][1,4]diazepin-6-yl)acetate (MMH271)**

To a solution of **MMH263** (12.0 mg, 0.022 mmol) in 3 mL of toluene, was added cesium carbonate (21.4 mg, 0.066 mmol), Xphos Pd G2 (4.73 mg, 0.006 mmol), and acrylamide (3.34 mg, 0.047). The reaction was heated to 110 °C and left to stir overnight. The reaction was filtered through celite and evaporated under reduced pressure. The mixture was purified by HPLC (method 2) to yield **MMH271** (3.48 mg, 32.2% yield).^1^H NMR (500 MHz, MeOD) δ 7.64 (d, *J* = 8.5 Hz, 2H), 7.58 (d, *J* = 15.9 Hz, 1H), 7.52 (d, *J* = 8.4 Hz, 2H), 6.73 (d, *J* = 15.8 Hz, 1H), 4.61 (dd, *J* = 8.7, 5.9 Hz, 1H), 3.54 – 3.40 (m, 2H), 2.74 (s, 3H), 2.49 (d, *J* = 0.8 Hz, 3H), 1.74 (d, *J* = 0.9 Hz, 3H), 1.53 (s, 9H). MS (ESI) for C_26_H_29_N_5_O_3_S [M+H]^+^: m/z calcd, 492.61; found 492.30.

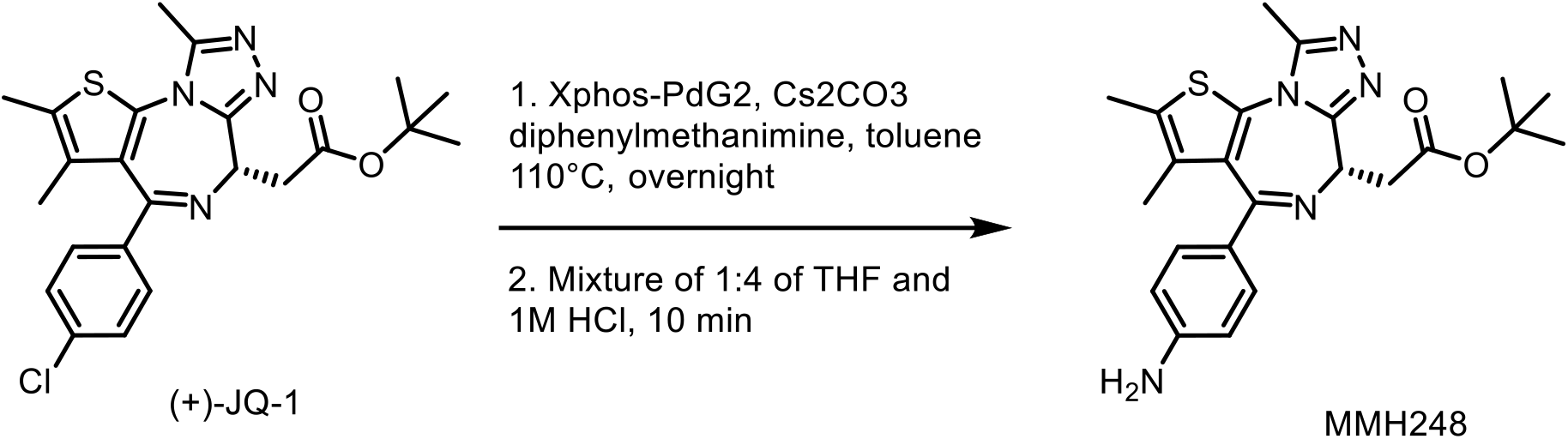

**tert-butyl (S)-2-(4-(4-aminophenyl)-2,3,9-trimethyl-6H-thieno[3,2-f][1,2,4]triazolo[4,3- a][1,4]diazepin-6-yl)acetate (MMH248)**

**MMH248** was prepared according to the procedure previously described by Dragovich *et al*. with slight modifications.^1^ Briefly, a mixture of JQ1 (1.40 g, 3.06 mmol), diphenylmethanimine (1.055 g, 5.82 mmol), Xphos-PdG2 (0.241 g, 0.306 mmol), and cesium carbonate (2.994 g, 9.19 mmol) in toluene (20 mL) was heated at 110 °C and left to stir overnight. The next day, the mixture was cooled to room temperature, filtered, and the filtrate was concentrated under vacuum. The residue was purified by flash column chromatography (EtOAc:hexane) and the benzophenone-imine protected product identity was confirmed by LCMS. MS (ESI) for C_36_H_36_N_5_O_2_S [M+H]^+^: m/z calcd, 602.77; found 602.40. The imine was hydrolyzed by the addition of 20 mL THF, and adding 5 mL of 1 M HCl, followed by stirring for approximately 10 minutes at room temperature. The mixture was dried under vacuum to yield **MMH248** (927 mg, 2.12 mmol, overall yield 69.2 %). ^1^H NMR (500 MHz, DMSO) δ 7.12 (d, *J* = 8.1 Hz, 2H), 6.52 (d, *J* = 8.9 Hz, 2H), 2H), 5.62 (s, 2H), 4.30 (dd, *J* = 8.3, 6.3 Hz, 1H), 3.30 – 3.21 (m, 2H), 2.58 (s, 3H), 2.43 (d, *J* = 1.0 Hz, 3H), 1.74 (d, *J* = 0.9 Hz, 2H), 1.42 (s, 9H). MS (ESI) for C_23_H_27_N_5_O_2_S [M+H]^+^: m/z calcd, 438.57; found 438.34. MS (ESI) for C_23_H_28_N_5_O_2_S [M+H]^+^: m/z calcd, 438.56; found 438.35.

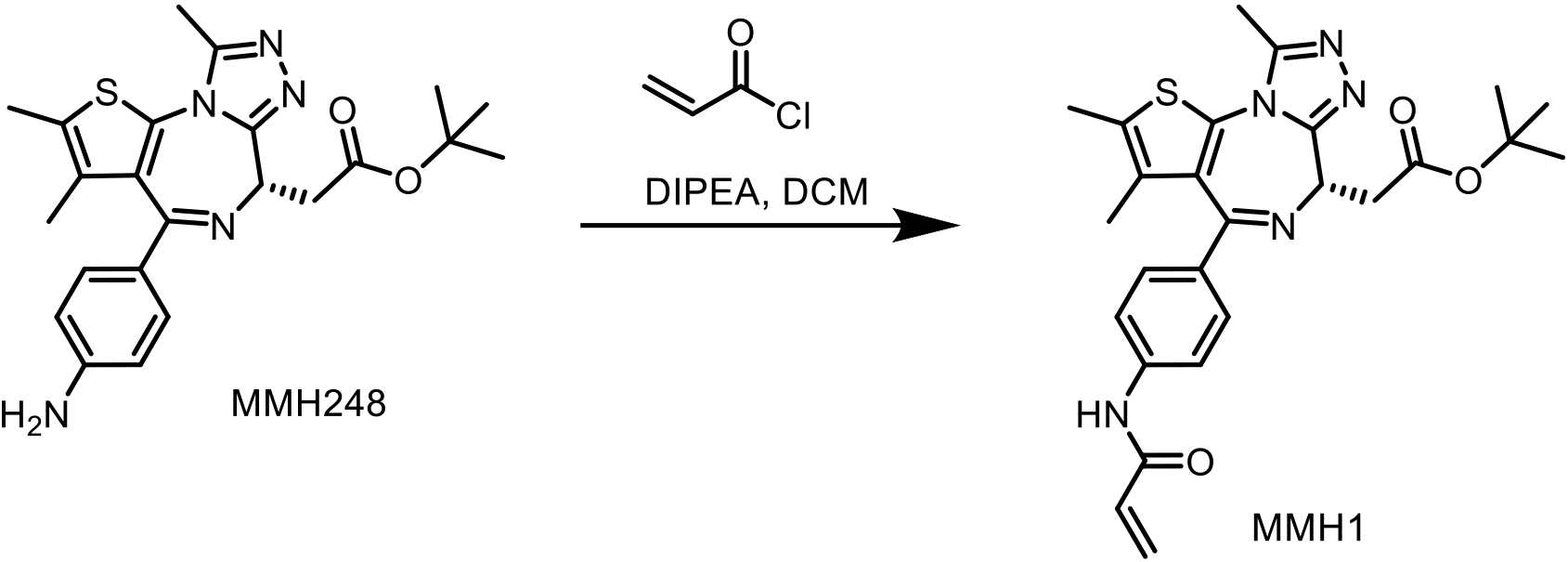

**tert-butyl (S)-2-(4-(4-acrylamidophenyl)-2,3,9-trimethyl-6H-thieno[3,2-f][1,2,4]triazolo[4,3- a][1,4]diazepin-6-yl)acetate (MMH1)**

**MMH1** was prepared by adding acryloyl chloride (4.5 µL, 0.055 mmol) to a mixture of **MMH248** (20.0 mg, 0.046 mmol) and DIPEA (23.4 µL, 0.137 mmol) stirred in 2 mL dichloromethane. The reaction was stirred for 5 minutes and monitored by LCMS, and subsequently quenched with methanol followed by drying under vacuum. The mixture was reconstituted in methanol and purified by prep HPLC (method 2) to give **MMH1** (1.40 mg, 0.00285 mmol, 6.23% yield). ^1^H NMR (500 MHz, DMSO) δ 10.35 (s, 1H), 7.71 (d, *J* = 9.0 Hz, 2H), 7.40 (d, *J* = 8.4 Hz, 2H), 6.44 (dd, *J* = 16.9, 10.1 Hz, 1H), 6.28 (dd, *J* = 16.9, 2.0 Hz, 1H), 5.78 (dd, *J* = 10.1, 2.0 Hz, 1H), 4.39 (dd, *J* = 8.2, 6.3 Hz, 1H), 3.35 – 3.25 (m, 2H), 2.61 (s, 3H), 2.43 (s, 3H), 1.67 (s, 3H), 1.43 (s, 9H). MS (ESI) for C_26_H_30_N_5_O_3_S [M+H]^+^: m/z calcd, 492.61; found 492.35.

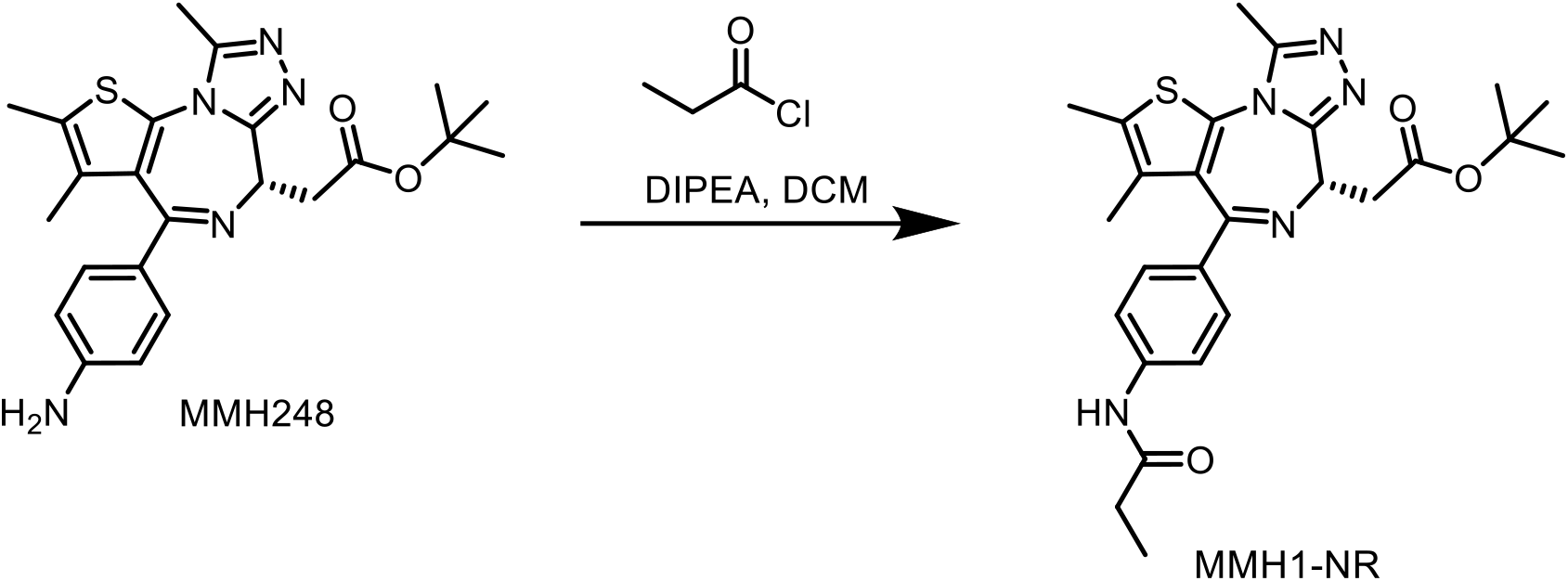

**tert-butyl (S)-2-(2,3,9-trimethyl-4-(4-propionamidophenyl)-6H-thieno[3,2- f][1,2,4]triazolo[4,3-a][1,4]diazepin-6-yl)acetate (MMH1-NR)**

Propionyl chloride (2.5 µL, 0.0282 mmol) was added to a mixture containing **MMH248** (10.3 mg, 0.0235 mmol) and DIPEA (8.05 µL, 0.0471 mmol) in 2 mL dichloromethane. The reaction was left to stir for 1 h, before evaporation under vacuum. The mixture was redissolved in methanol and purified by prep HPLC (method 2) to give **MMH1-NR** (7.75mg, 0.0157 mmol, 66.8% yield). ^1^H NMR (500 MHz, DMSO) δ 10.08 (s, 1H), 7.64 (d, *J* = 9.1 Hz, 2H), 7.36 (d, *J* = 8.4 Hz, 2H), 4.39 (dd, *J* = 8.2, 6.3 Hz, 1H), 3.38 – 3.24 (m, 2H), 2.61 (s, 3H), 2.43 (d, *J* = 0.9 Hz, 3H), 2.34 (q, *J* = 7.6 Hz, 2H), 1.66 (d, *J* = 1.0 Hz, 3H), 1.43 (s, 9H), 1.08 (t, *J* = 7.6 Hz, 3H). MS (ESI) for C_26_H_32_N_5_O_3_S [M+H]^+^: m/z calcd, 494.63; found 494.42.

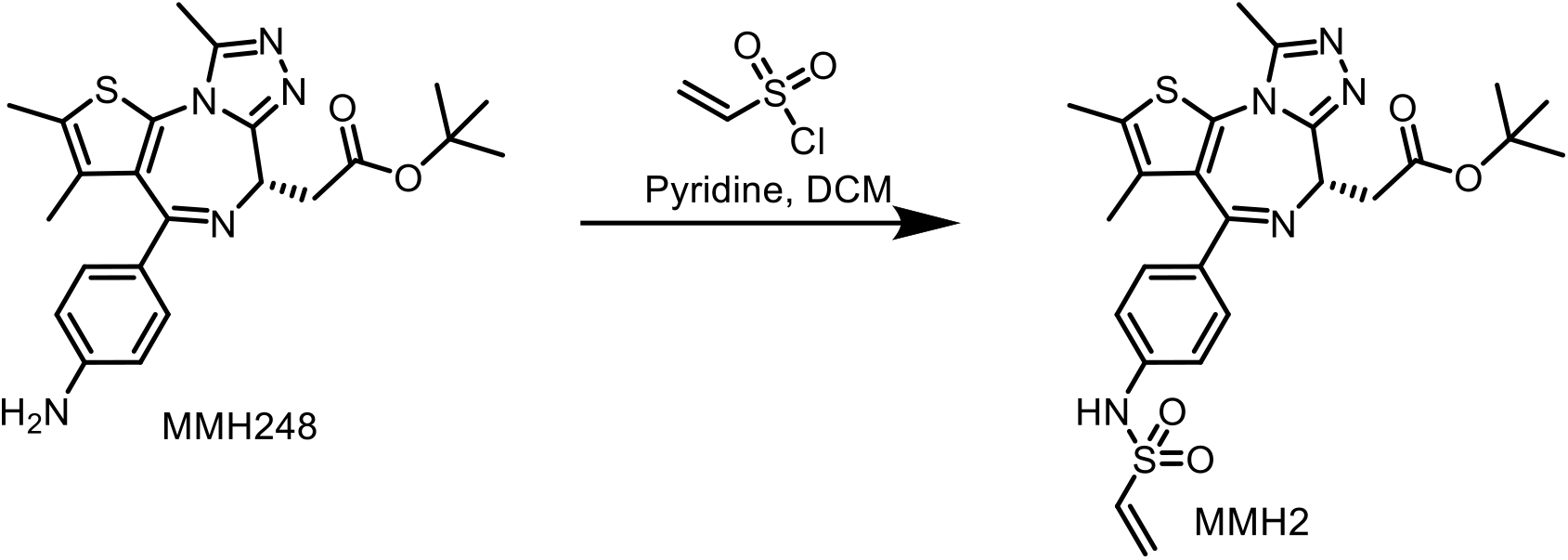

**tert-butyl (S)-2-(2,3,9-trimethyl-4-(4-(vinylsulfonamido)phenyl)-6H-thieno[3,2- f][1,2,4]triazolo[4,3-a][1,4]diazepin-6-yl)acetate (MMH2)**

Ethenesulfonyl chloride (23.9 µL, 0.263 mmol) was added to a mixture of **MMH248** (9.59 mg, 0.0219 mmol) and pyridine (35.5 µL, 0.438 mmol). The reaction was instantaneous as shown by LCMS. The reaction was washed once with 1 M HCl (1 mL), then the remaining DCM mixture was evaporated. The mixture was reconstituted in methanol to quench excess electrophile, and was subsequently purified by prep HPLC (method 2) to give **MMH2** (5.94 mg, 0.0113 mmol, 51.4% yield). ^1^H NMR (500 MHz, DMSO-*d*_6_) δ 10.32 (s, 1H), 7.36 (d, *J* = 8.4 Hz, 2H), 7.17 (d, *J* = 8.85 2H), 6.79 (dd, *J* = 16.4, 10.0 Hz, 1H), 6.14 (d, *J* = 16.4 Hz, 1H), 6.05 (d, *J* = 9.9 Hz, 1H), 4.39 (dd, *J* = 8.3, 6.3 Hz, 1H), 3.38 – 3.24 (m, 2H), 2.60 (s, 3H), 2.42 (s, 3H), 1.65 (s, 1H), 1.43 (s, 9H). MS (ESI) for C_25_H_30_N_5_O_4_S_2_ [M+H]^+^: m/z calcd, 528.67; found 528.21.

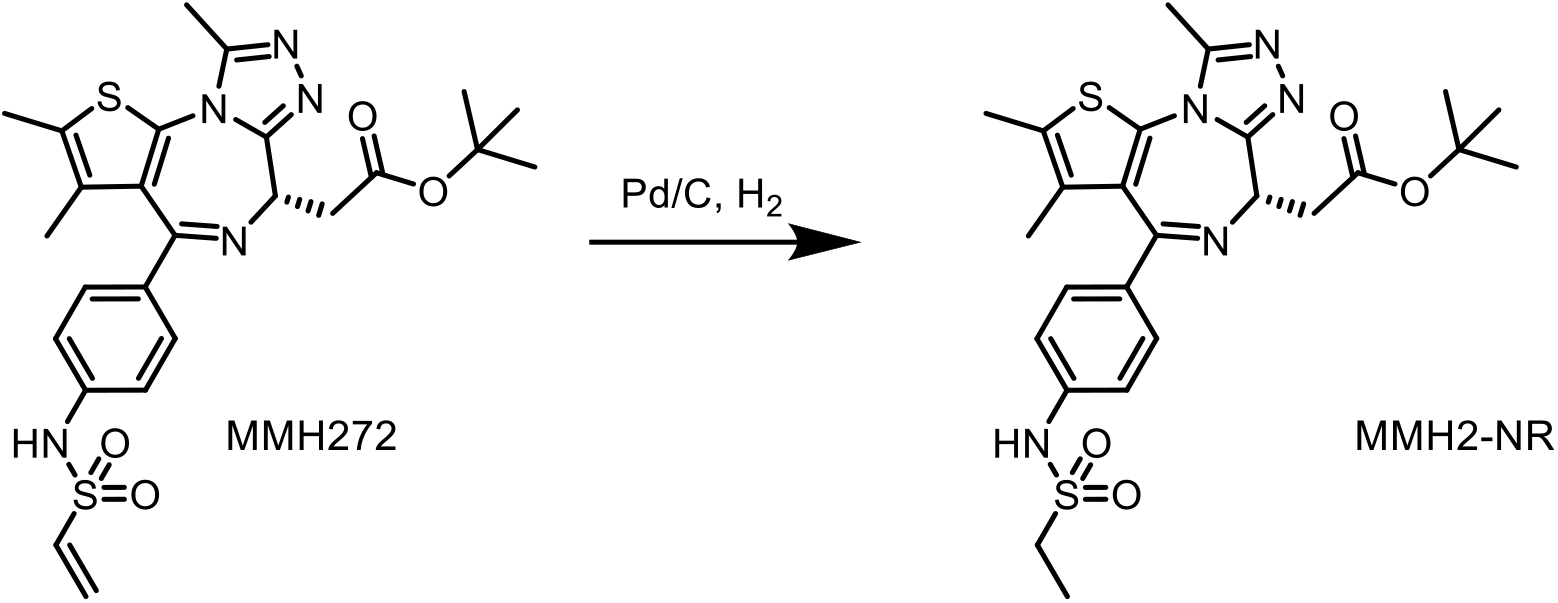

**tert-butyl (S)-2-(4-(4-(ethylsulfonamido)phenyl)-2,3,9-trimethyl-6H-thieno[3,2- f][1,2,4]triazolo[4,3-a][1,4]diazepin-6-yl)acetate (MMH2-NR)**

To a solution of 4 mL of ethanol, was added **MMH2** (2.26 mg, 0.00428 mmol) and 10% palladium on carbon (10.5 mg, 0.00985 mmol), followed by purging with hydrogen gas via a balloon for 10 min. The reaction was subsequently stirred under hydrogen for another 10 min, flushed through celite, then evaporated and lyophilized to quantitatively yield **MMH2-NR** (2.26 mg, 0.00427 mmol, 99.7% yield). ^1^H NMR (500 MHz, DMSO) δ 10.09 (s, 1H), 7.37 (d, *J* = 8.5 Hz, 2H), 7.23 (d, *J* = 9.0 Hz, 2H), 4.39 (dd, *J* = 8.3, 6.2 Hz, 1H), 3.33 – 3.24 (m, 2H), 3.12 (q, *J* = 7.3 Hz, 2H), 2.59 (s, 3H), 2.42 (s, 3H), 1.67 (s, 3H), 1.43 (s, 9H), 1.16 (t, *J* = 7.3 Hz, 3H). MS (ESI) for C_25_H_32_N_5_O_4_S_2_ [M+H]^+^: m/z calcd, 530.68; found 530.33.

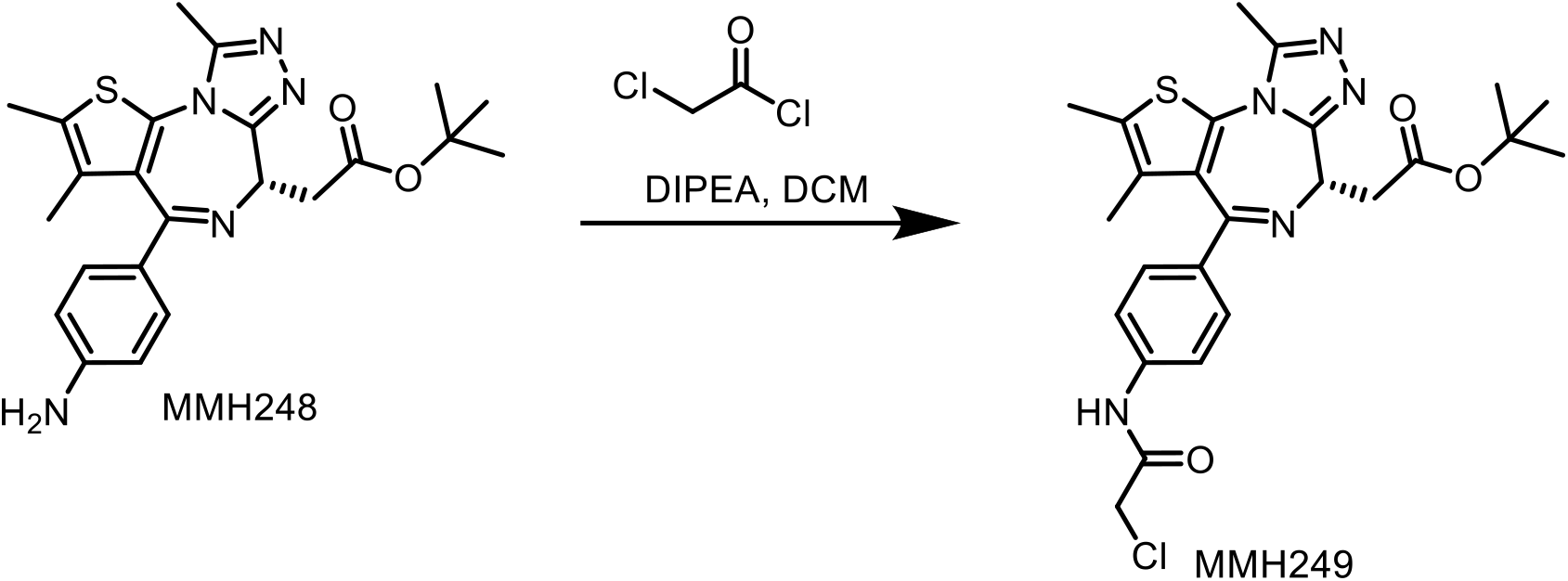

**tert-butyl (S)-2-(4-(4-(2-chloroacetamido)phenyl)-2,3,9-trimethyl-6H-thieno[3,2- f][1,2,4]triazolo[4,3-a][1,4]diazepin-6-yl)acetate (MMH249)**

To a solution of **MMH248** (40 mg, 0.0914 mmol), chloroacetic acid (21.6 mg, 0.229 mmol), and DIPEA (47 µL, 0.274 mmol) in 3 mL DMF, was added HATU (104 mg, 0.274 mmol). The reaction was purified by HPLC (method 2) to yield **MMH249** which was again purified by column chromatography using DCM and methanol (14 mg, 29.8 % yield). ^1^H NMR (500 MHz, DMSO) δ 10.51 (s, 1H), 7.64 (d, *J* = 8.7 Hz, 2H), 7.40 (d, *J* = 8.5 Hz, 2H), 4.40 (dd, *J* = 8.2, 6.3 Hz, 1H), 4.27 (s, 2H), 3.33 – 3.27 (m, 2H), 2.60 (s, 3H), 2.43 (d, *J* = 0.9 Hz, 3H), 1.66 (d, *J* = 0.9 Hz, 3H), 1.43 (s, 9H). MS (ESI) for C_25_H_28_ClN_5_O_3_S [M+H]^+^: m/z calcd, 514.16 and 516.16 for ^35^Cl and ^37^Cl respectively; found 514.26 and 516.21 respectively.

## Spectral Characterization of Final Compounds (_1-_H NMR, LC, and MS Spectra)

### (*S*)-GNE011 (500 MHz ^1^H NMR in DMSO-*d*_6_)

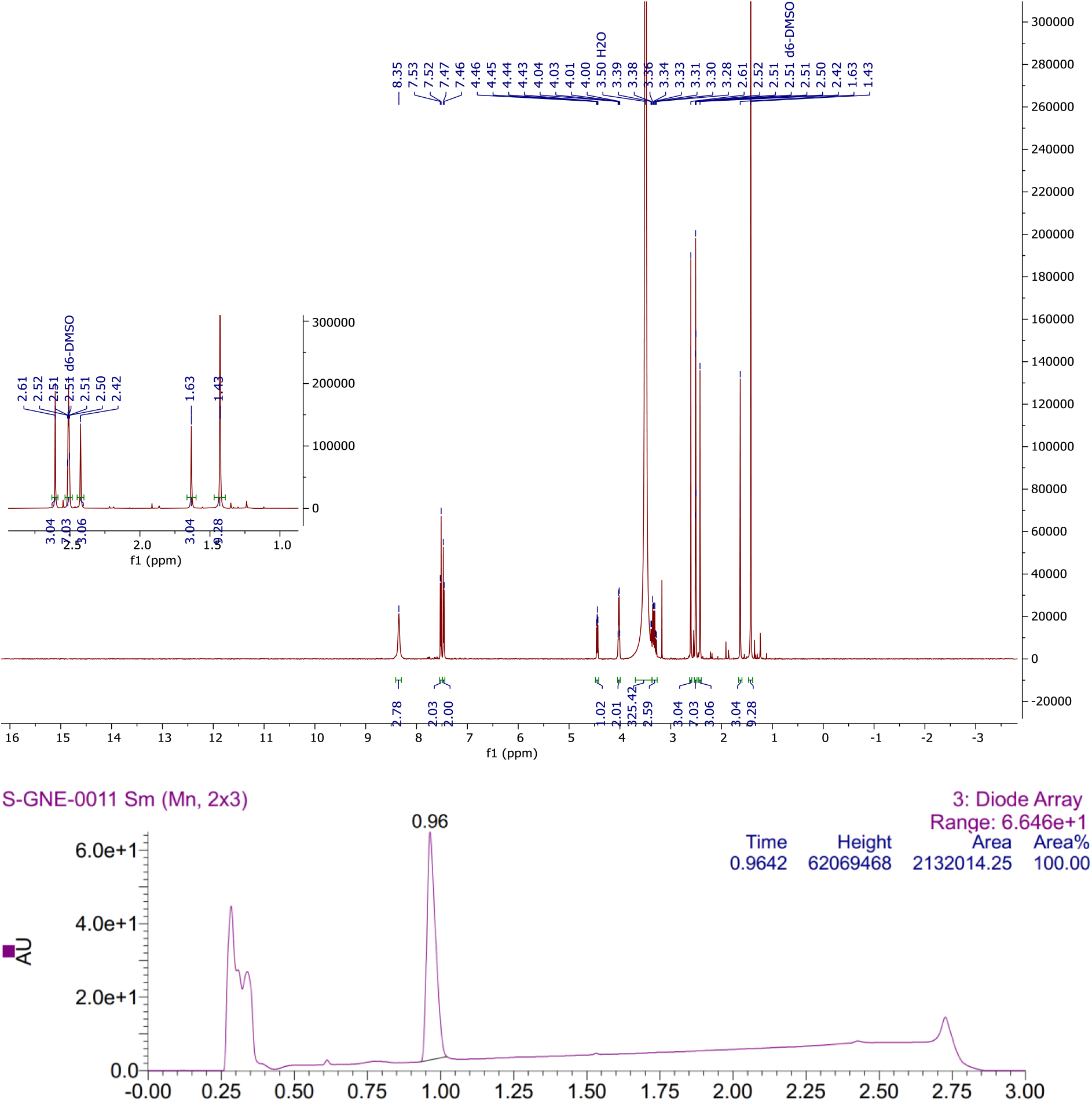

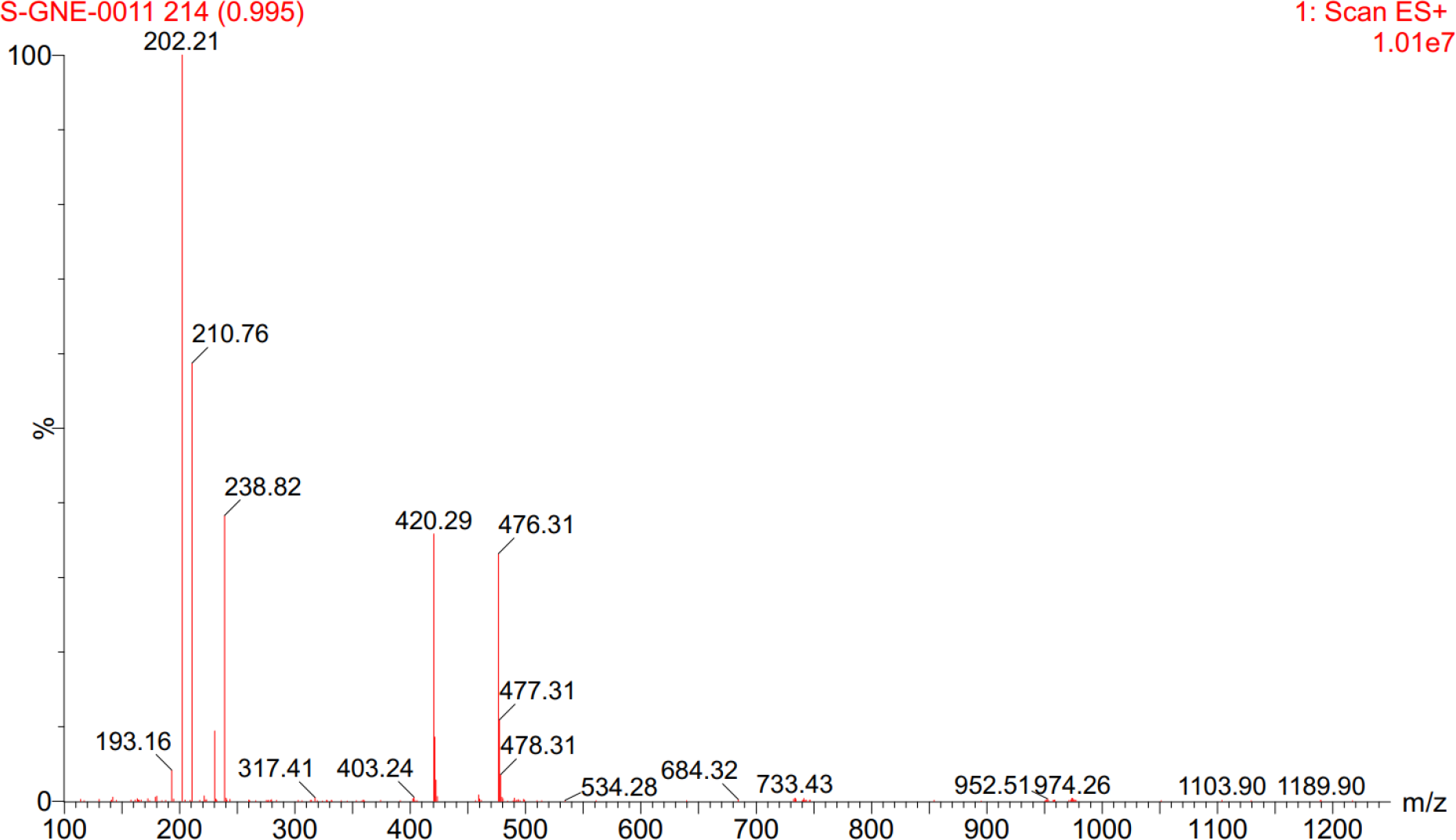

### TMX1 (500 MHz ^1^H NMR in DMSO-*d*_6_)

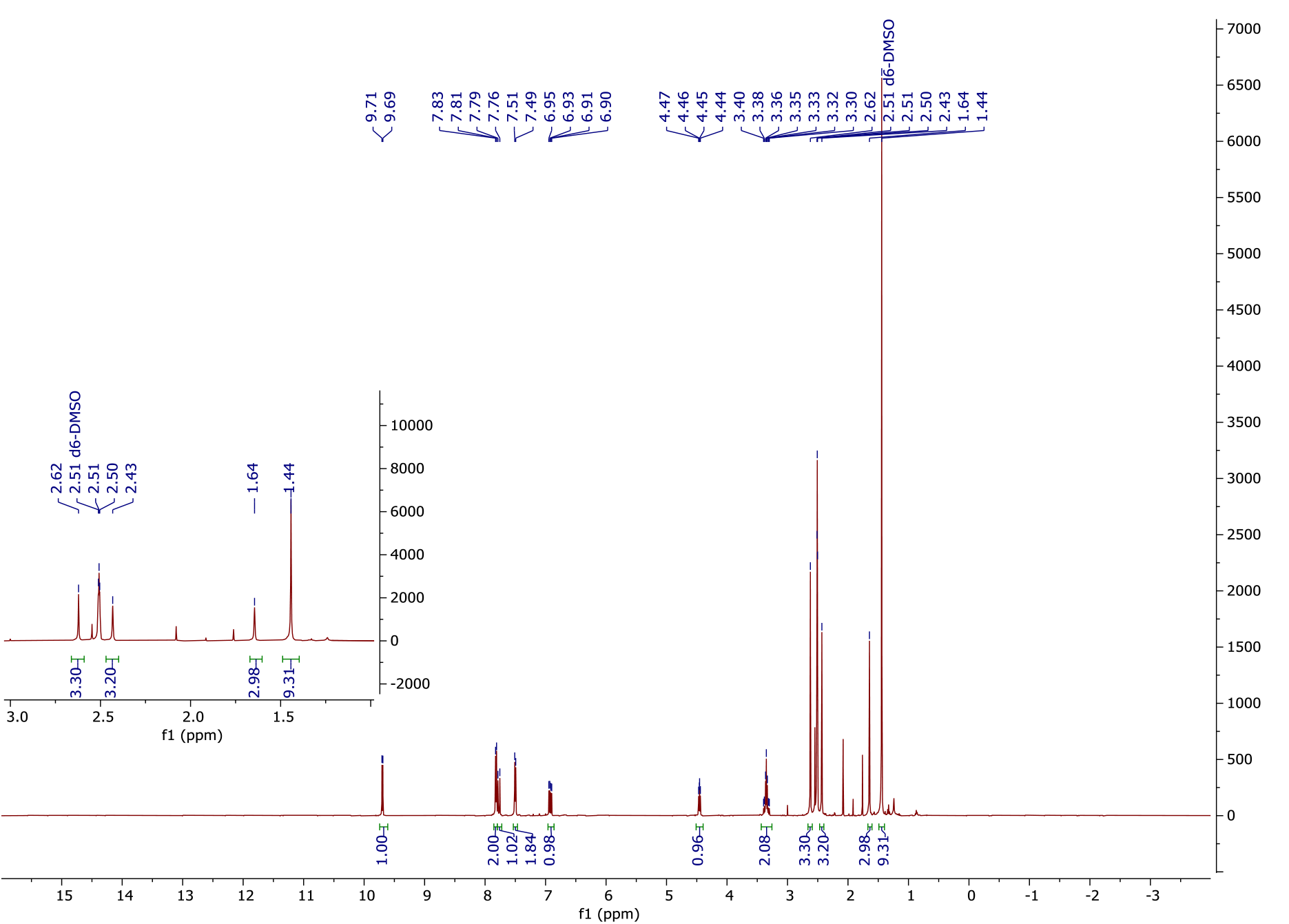

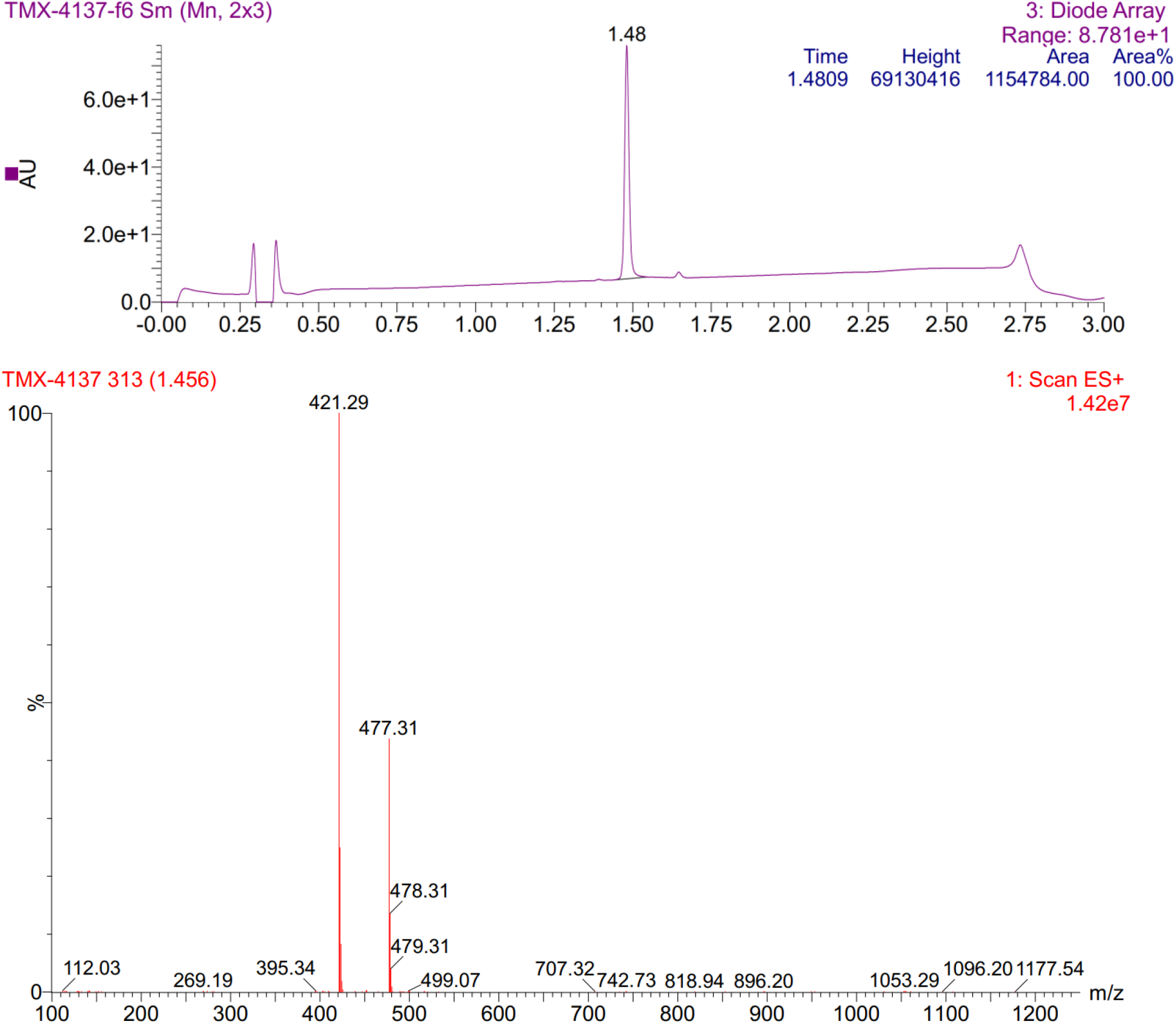

### MMH1 (500 MHz ^1^H NMR in DMSO-*d*_6_)

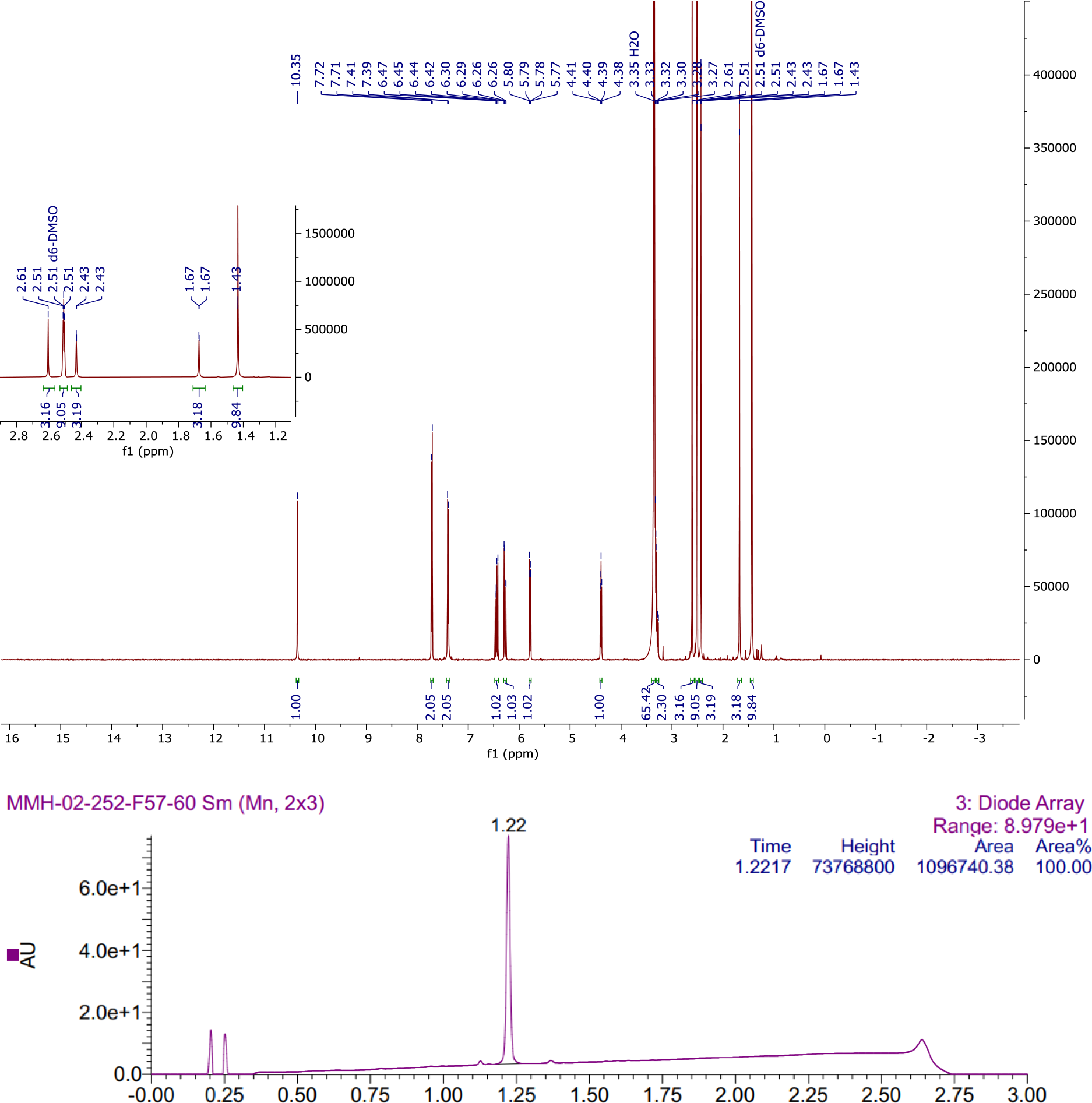

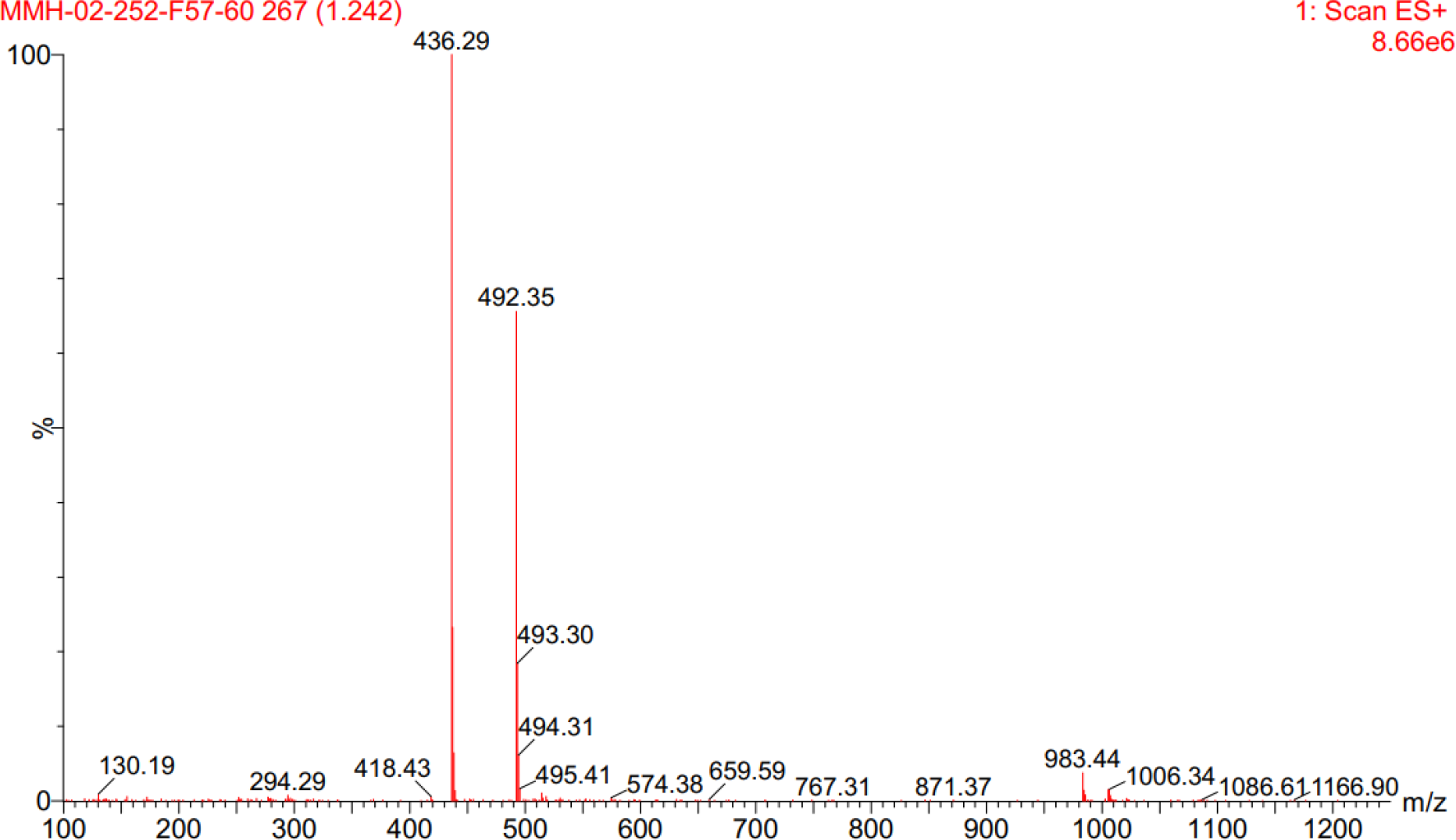

### MMH1-NR (500 MHz ^1^H NMR in DMSO-*d*_6_)

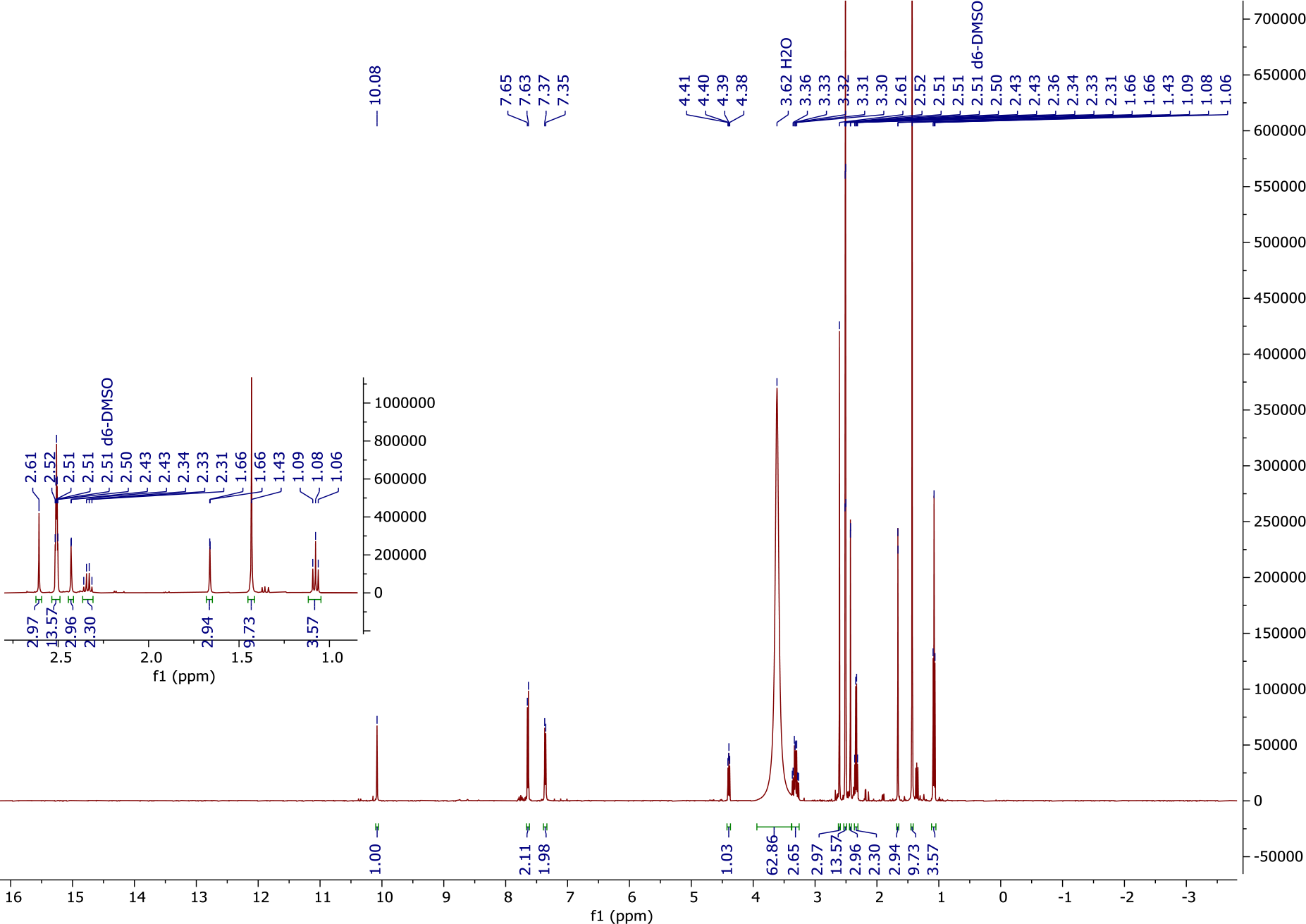

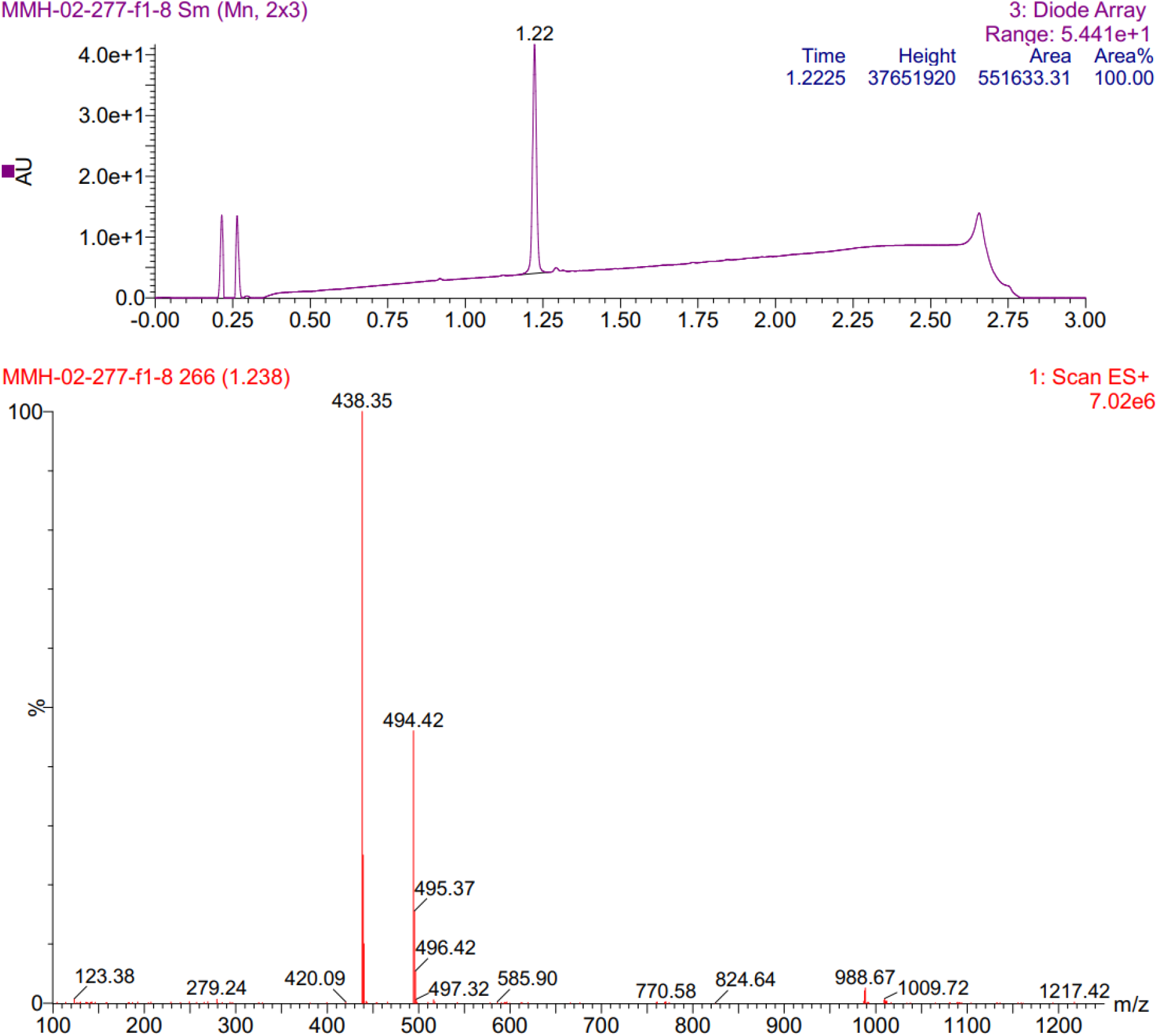

### MMH2 (500 MHz ^1^H NMR in DMSO-*d*_6_)

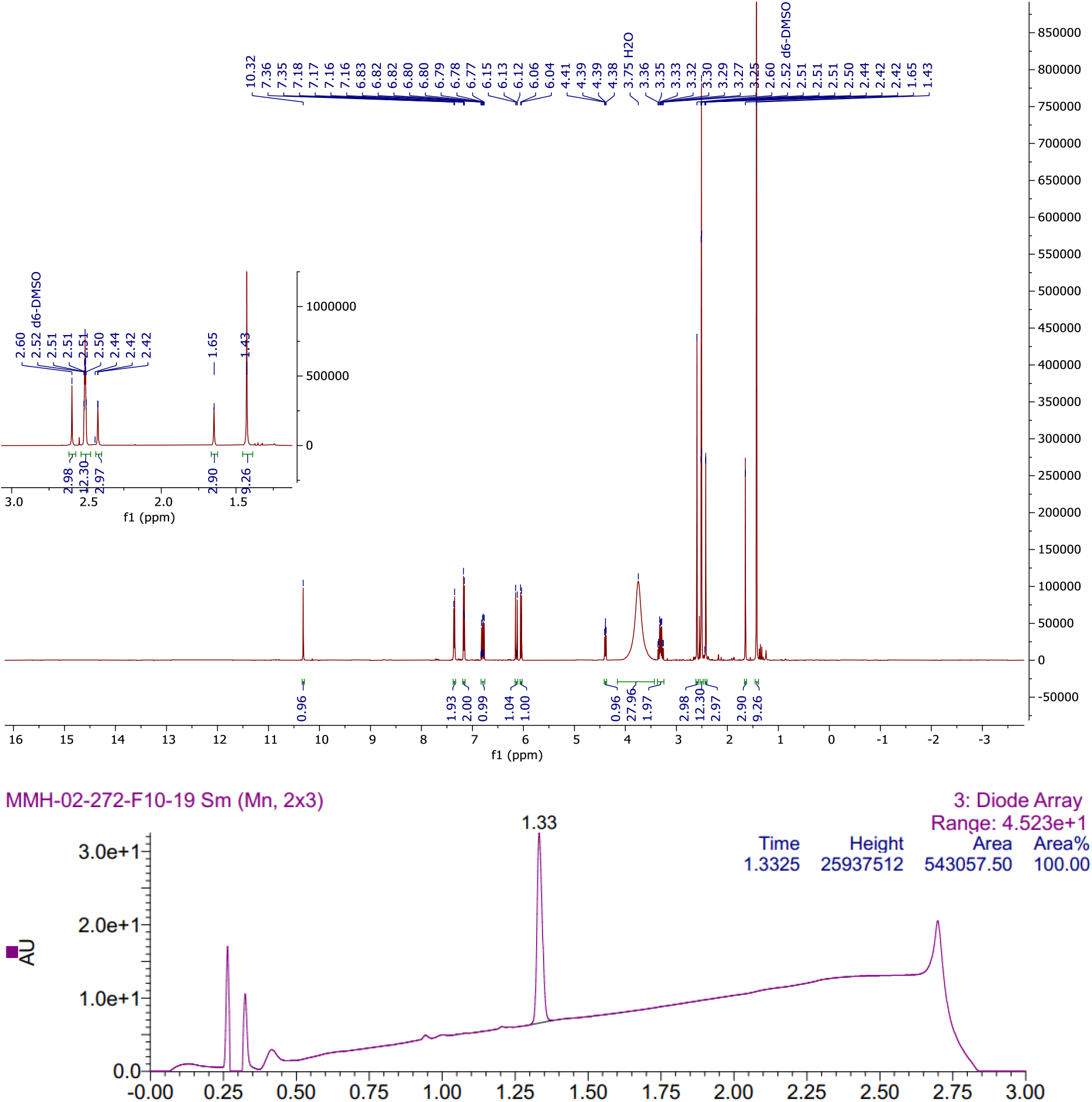

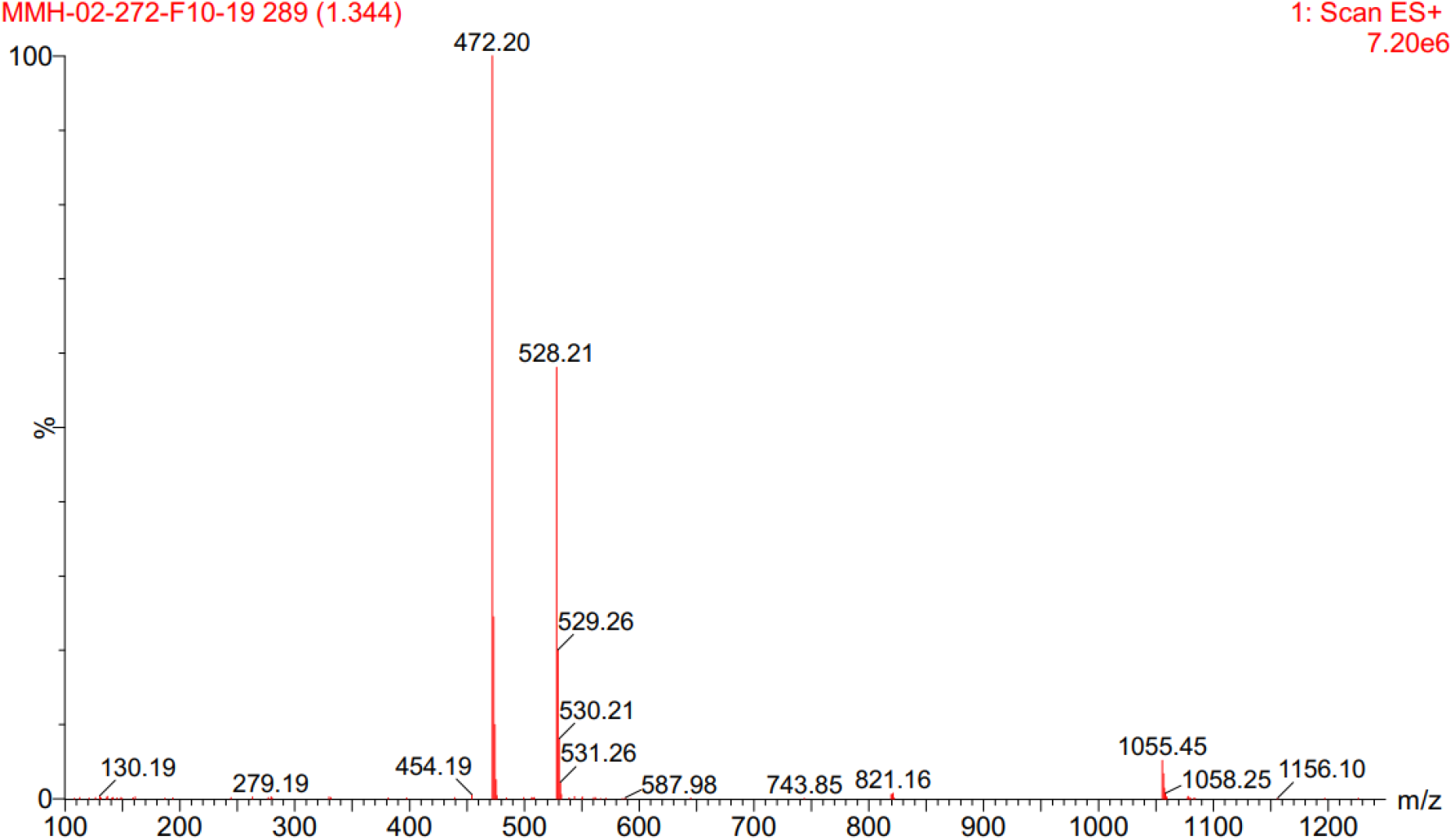

### MMH2-NR (500 MHz ^1^H NMR in DMSO-*d*_6_)

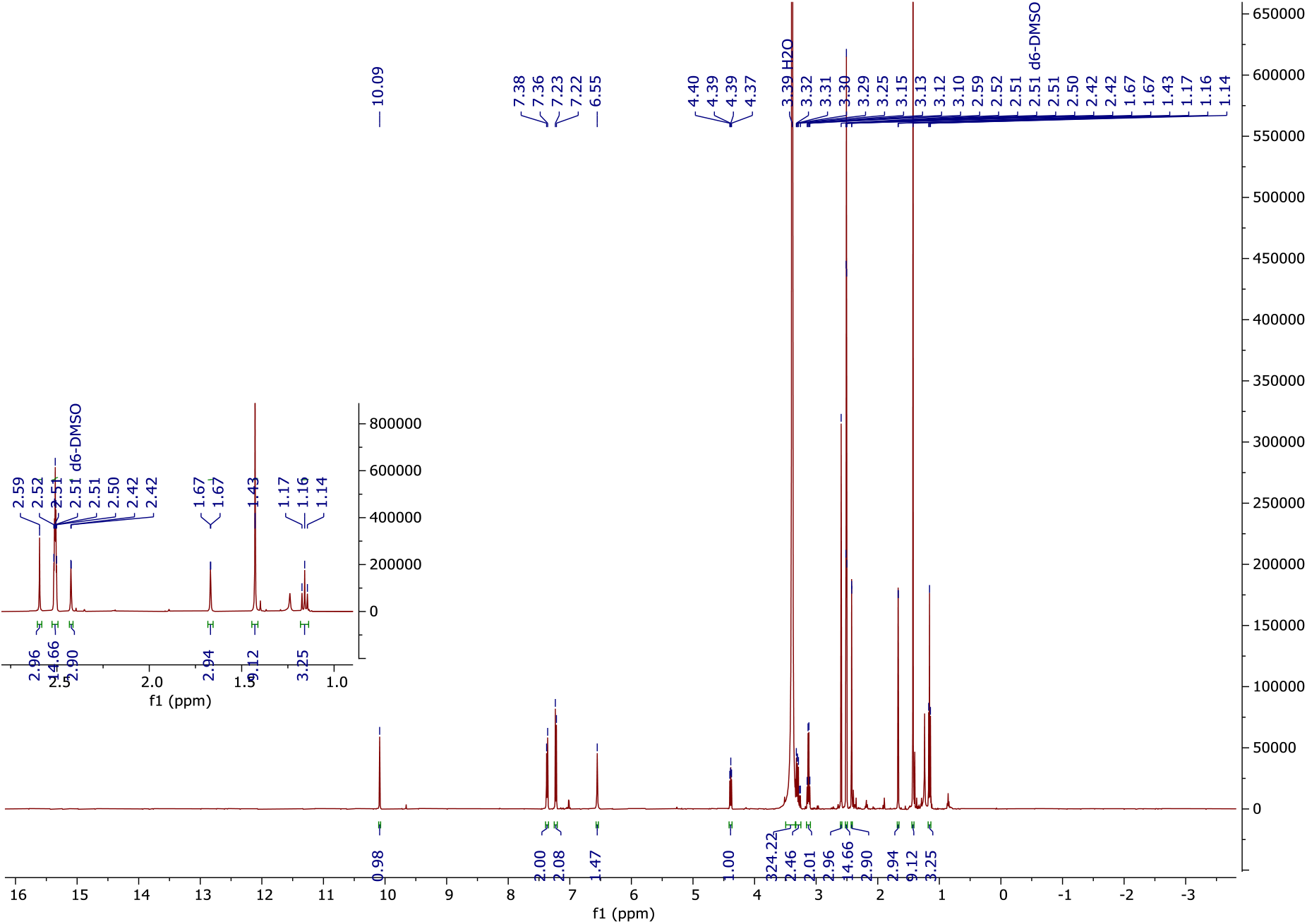

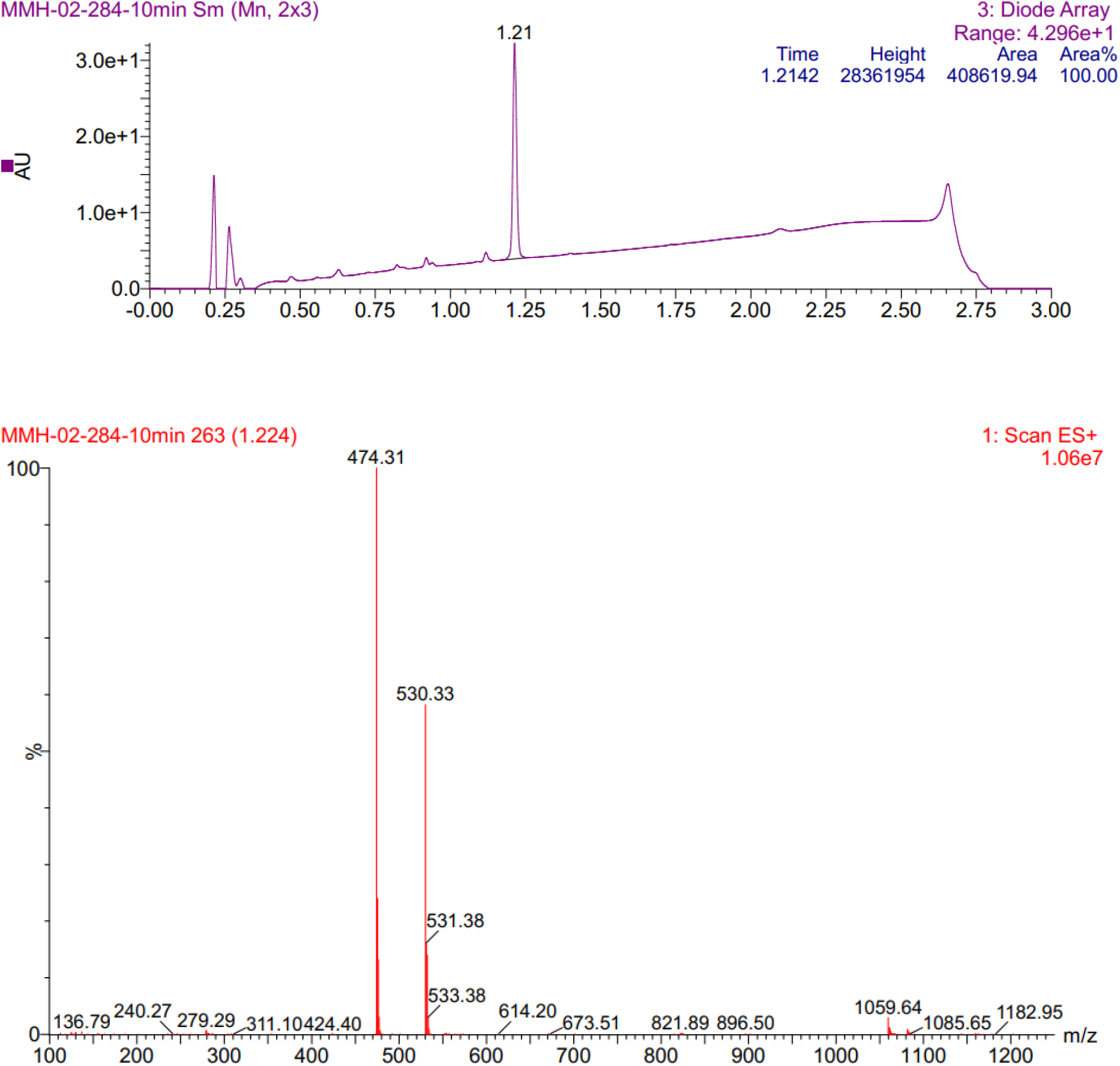

### MMH248 (500 MHz ^1^H NMR in DMSO-*d*_6_)

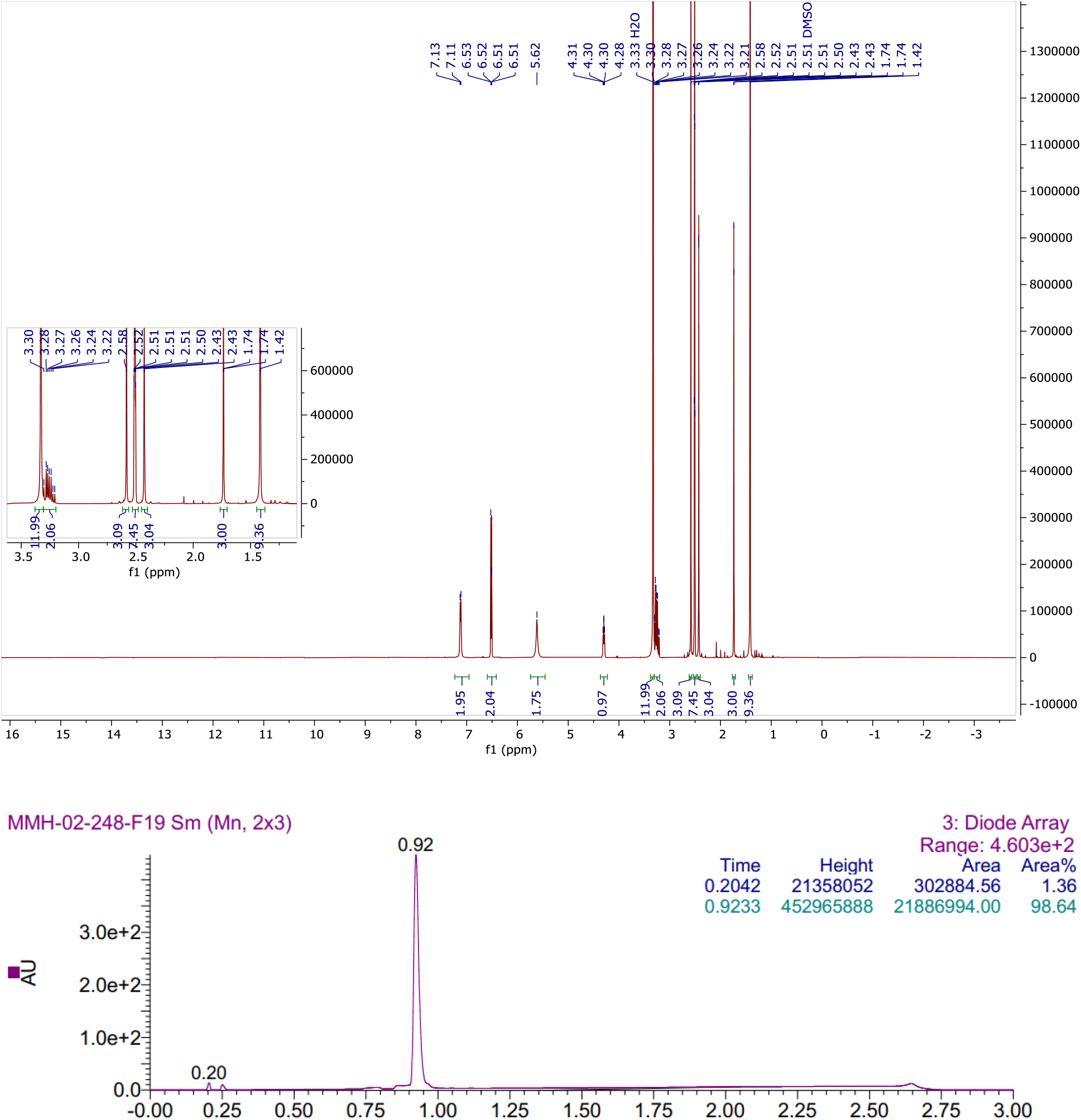

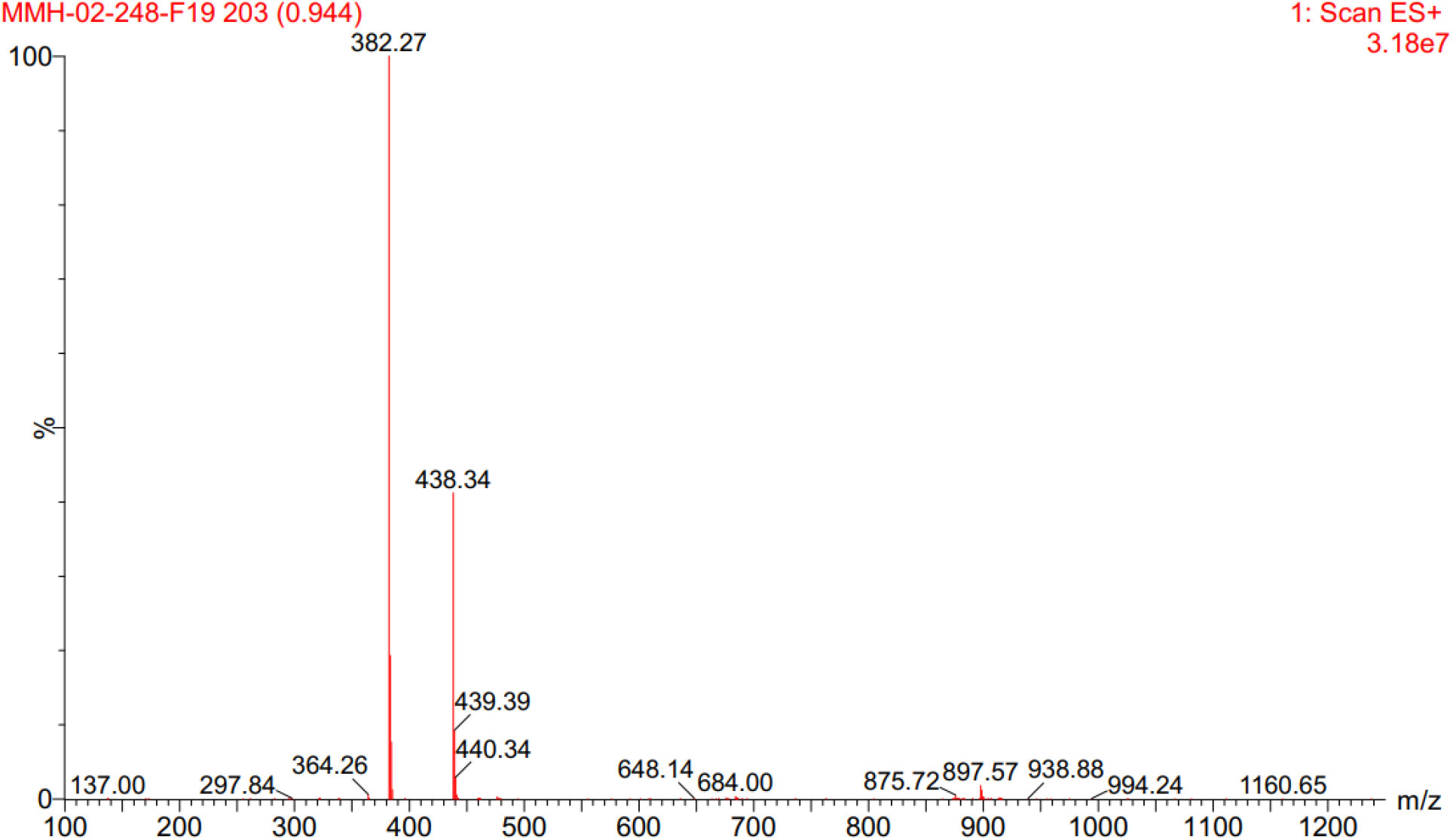

### MMH249 (500 MHz ^1^H NMR in DMSO-*d*_6_)

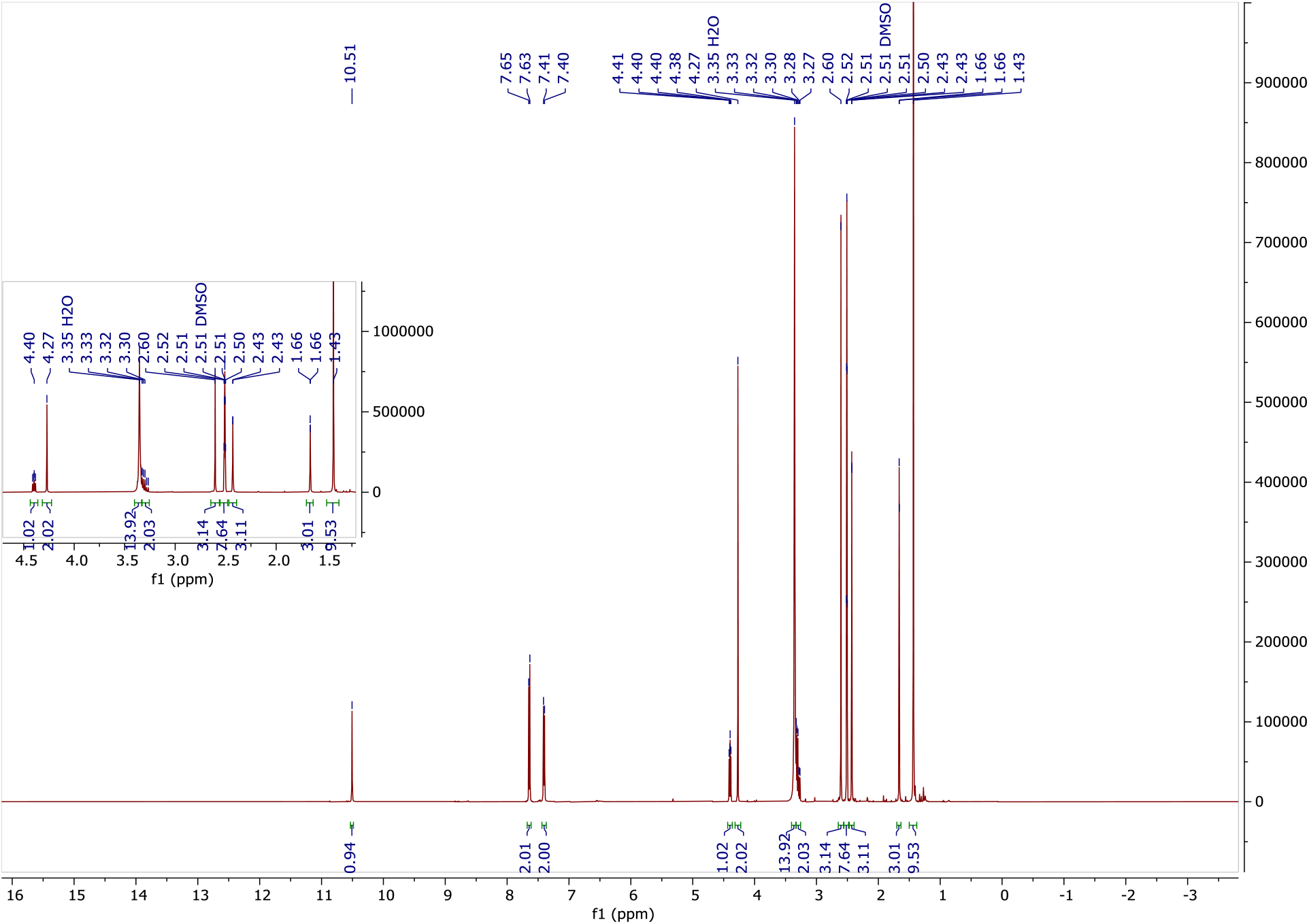

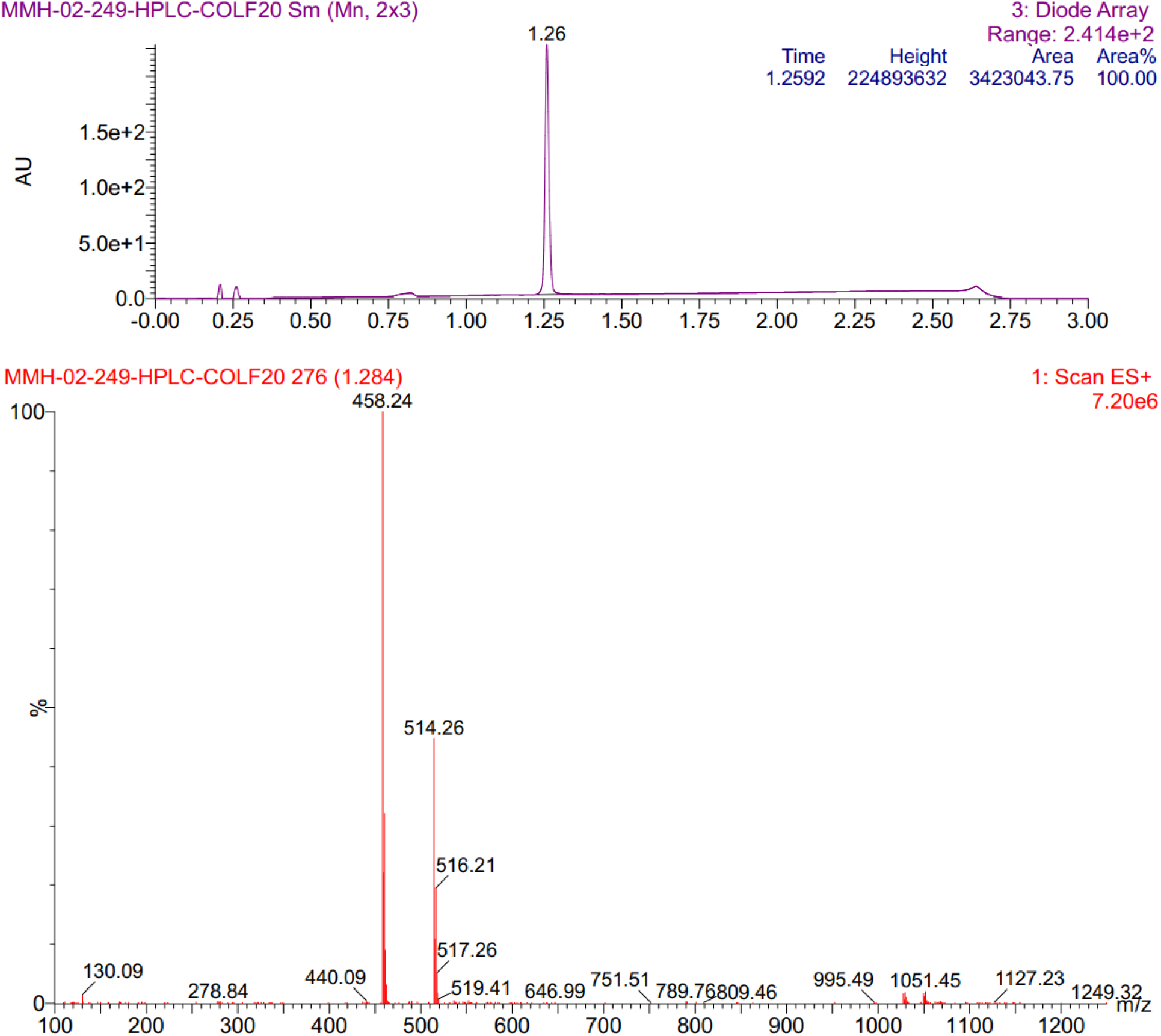

### MMH263 (500 MHz ^1^H NMR in DMSO-*d*_6_)

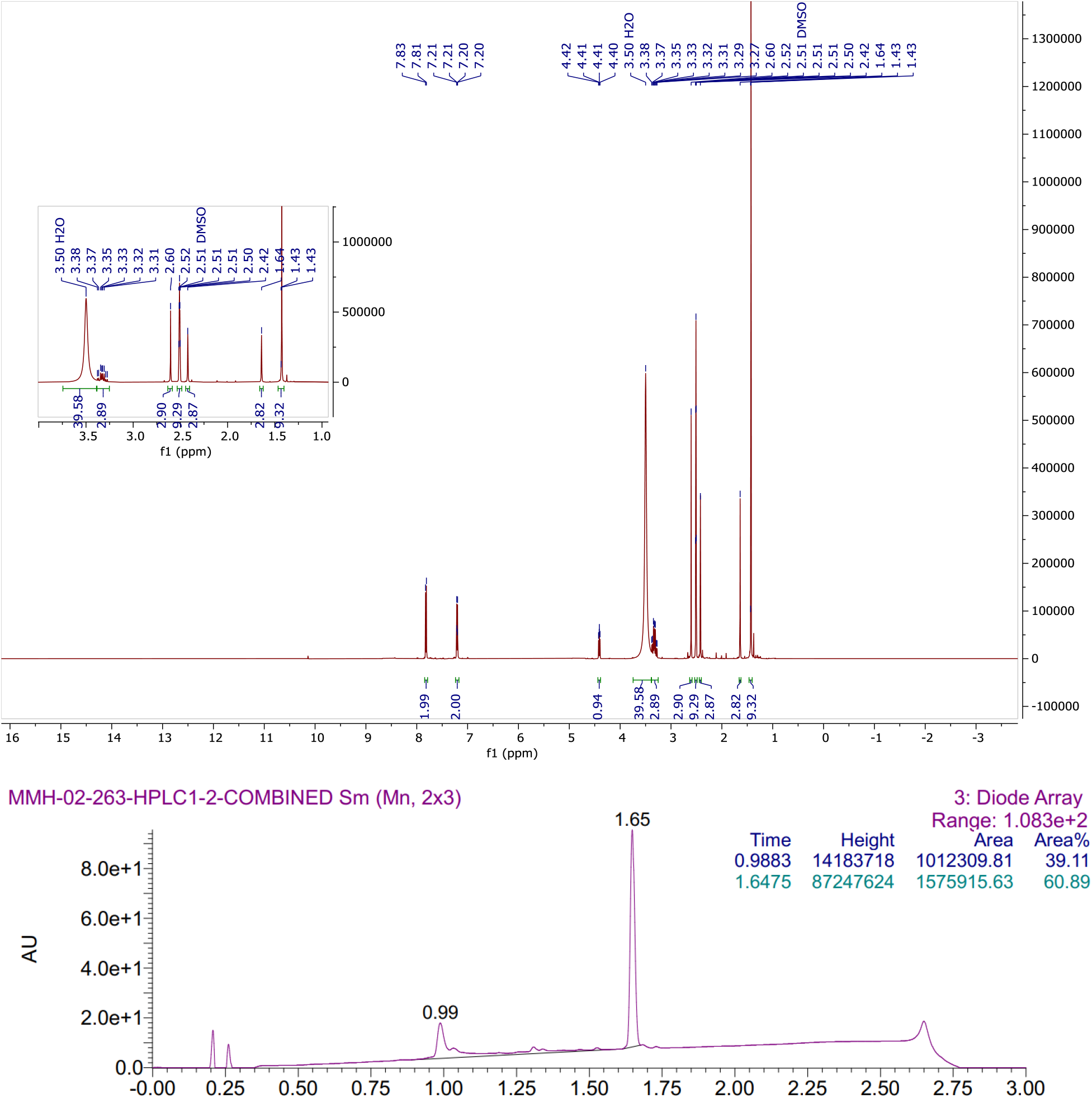

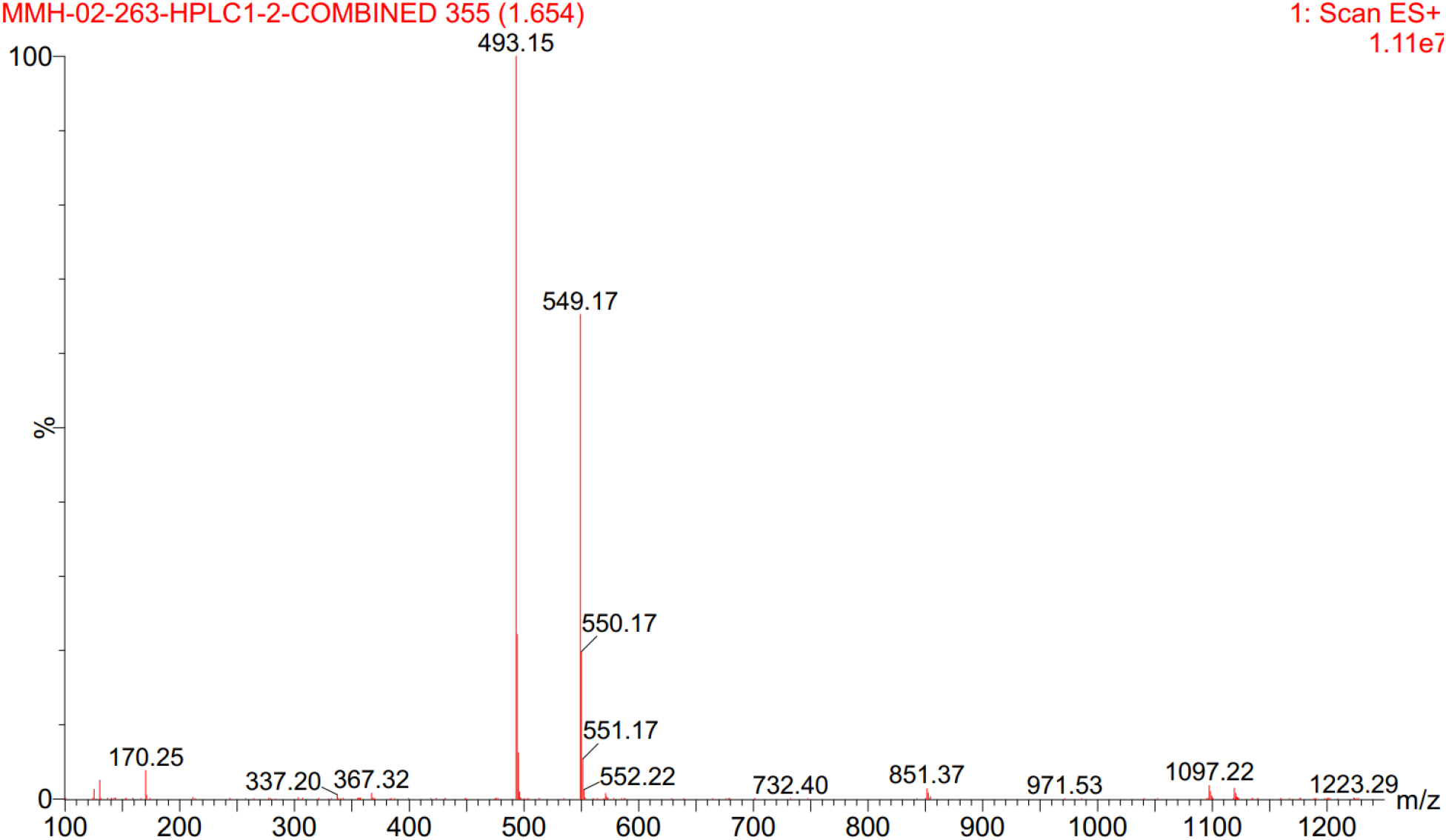

### MMH269 (500 MHz ^1^H NMR in DMSO-*d*_6_)

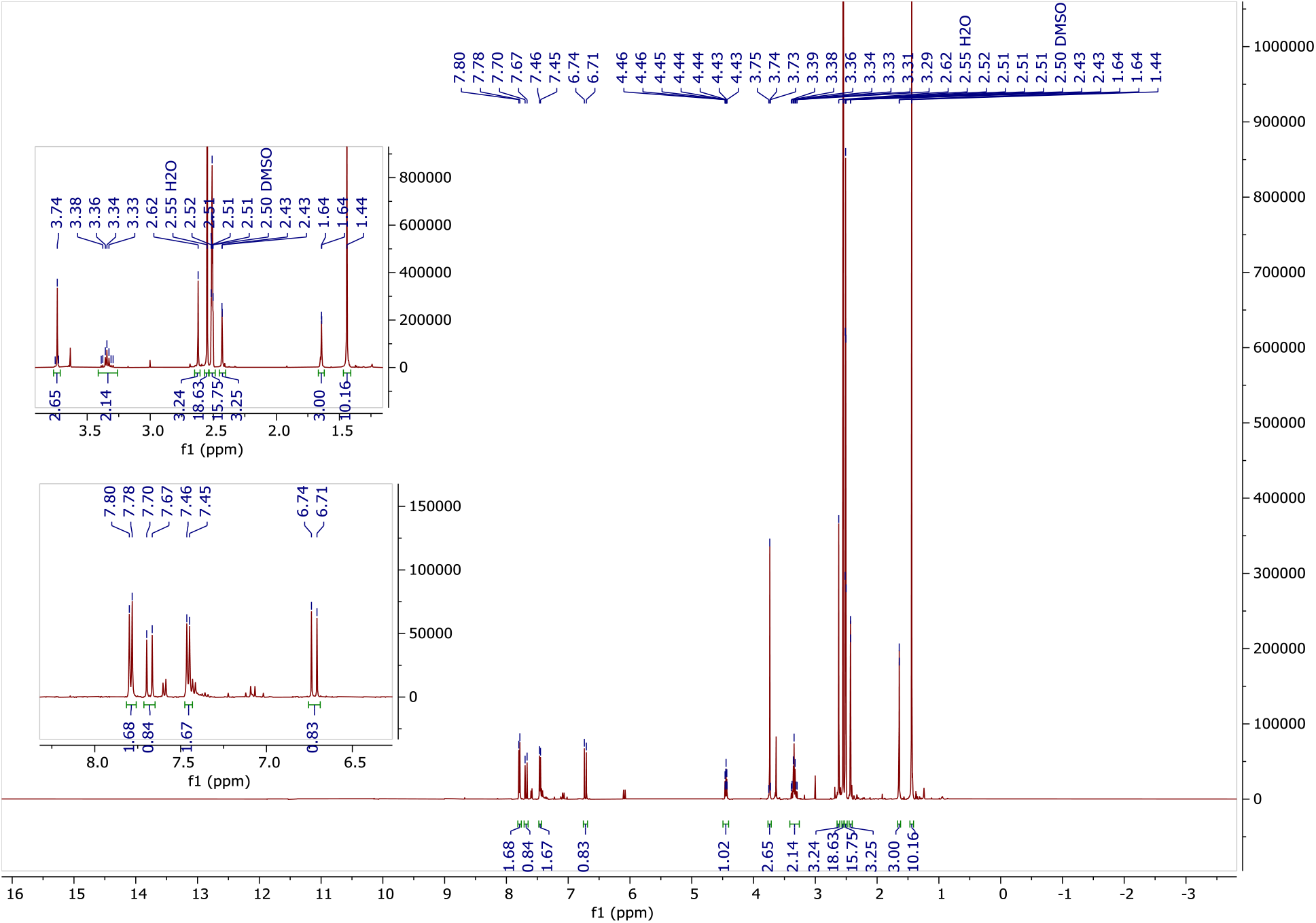

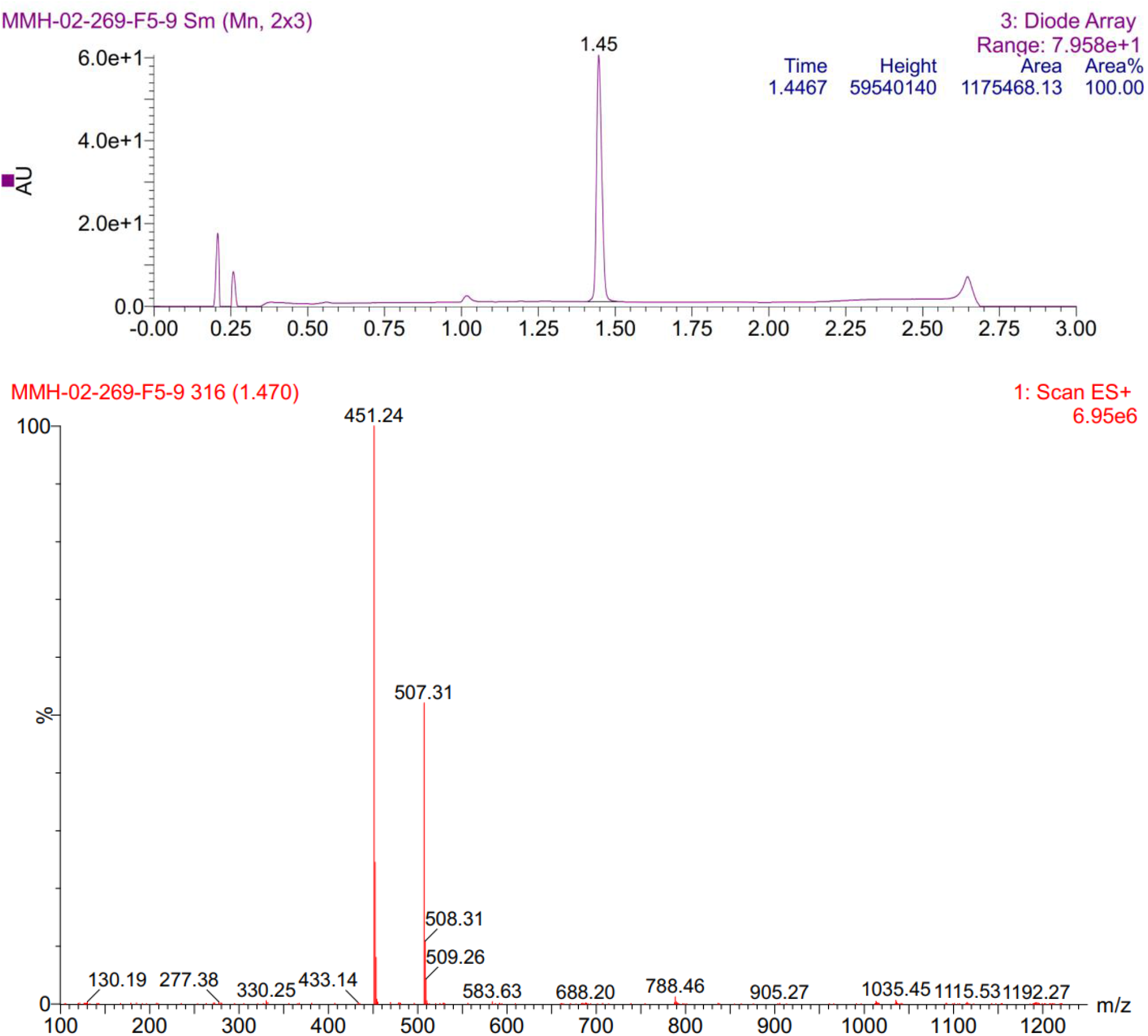

### MMH271 (500 MHz ^1^H NMR in DMSO-*d*_6_)

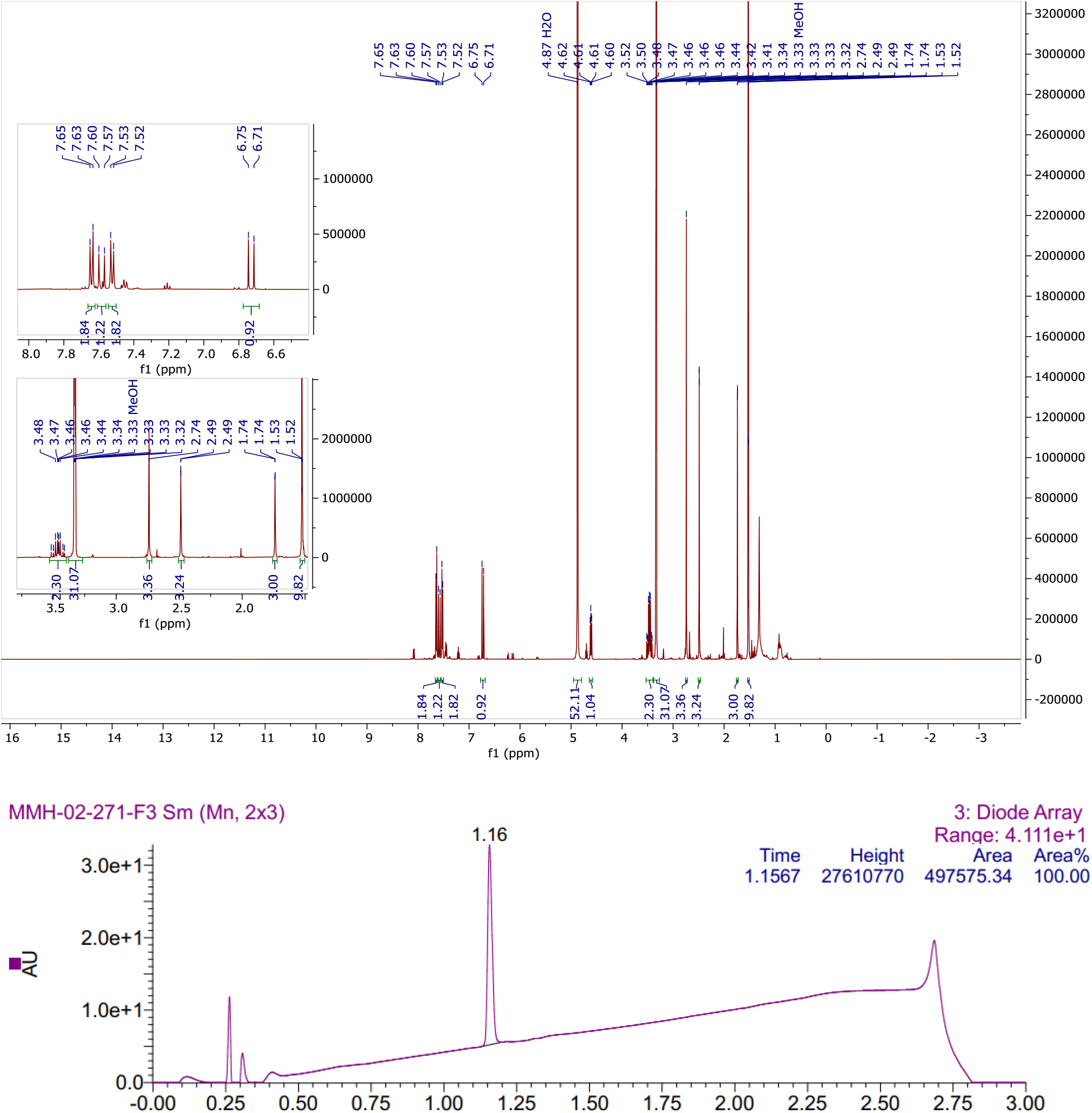

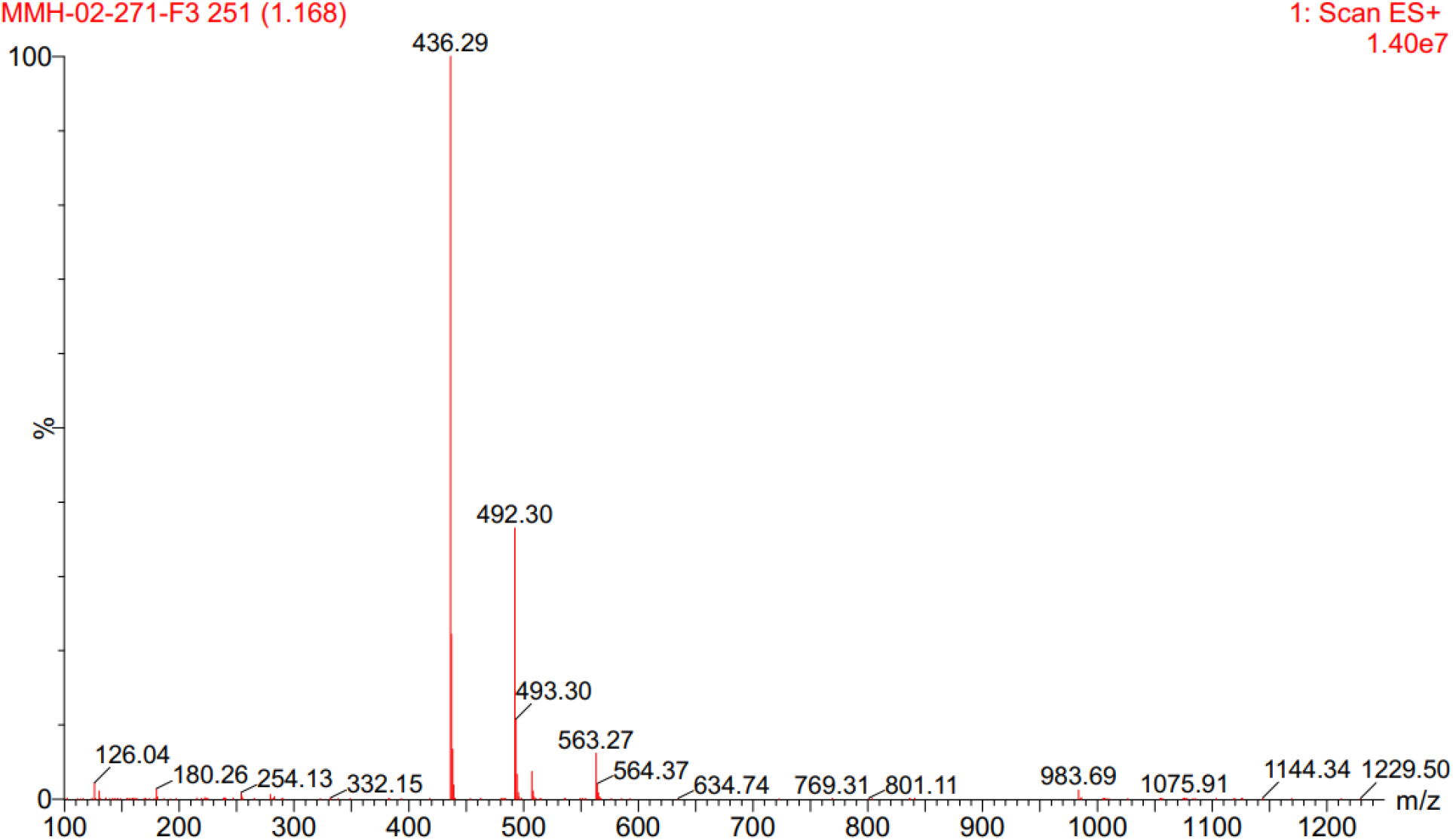

### MMH287 (500 MHz ^1^H NMR in DMSO-*d*_6_)

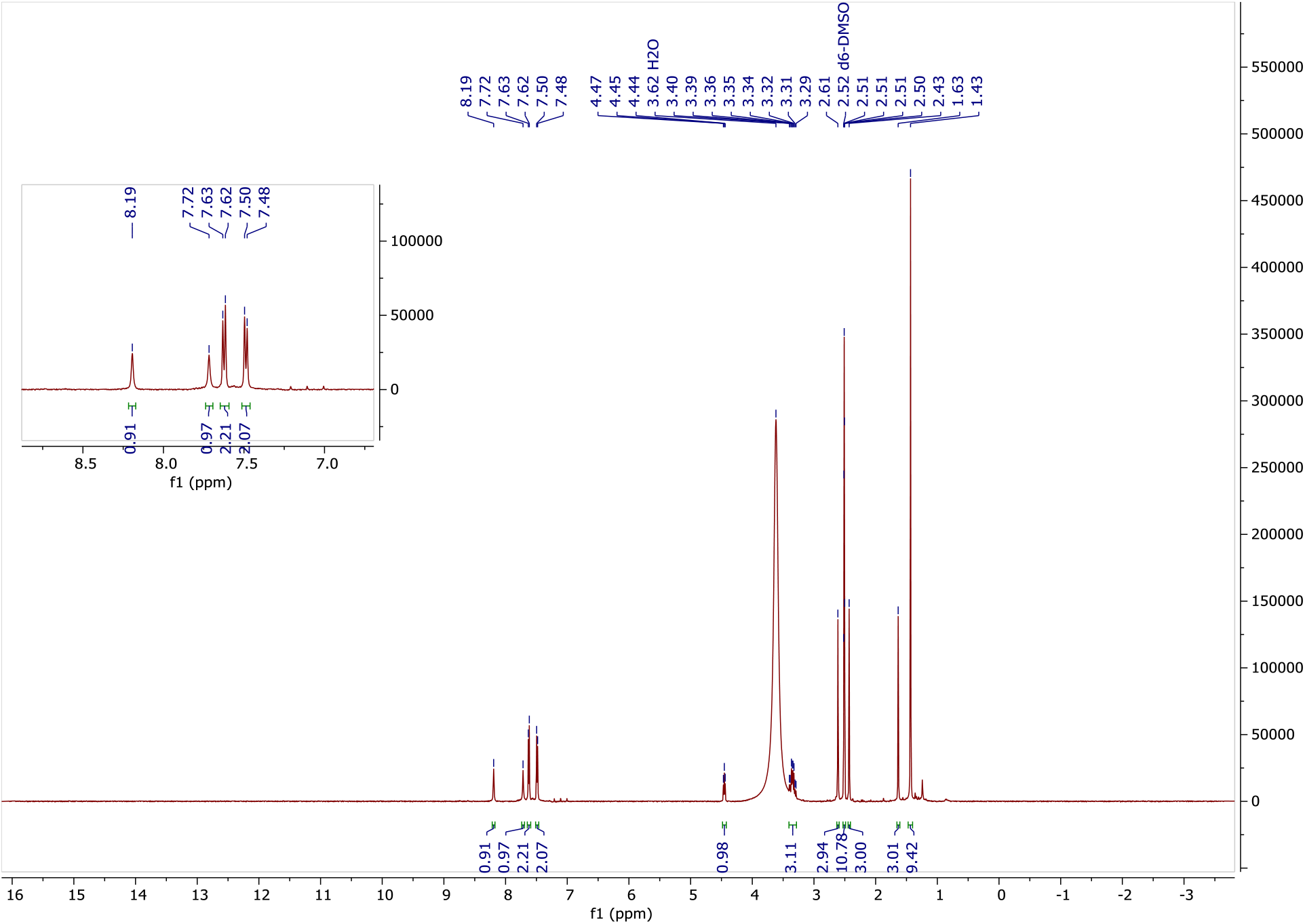

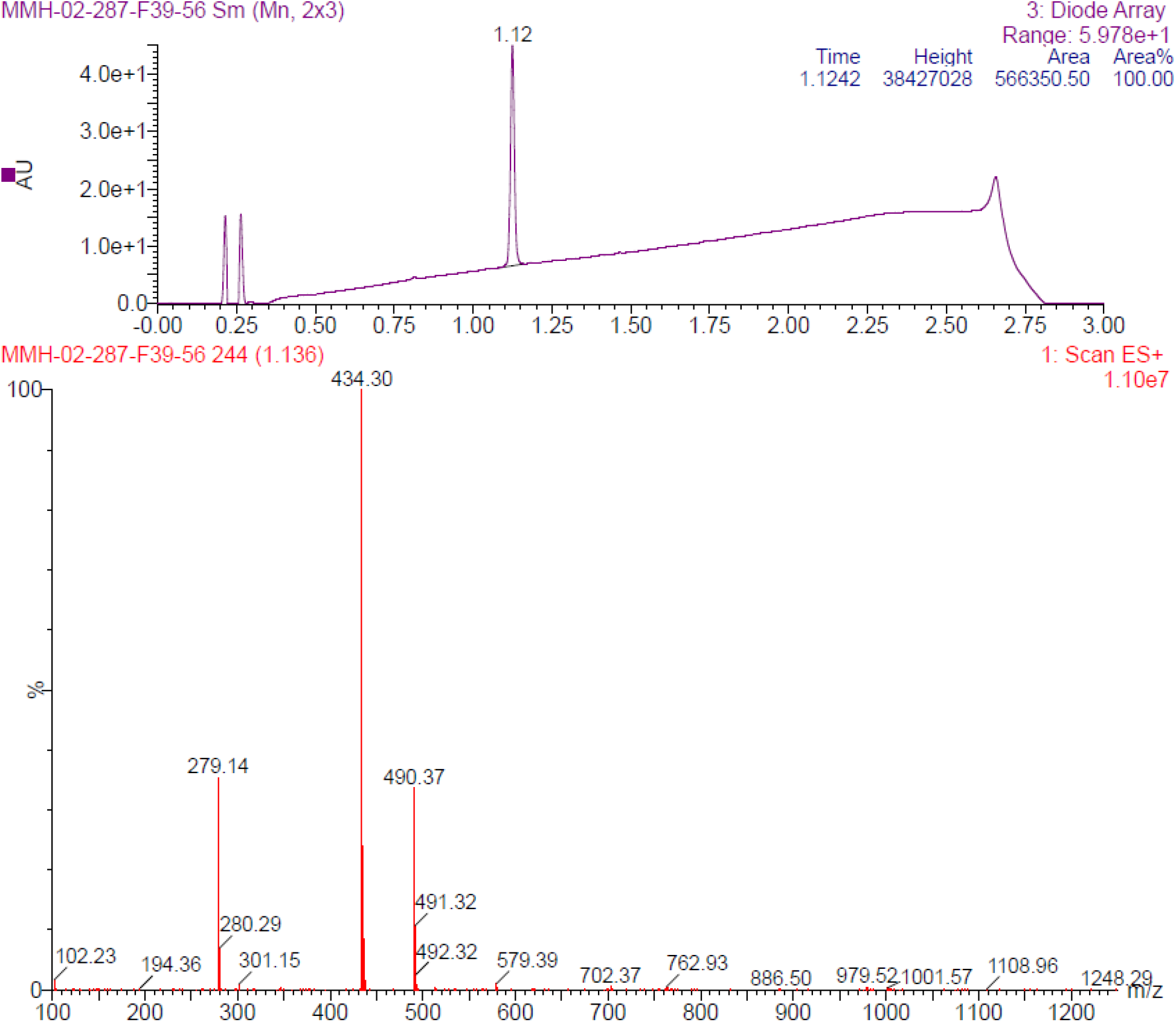

### TMX458 (500 MHz ^1^H NMR in DMSO-*d*_6_)

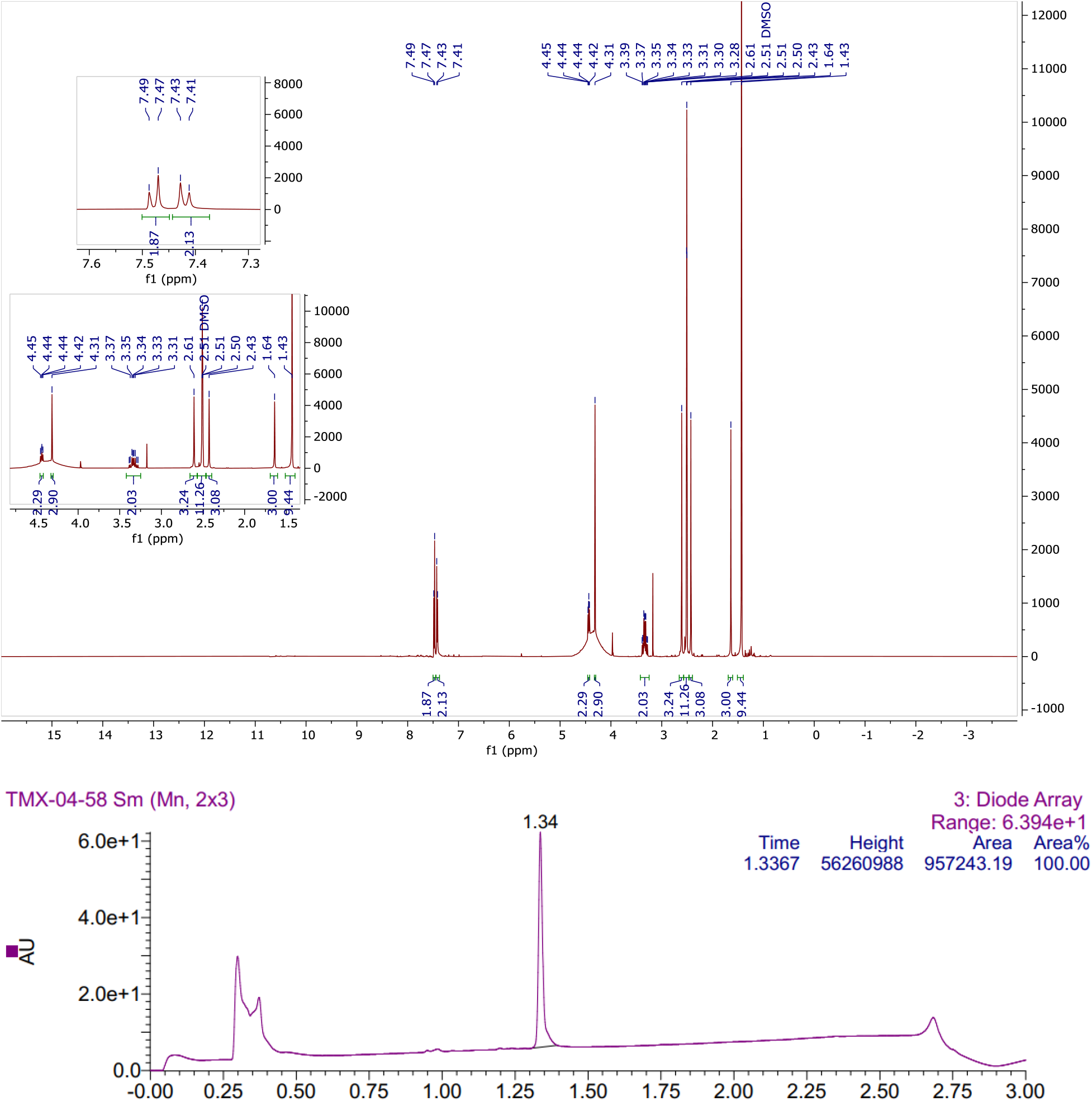

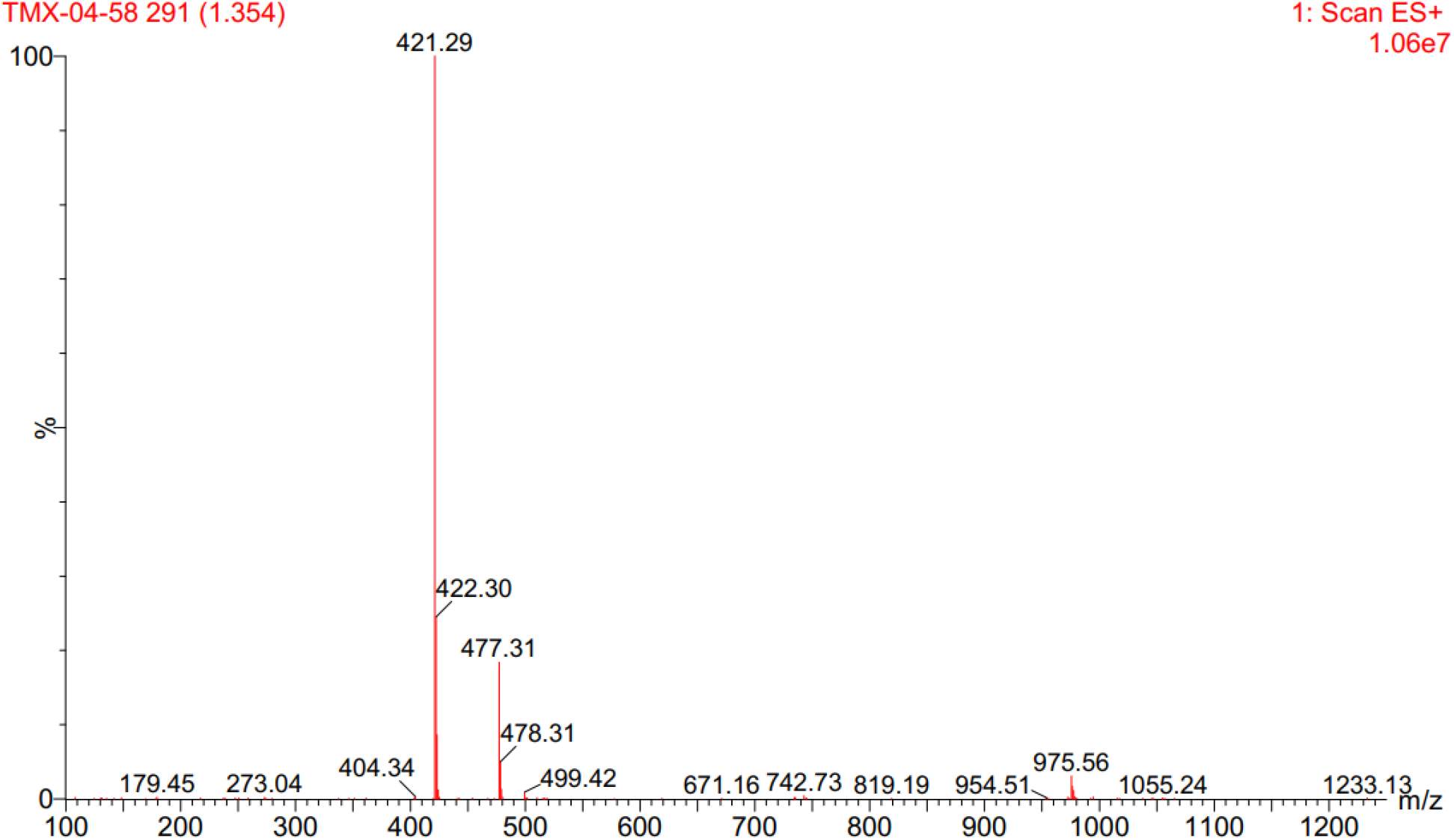

### TMX4128 (500 MHz ^1^H NMR in DMSO-*d*_6_)

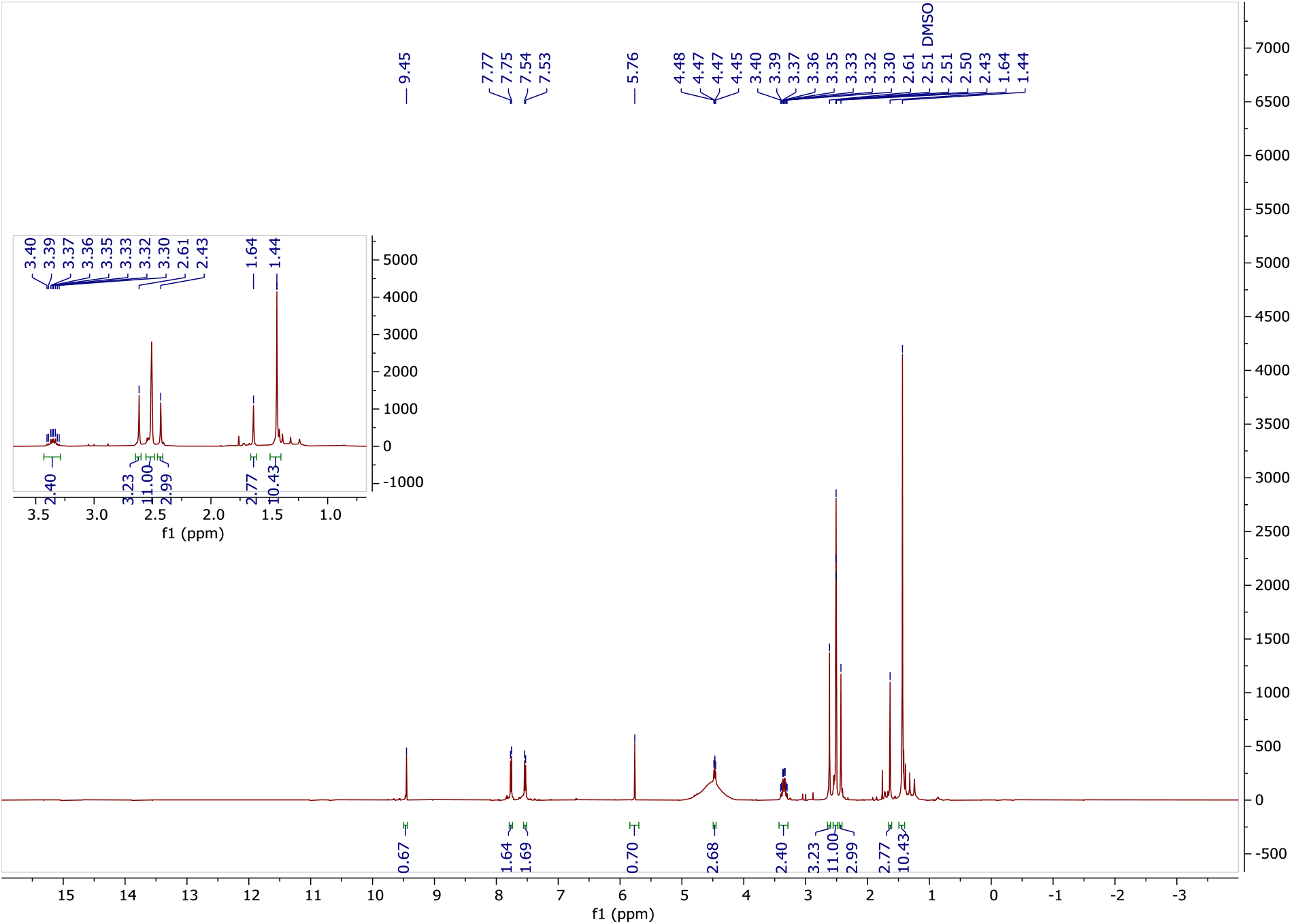

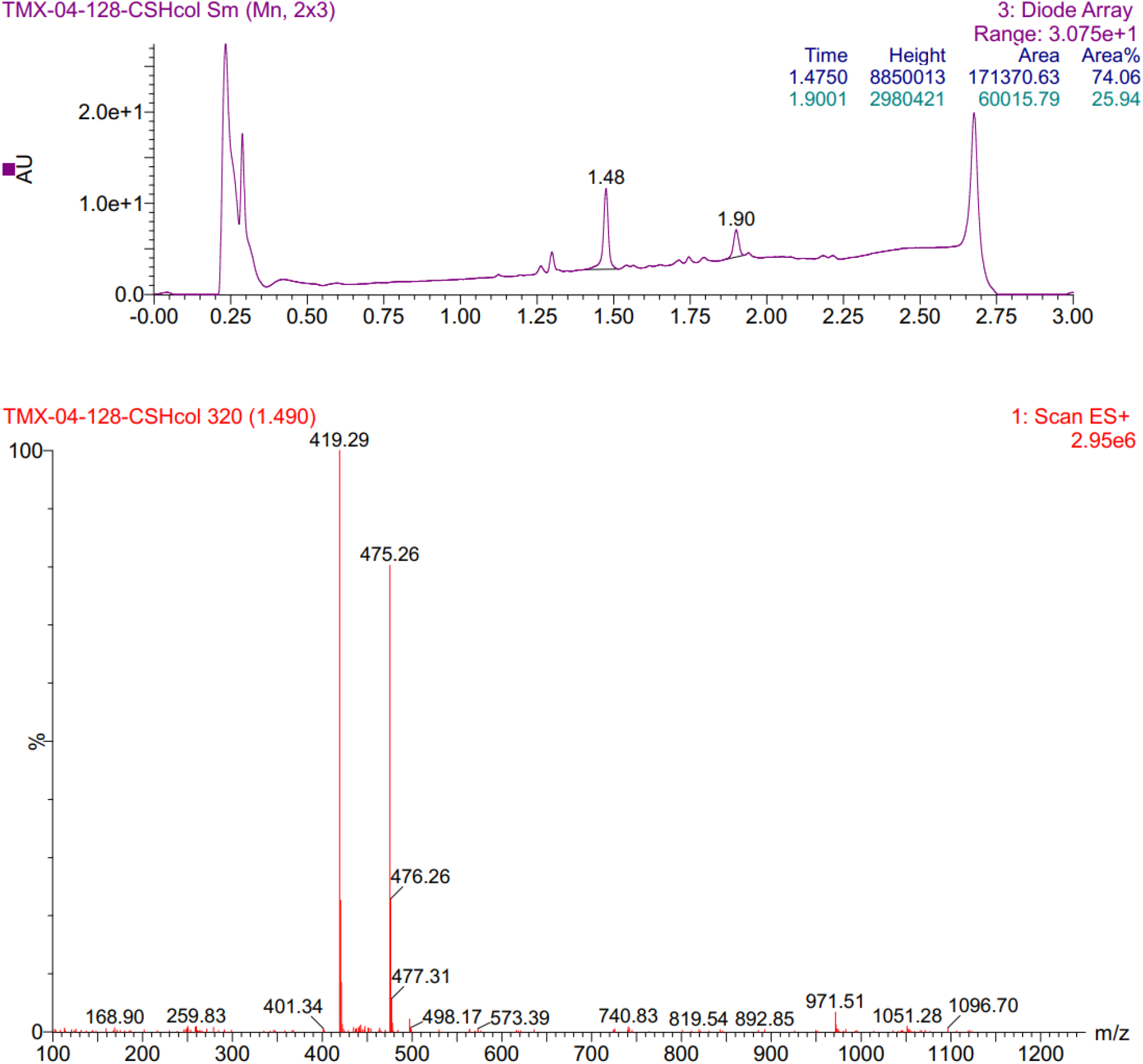

